# Remote Control of Cell Signaling through Caveolae Mechanics

**DOI:** 10.1101/2025.07.27.666936

**Authors:** Satish Kailasam Mani, Nicolas Tardif, Olivier Rossier, Ismail M. Khater, Xuesi Zhou, Victor Breton, Filipe Nunes Vicente, AV Radhakrishnan, Céline Gracia, Pamela Gonzalez Troncoso, Isabel Brito, Richard Ruez, Melissa Dewulf, Ghassan Hamarneh, Ivan Robert Nabi, Philippe Cuniasse, Pierre Sens, Grégory Giannone, Cédric M. Blouin, Christophe Lamaze

## Abstract

Caveolae are invaginated plasma membrane nanodomains traditionally associated with membrane trafficking and signaling. These multifunctional organelles are also essential mechanosensors mediating the cell response to mechanical stress. We investigated the role of caveolae mechanics in regulating various signaling pathways. Single molecule imaging and super resolution microscopy revealed that mechanical stress rapidly triggers caveolae disassembly and the release of caveolin-1 scaffolds, which exhibit enhanced diffusion at the plasma membrane. This promoted direct interaction between the caveolin-1 scaffolding domain and the tyrosine kinase JAK1, leading to the inhibition of its catalytic activity. A similar process was observed for eNOS, PTEN, and PTP1B. Remote control of signaling by caveolae was validated by a theoretical model based on caveolae thermodynamics. These findings establish a novel mechanotransduction paradigm where signaling information is decoded remotely from the initial mechanosensing caveola, through dynamic and reversible assembly of tension-controlled complexes between signaling effectors and caveolin-1 scaffolds.

## INTRODUCTION

Caveolae are discrete bulb shaped structures located at the plasma membrane with diameters ranging between 50-80 nm in diameter and that are occasionally clustered in ‘rosettes’ ^1,2^. They were first identified through electron microscopy (EM) in epithelial and endothelial cells over 70 years ago ^3,4^. The integral membrane protein caveolin-1 oligomerizes to form the primary 8S building blocks of caveolae assembly in many cell types, particularly abundant in adipocytes, endothelial cells, and muscle cells. The caveolin family of proteins consist of three isoforms: caveolin-1 (Cav1), caveolin-2 (Cav2), and muscle-specific caveolin-3 (Cav3) ^5–7^. Despite the presence of electron dense regions on their cytoplasmic face observed in early EM experiments, caveolae were traditionally described as non-coated invaginations, in contrast to the characteristic fuzzy coat seen on clathrin-coated pits. The composition and organization of the caveolar coat were ultimately elucidated with the discovery of the cavin family of cytosolic proteins, which include cavin1, cavin2, cavin3, and cavin4. Cavin1, is essential for caveolae morphogenesis in all cell types ^8^ while cavin4 expression is limited to muscle cells ^9–11^. Several accessory proteins including EHD2, pacsin2/syndapin2 (pacsin3/syndapin3 in muscle cells), and filamin A have also been localized at the neck of caveolae and proposed to control caveolae stability and dynamics through interactions with the actin cytoskeleton ^12–16^. In a recent cryo-EM study, the quaternary structure of the Cav1 complex was resolved ^17^. The 8S complex has been proposed to consist of 11 primary α-helical protomers that are tightly packed into a 14 nm spiral disk. The N-termini are located on the outer ring, while the C-termini form a central cylindrical β-barrel structure.

Multiple studies have investigated the role of caveolae and/or Cav1 in various vital biological processes, including transcytosis and endocytosis, lipid homeostasis, and signal transduction ^18–21^. The diverse range of functions attributed to caveolae and their associated proteins accounts for their involvement in several human diseases ^22^. For example, mutations or impaired expression of caveolins and cavins have been associated with lipodystrophy, vascular dysfunction, cancer, and muscle dystrophies ^23-^ 26.

The most explored function of Cav1 concerns the regulation of intracellular signal transduction ^1,27,28^. Earlier studies have proposed that caveolae and/or Cav1 can modulate the activity of various growth factors, signaling receptors, and kinases. These include endothelial nitric oxide synthase (eNOS), insulin receptor, epidermal growth factor receptor (EGFR), Src-like kinases, HRas and KRas ^29–33^. Difficulties to unambiguously localize these signaling effectors into caveolae has however questioned the reliability of these studies which were primarily based on overexpression of signaling proteins and Cav1, possibly leading to localization artefacts. Consequently, caveolae were found to exclude bulk plasma membrane protein with transmembrane and cytosolic domains, and instead to concentrate membrane lipids ^34^. In this context, a role for Cav1 in controlling signaling outside of caveolae was proposed based on the ability of endogenous Cav1 to inhibit EGFR signaling in tumor cells that lack caveolae ^35^. Subsequent studies have shown that Cav1 controls cancer cell migration and tumor progression by regulating focal adhesion signaling and tension in prostate cancer PC3 cells that lack cavin1 and therefore caveolae ^9,36–38^. These non-caveolar Cav1 assemblies were termed scaffolds ^39^ and were recently visualized at the plasma membrane by single molecule localization super-resolution microscopy ^40,41^ and by EM ^42^. The question of how the central function of Cav1 in caveolae morphogenesis relates to its control over the activity of signaling receptors that are not located within caveolae is unresolved, making it an essential area of research.

In 2011, we uncovered a novel function of caveolae in mechano-sensing. Upon exposure to mechanical stress, caveolar invaginations undergo rapid flattening within the plasma membrane to expand membrane area, effectively acting as a buffer against sudden increases in membrane tension, thereby protecting the plasma membrane against rupture ^43^. This essential role of caveolae in cell mechanics was further confirmed in several cell types *in vitro* and *in vivo* ^42,44–48^. The new role of caveolae in cell mechanics has prompted a re-evaluation of their conventional functions, and there is now a consensus that caveolae need to be revisited in light of this new understanding^2,49–52^.

Here, we investigated the effects of the tension-dependent cycle of caveolae disassembly and reassembly on intracellular signaling. Using super resolution microscopy and live-cell single molecule imaging, we found that in response to mechanical stress, caveolae rapidly disassembled into smaller Cav1 assemblies corresponding to scaffolds ^40,41^. The pool of released Cav1 scaffolds was found to diffuse rapidly in the plasma membrane and to directly interact through their scaffolding domain with several signaling effectors outside of caveolae, including the Janus kinase JAK1. As a result, JAK1 catalytic activity was inhibited as shown by the lack of STAT3 activation by interferon-α (IFN-α). Upon stress release, JAK-STAT activation by IFN-α resumed to normal levels. AlphaFold3 predictions combined with molecular dynamics simulations yielded a structural model of the Cav1–JAK1 kinase domain interaction that is consistent with our Cav1 mutagenesis data and identifies candidate JAK1 interface residues. We could extend the mechanical regulation of signaling by Cav1 scaffolds to PTEN and PTP1B tyrosine phosphatases, and to eNOS. A theoretical model based on the thermodynamics of caveolae assembly and disassembly under mechanical stress could recapitulate these observations and further establish the generality of the proposed remote signaling mechanism. Altogether, our study provides a new paradigm in mechanotransduction by which selective signaling pathways are remotely controlled at locations distant from budded caveolae.

## RESULTS

### The JAK/STAT signaling pathway is controlled by caveolae under mechanical stress

Our primary objective was to investigate whether the mechano-dependent cycle of caveolae disassembly and reassembly could serve as a mechanical switch, potentially enabling caveolae and /or caveolins to modulate specific signaling pathways ^1,49^. Our previous studies suggesting that caveolae mechanics could regulate JAK/STAT signaling in muscle cells ^53^ prompted us to further explore this regulatory mechanism in mouse lung endothelial cells (MLEC), which are particularly enriched in caveolae. We analyzed the activation of the JAK/STAT signaling pathway induced by IFN-α in wild-type (WT) and Cav1 knockout (Cav1^-/-^) MLEC cells subjected to 25% uniaxial stretching. We initially investigated the level of STAT3 phosphorylation (pSTAT3) at tyrosine 705 upon IFN-α stimulation. Western blot analysis in MLEC WT and MLEC Cav1^-/-^ cells revealed a 52% decrease in pSTAT3 levels in stretched cells as compared to non-stretched or resting cells (Fig. 1a). No activation of STAT3 was measurable in the absence of IFN-α stimulation in both cell types (Extended Data Fig. 1a). The level of STAT3 tyrosine phosphorylation was unaltered in MLEC Cav1^-/-^ cells that were subjected to stretch, further confirming that this regulatory mechanism is dependent on the presence of functional caveolae and/or Cav1 (Fig. 1a). Consequently, the reduction in STAT3 phosphorylation levels resulted in a corresponding inhibition of STAT3 nuclear translocation in stretched cells (Fig. 1b,d). Similarly, the level of pSTAT3 in the nucleus was not affected by stretch in MLEC Cav1^-/-^ cells (Fig. 1c,d). Under these conditions, we found that STAT1 nuclear translocation remained unaffected (Fig. 1b-d). These results strongly suggest the involvement of caveolae and/or Cav1 in the specific modulation of STAT3 phosphorylation levels in response to mechanical stress.

**Figure 1:**
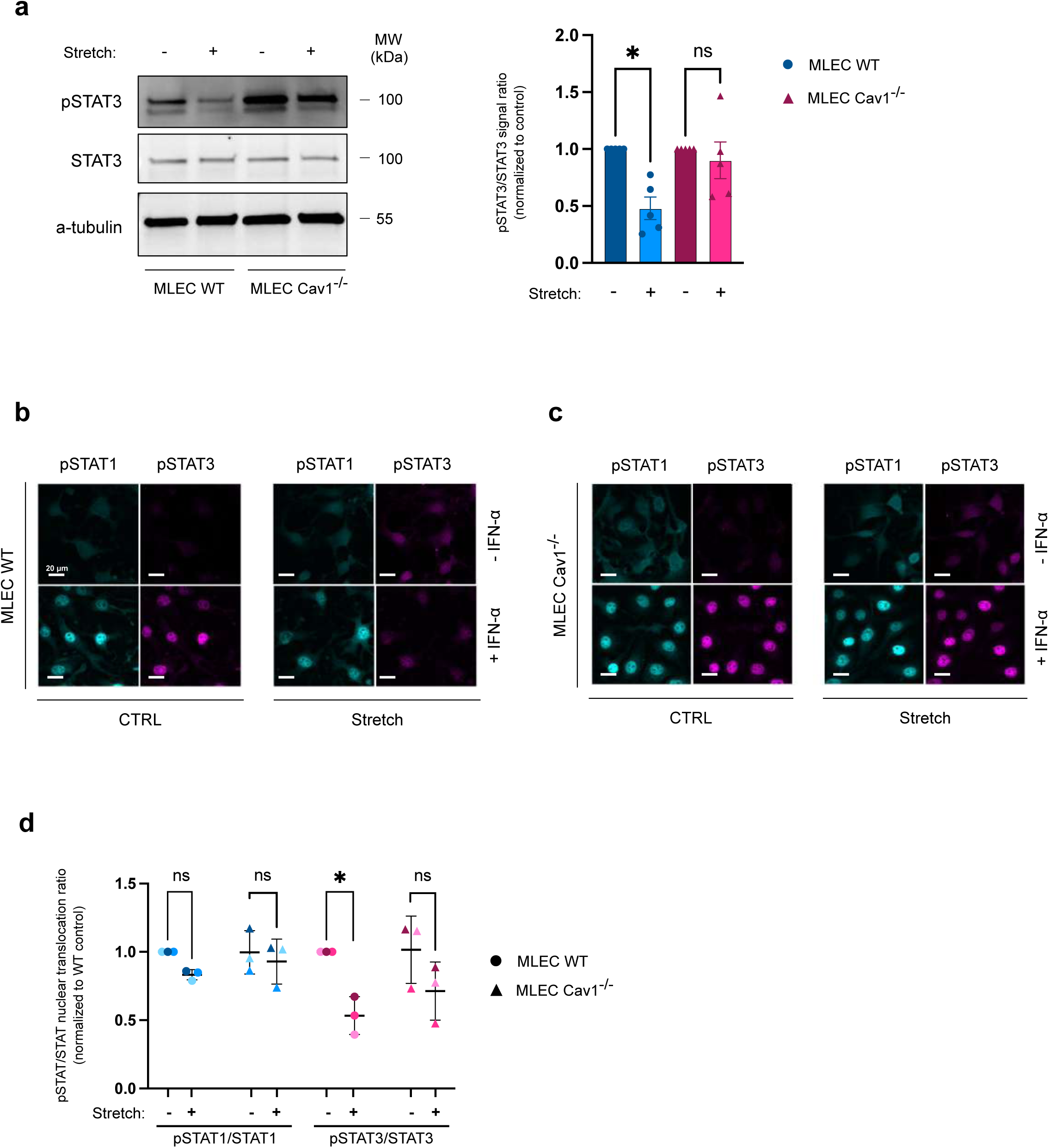
Caveolae mechanics modulate the JAK-STAT signaling pathway. **(a)** STAT3 phosphorylation levels induced by IFN-α stimulation in MLEC WT and MLEC Cav1^-/-^ cells subjected to uniaxial stretch or not. Representative immunoblots and quantification of signal ratio relative to control condition for N=5 independent experiments; mean values ± SEM. **(b-d)** Immunofluorescence images of nuclear translocation of pSTAT1 and pSTAT3 in MLEC WT **(b)** and MLEC Cav1^-/-^ **(c)** cells subjected to uniaxial stretch or not. Representative data for N=3 independent experiments. Statistical analysis **(b)** one-way ANOVA with Friedman test, **(d)** two-way ANOVA with Šídák test. *P<0.05 and ns: not significant.

### Interaction of Cav1 with JAK1 results in STAT3 inhibition

The activation of JAK-STAT signaling by IFN-α depends on the ubiquitous IFNAR receptor, which consists of two receptor subunits, IFNAR1 and IFNAR2 ^54^. STAT3 is a direct cytosolic downstream effector of TYK2 and JAK1 tyrosine kinases, which are associated with IFNAR1 and IFNAR2, respectively ^55^. We hypothesized that the modulation of STAT3 tyrosine phosphorylation by caveolae/Cav1 in response to mechanical stress could be mediated through the interaction of Cav1 with JAK1 or TYK2. Treatment of cells with a hypo-osmotic medium (30 mOsm) for 5 minutes results in increased membrane tension due to cell swelling and leads to rapid caveolae disassembly, similar to the response observed during cell stretching ^43^. Co-immunoprecipitation (co-IP) of both endogenous Cav1 and JAK1 was performed on MLEC WT and MLEC Cav1^-/-^ cells subjected to hypo-osmotic shock. We found that Cav1 interacts with JAK1 in resting cells, and this interaction is significantly increased by up to 3-fold in response to mechanical stress (Fig. 2a). This effect was rapid, with maximal interaction between JAK1 and Cav1 observed after 5 min of hypo-osmotic shock. The interaction increased with the intensity of the shock and did not further increase with longer exposure times (Fig. 2b,c). We repeated the co-IP experiment with IFN-α stimulation to assess the effect of increased Cav1-JAK1 interaction on STAT3 phosphorylation. The increase in interaction between Cav1 and JAK1, induced by hypo-osmotic shock, was correlated with a substantial reduction of up to 62% in STAT3 phosphorylation level (Fig. 2d). Remarkably, upon returning to iso-osmotic conditions when caveolae had reassembled to their initial numbers during the recovery phase ^43^, we observed that both the levels of Cav1 interaction with JAK1 and IFN-α-induced STAT3 phosphorylation reverted to the levels measured in the resting state. We also measured a higher level of STAT3 phosphorylation in MLEC Cav1^-/-^ cells compared to WT cells, in agreement with the known regulation of STAT3 activity by Cav1 ^56,57^. These data suggest that the amount of Cav1 released from caveolae during mechanical disassembly can control the degree of Cav1-JAK1 interaction, which in turn can regulate STAT3 phosphorylation. This process is reversible and governed by mechanical stress.

**Figure 2:**
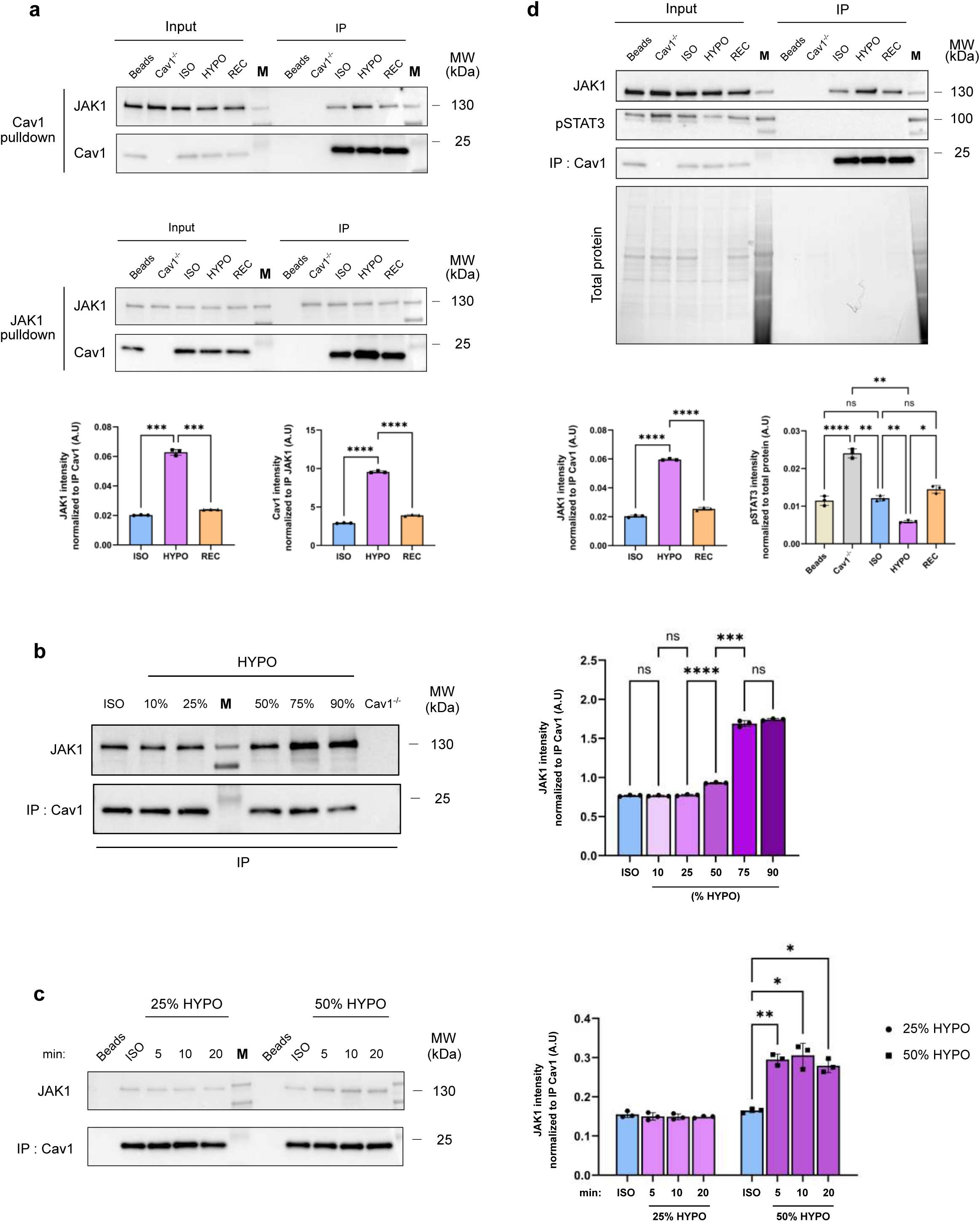
Cav1 dependent inhibition of STAT3 activation is mediated through JAK1 interaction. **(a)** Co-immunoprecipitation of endogenous Cav1 and JAK1 in iso-osmotic (ISO), hypo-osmotic (HYPO) and successive hypo-osmotic and iso-osmotic treatment (REC) in MLEC WT cells. Quantification is based on the signal intensity ratio (JAK1 and Cav1) relative to the intensity of the corresponding immuno-precipitated protein (Cav1 and JAK1 respectively). **(b)** Representative blots of co-immunoprecipitation of endogenous JAK1 with Cav1 in MLEC WT cells under ISO condition and increasing percentages of hypo-osmotic (HYPO) shock. Quantification is based on the signal intensity ratio relative to the corresponding immunoprecipitated Cav1 protein levels. **(c)** Representative blots of co-immunoprecipitation of endogenous JAK1 with Cav1 in MLEC WT cells under ISO condition and increasing exposure times to 25% and 50% hypo-osmotic (HYPO) shock. Quantification is based on the signal intensity ratio relative to the corresponding immunoprecipitated Cav1 protein levels. **(d)** Co-immunoprecipitation of endogenous JAK1 with Cav1 and corresponding Tyr705 pSTAT3 levels in MLEC WT cells upon IFN-α stimulation in ISO, HYPO and REC conditions. Quantification for JAK1 is based on the signal intensity ratio of JAK1 relative to the intensity of immuno-precipitated Cav1 while quantification for Tyr705 pSTAT3 is based on the signal intensity ratio of Tyr705 pSTAT3 relative to the intensity of total protein obtained from strain-free blot. The corresponding representative stain-free blot is shown. All panels exhibit representative immunoblots for N=3 independent experiments. Data shown are mean values ± SEM. Statistics were performed using repeated measures multiple-comparison one-way ANOVA. *P<0.05; **P<0.01; ***P<0.001; ****P<0.0001 and ns: not significant.

### Mechanical stress enhances diffusion of Cav1 oligomers at the plasma membrane

We have initially hypothesized that the disassembly of caveolae in response to increased membrane tension would release non caveolar caveolins at the PM and coat proteins into the cytosol ^43,49^. Indeed, single-molecule fluorescence analysis had revealed that caveolae flattening induced by membrane tension surges results in the disassembly of the cavin coat into two distinct cavin-1/cavin-2 and cavin-1/cavin-3 cytosolic sub-complexes ^43,58^. In addition, we showed that an increase in membrane tension causes the EHD2 ATPase to be released from disassembled caveolae and translocated to the nucleus where it regulates gene transcription ^59^. The comprehension of the topology of Cav1 oligomers released upon caveolae flattening is crucial for understanding the functions of caveolae, but it is still limited. Caveolins could remain organized as a flat caveolar structure, as observed by deep-etch electron microscopy ^43^, or be released as non-caveolar Cav1 oligomers. Indeed, FRAP experiments have showed an increased mobile fraction of Cav1 upon mechanical stress, suggesting that a higher number of Cav1 molecules are freely diffusing outside of caveolae as reported by Sinha et al. in 2011 ^43^. Therefore, we aimed to investigate the kinetics and dynamics of Cav1 molecules in response to mechanical stress. To monitor the fate of single Cav1 molecules with high spatiotemporal resolution at the PM, we performed single particle tracking (spt) coupled with photoactivation localization microscopy (PALM) using total internal reflection fluorescence microscopy (TIRF) ^60–62^. By fusing Cav1 with the photo-switchable mEos3.2 fluorophore, we were able to conduct high frequency sptPALM acquisitions (50 Hz) and to analyze thousands of reconstructed mEos3.2-Cav1 trajectories. Cav1 trajectories were sorted according to their diffusion mode (diffusive, confined or immobile; Fig. 3a), and diffusion coefficients (D) were computed (Fig. 3b; Methods).

**Figure 3:**
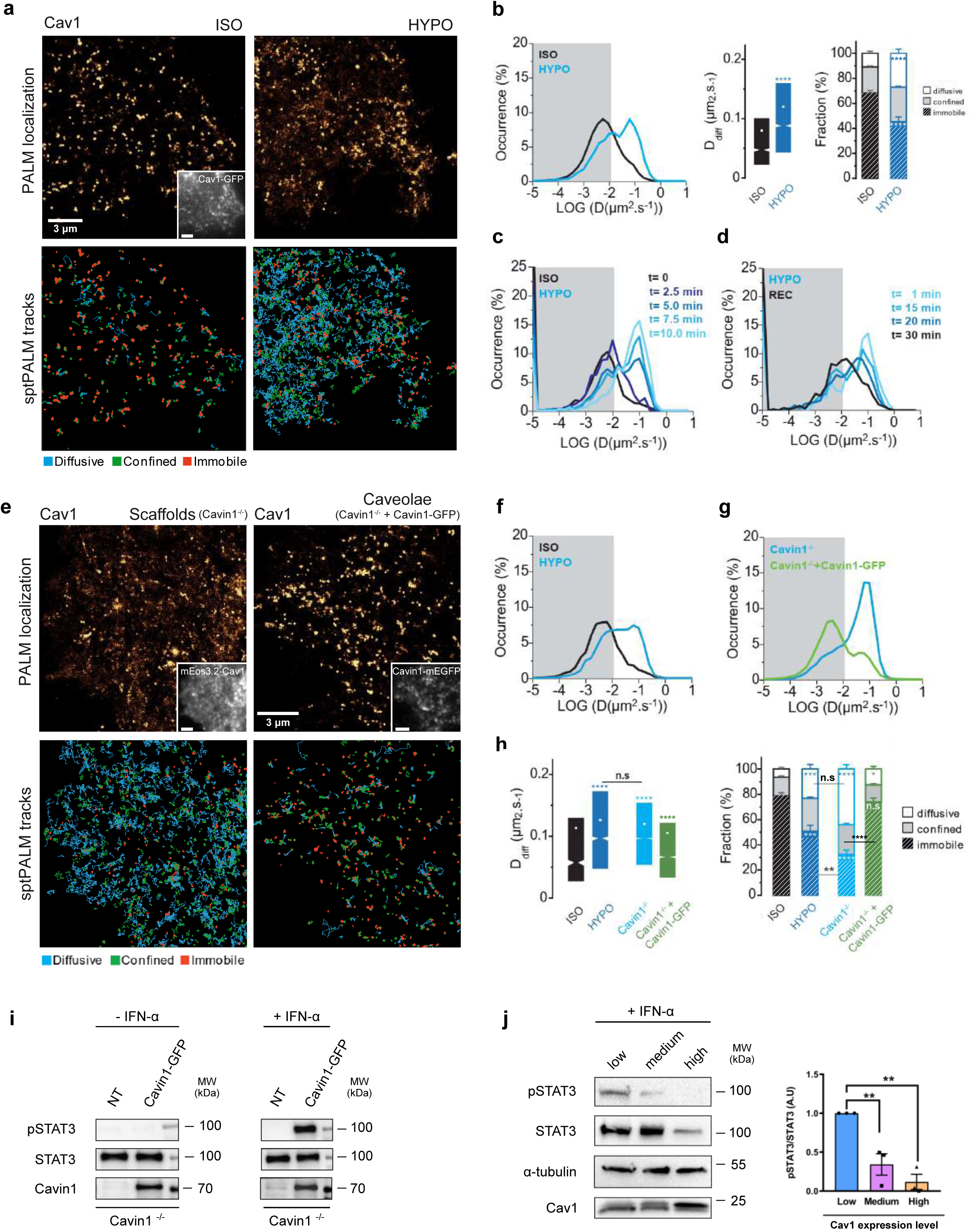
Upon osmotic shock, Cav1 is released from caveolae and undergoes lateral free diffusion along the plasma membrane. **(a)** Top: Super-resolution PALM intensity images of mEos3.2-Cav1 in MLEC WT cells in isotonic (ISO) medium (left) and during hypotonic (HYPO) shock (right) obtained from a sptPALM sequence (50 Hz, >80 s). Inset: low resolution image of Cav1-GFP, which was co-expressed for caveolae labelling (scale bar: 3 µm). Bottom: corresponding color-coded trajectories of Cav1 in the same cell showing the various diffusion modes: free diffusion (blue), confined diffusion (green) and immobilization (red). **(b)** Distribution of the diffusion coefficient D computed from the trajectories of mEos3.2-Cav1 in MLEC WT cells in ISO (black) and HYPO (blue) conditions, shown in a logarithmic scale. The grey area including D values inferior to 0.011 µm².s-1 corresponds to immobilized proteins. Data shown are mean values ± SEM. Box plots displaying the median (notch) and mean (square) ± percentile (25–75%) of diffusion coefficients corresponding to the free diffusive trajectories of mEos3.2-Cav1 in MLEC WT cells under ISO and HYPO conditions. Fraction of mEos3.2-Cav1 undergoing free diffusion, confined diffusion or immobilization in MLEC WT cells under ISO and HYPO conditions. **(c and d)** Distribution of the diffusion coefficient D computed from the trajectoies of mEos3.2-Cav1 in MLEC WT cells subjected to HYPO condition between t=0 mins to t=10 mins **(c)** and in MLEC WT cells initially at HYPO condition which are subsequently subjected to REC condition between t=1 min to t=30 mins **(d)**. **(e)** Super-resolution PALM intensity images of mEos3.2-Cav1 in MEF Cavin1^-/-^ cells (left) and MEF Cavin1^-/-^ cells rescued with transfected Cavin1-mEGFP (right) obtained from a sptPALM sequence (50 Hz, >80 s). Inset: low resolution image of mEos3.2-Cav1 (left) and Cavin1-mEGFP (right), for caveolae labelling (scale bar: 3 µm). Corresponding color-coded trajectories of caveolin-1 in the same cell showing the various diffusion modes: free diffusion (blue), confined diffusion (green) and immobilization (red). **(f)** Distribution of the diffusion coefficient D computed from the trajectories of mEos3.2-Cav1 in MEF WT cells in ISO (black) and HYPO (blue) conditions, shown in a logarithmic scale. The grey area including D values inferior to 0.011 µm².s-1 corresponds to immobilized proteins. Data shown are mean values ± SEM. **(g)** Trajectories of mEos3.2-Cav1 in MEF Cavin1^-/-^ cells (blue) and MEF Cavin1^-/-^ cells rescued with transfected Cavin1-mEGFP (green). **(h)** Box plots displaying the median (notch) and mean (square) ± percentile (25–75%) of diffusion coefficients corresponding to the free diffusive trajectories of mEos3.2-Cav1 in MEF WT cells under ISO and HYPO conditions and in MEF Cavin1^-/-^ cells transfected or not with Cavin1-mEGFP. Fraction of mEos3.2-Cav1 undergoing free diffusion, confined diffusion or immobilization in MEF WT cells under ISO and HYPO conditions and in MEF Cavin1^-/-^ cells transfected or not with Cavin1-mEGFP. **(i)** Immunoblots for STAT3 phosphorylation levels in MEF Cavin1^-/-^ cells (NT) and MEF Cavin1^-/-^ cells transfected with Cavin1-mEGFP (Cavin1-mEGFP) at steady state (left) or upon IFN-α stimulation (right). **(j)** Representative immunoblots of IFN-α induced STAT3 phosphorylation levels in MEF Cavin1^-/-^ cells expressing either low, medium or high levels of Caveolin-1 and quantification of signal ratio relative to the "low" condition. In **(a-h)**, data shown for mEos3.2-Cav1 diffusions in MLEC cells (ISO, n=9; HYPO, n=10), MEF WT cells (ISO, n=8; HYPO, n=8), in MEF Cavin1-/- cells rescued with transfected Cavin1-mEGFP (ISO, n=11) and MEF Cavin1-/- cells (ISO, n=9) are pooled from N=2 independent experiments. Statistical significance was obtained for **(a-h,j)** using multiple-comparison ANOVA. ****P<0.0001, ***P<0.001, **P<0.01, *P<0.05 and ns, not significant.

During the resting state, mEos3.2-Cav1 displayed a large fraction of immobile trajectories primarily confined to static mEos3.2-Cav1 structures, indicating that they are most likely confined within *bona fide* caveolae (Fig. 3a,b and Supplementary Video 1). When we induced an acute increase in membrane tension through hypo-osmotic shock, we observed a dramatic increase in the fraction of diffusive mEos3.2-Cav1 trajectories that displayed fast free diffusion (Fig. 3a,b and Supplementary Video 2). Importantly, the population of diffusive mEos3.2-Cav1 trajectories increased with the duration of the hypo-osmotic shock (Fig. 3c), in agreement with the visualization that mEos3.2-Cav1 trajectories explored a wider area with time. Furthermore, after subjecting cells to a hypo-osmotic shock and then returning them to iso-osmotic conditions (i.e., recovery), the population of diffusive mEos3.2-Cav1 decreased to levels similar to those recorded during the resting state (Fig. 3d). This suggests that the disassembly process is reversible and that Cav1 molecules become immobilized again within caveolae upon their reassembly. Notably, the hypo-osmotic shock did not affect the diffusion of a plasma membrane targeting sequence containing the CAAX motif (CAAX-mEos3.2), which was used as a control for bulk membrane dynamics (Extended Data Fig. 2a). Taken together, these results indicate that in response to mechanical stress, Cav1 molecules initially immobilized within caveolae are released into the PM in a highly dynamic and reversible manner, as evidenced by the kinetics of their trajectories.

### Cav1 oligomers interact with JAK1 to modulate the JAK-STAT pathway

Differentiating between the functions of caveolae and caveolin oligomers in various cellular processes has been a persistent challenge in the field of caveolae ^1,2,39,63^. In this context, we aimed to investigate whether the pool of Cav1 that interacts with JAK1 in response to changes in membrane tension is non-caveolar in nature. When cavin1 is absent, Cav1 is unable to assemble into morphologically distinguishable caveolae and remains instead as a pool of non-caveolar Cav1 with increased lateral mobility within the plasma membrane ^9^. Consequently, we used mouse embryonic fibroblasts knocked out for the *PTRF* gene encoding cavin1 (MEF Cavin1^-/-^) ^42^ and conducted sptPALM microscopy. We observed a higher fraction of the diffusive mEos3.2-Cav1 trajectories that displayed faster free diffusion in MEF cells devoid of Cavin1 (Fig. 3e,g,h). It is noteworthy that the proportion of diffusive and immobile Cav1 molecules, as well as the diffusion coefficients, were similar to those measured in WT MEF cells during hypo-osmotic shock (Fig. 3f,h). The exogenous expression of Cavin1, which allows *de novo* formation of caveolae in these cells, had a major effect on the diffusion parameters of Cav1 molecules, as it drastically reduced the fraction of diffusive Cav1 molecules to levels similar to that measured in WT MEF cells at rest (Fig. 3e-h). We next investigated the impact of highly diffusive Cav1 molecules on JAK/STAT signaling. For this, we measured the level of STAT3 phosphorylation in MEF Cavin1*^-/-^* cells stimulated or not with IFN-α. As expected, unstimulated cells did not exhibit any activation of STAT3 (Fig. 3i). While the stimulation of MEF Cavin1^-/-^ cells by IFN-α failed to activate STAT3, the reintroduction of cavin1 in these cells was sufficient to restore the IFN-α- induced activation of STAT3 (Fig. 3i). Finally, we used MEF Cavin1^-/-^ cells re-expressing different levels of Cav1 oligomers and found an inverse correlation between the amount of Cav1 oligomers present in the cells and the level of STAT3 activation (Fig. 3j). These results strongly suggest that non-caveolar Cav1 is solely responsible for the inhibition of STAT3 activation by IFN-α.

### Nanoscale imaging of Cav1 oligomers under mechanical stress

Recent advancements in super-resolution microscopy and machine-learning have provided novel insights into the nanoscale organization of various subcellular structures ^64–69^. We used stochastic optical reconstruction microscopy (STORM) to capture images of endogenous Cav1 and Cavin1 in MLEC cells and could visualize circular structures with an apparent diameter ranging from 50-100 nm, which is consistent with the known size of caveolae (Fig. 4a and Extended Data Fig. 3a,b). In addition, the use of an astigmatic lens with HILO illumination allowed to visualize caveolae in the three-dimensional space (Extended Data Fig. 3c). The disassembly of caveolae during hypo-osmotic shock is likely a two-step process in which budded caveolae first flatten out before completely releasing the coat components ^43^. Consistent with this, the observed diameter and area of Cav1-positive structures increase during hypo-osmotic shock, probably reflecting the flattening of caveolae (Extended Data Fig. 3a). Upon returning to iso-osmotic conditions (recovery), the diameter and area of caveolae decreased to their normal size as expected from the reassembly of *bona fide* budded caveolae.

**Figure 4:**
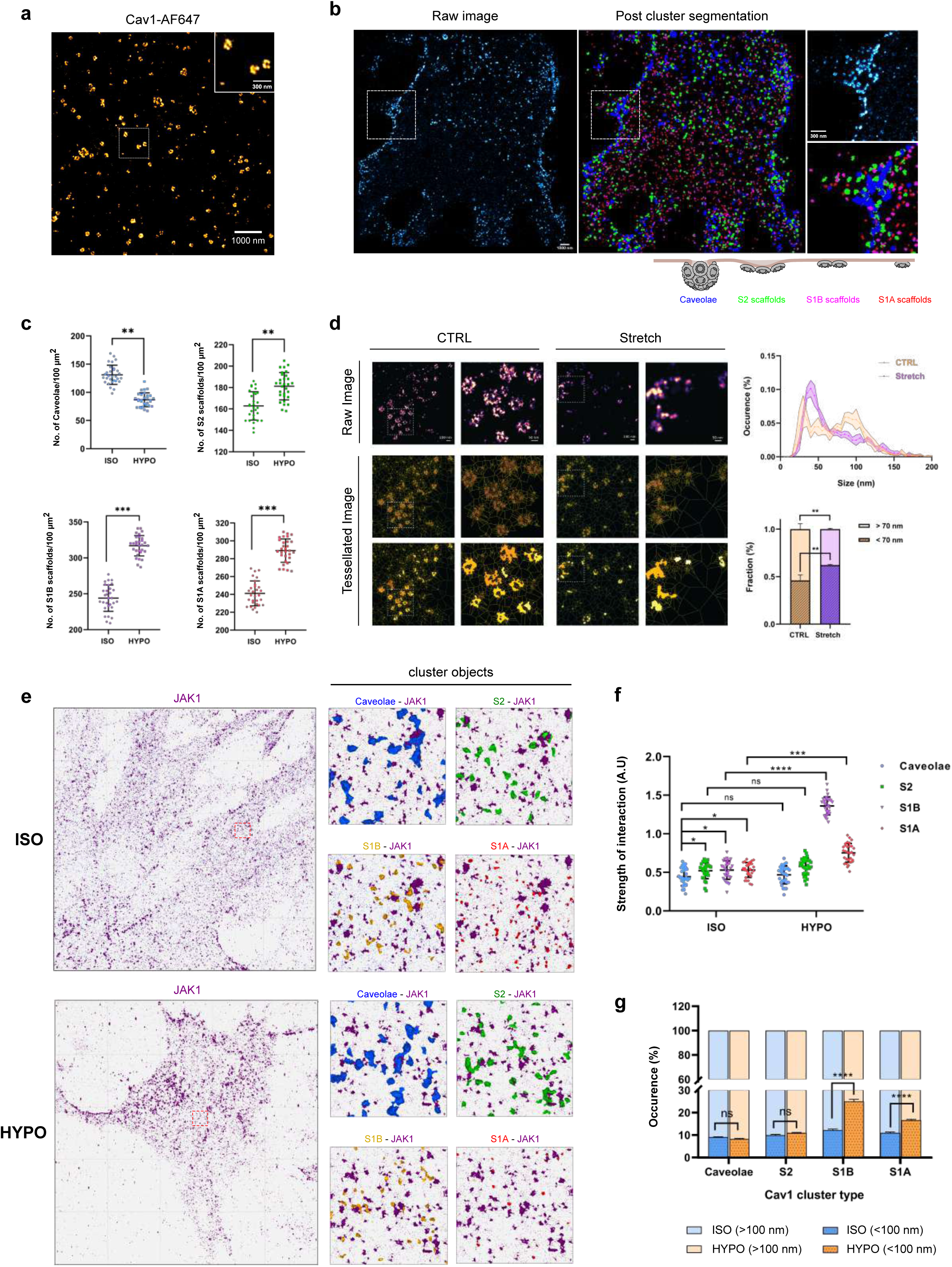
Nanoscale visualization of Cav1 and its fate in response to mechanical stress. **(a)** Representative 2D single-color dSTORM image of a MLEC WT cell stained for Cav1. Inset is higher magnification. **(b)** A comparison of representative raw 2D STORM image vs processed image to showcase the reliability and integrity of the 3D SMLM network analysis used to classify Cav1 clusters as caveolae and non-caveolar Cav1 scaffolds (S2, S1B and S1A). **(c)** Quantification per unit area of number of caveolae and the three subclasses of non-caveolar Cav1 scaffolds from post-cluster segmentation images of MLEC WT cells in ISO (n=30) and HYPO (n=30) conditions imaged using 3D dSTORM. Data shown are mean values ± SEM pooled from ROIs from N=3 independent experiments (15 cells each for ISO and HYPO). **(d)** DNA-PAINT images (top) of caveolin-1 in MLEC Cav1^-/-^ cells re-transfected with Cav1-GFP at steady state (CTRL) and under conditions of cyclic stretch (30% strain, 0.5 Hz for 30 min) on a PDMS stretching device (Stretch). Images show first-rank order density maps (middle) from SR-Tesseler, normalized by the average localization density. The corresponding detected clusters are show in bottom panel, highlighted with colors. The area outlined in the left panel is shown at a higher magnification in the right panel for the respective conditions. Cluster size distribution of Caveolin-1 in cells without (orange) or with (magenta) cyclic stretch. Bold lines represent mean values of n_CTRL_=1929 and n_Stretch_=3531 clusters pooled from N=3 independent experiments. Fraction of clusters of Caveolin-1 below 70 nm (dashed) and above 70 nm (smooth) in cells without (orange) or with (magenta) cyclic stretch. Data shown are mean values ± SEM. **(e)** Representative 3D STORM image of localizations of JAK1 and the various Cav1 populations superimposed with the JAK1 localizations within a defined ROI under iso-osmotic (top) and hypo-osmotic shock (bottom) conditions. The objects generated for clusters of JAK1 and Cav1 along with the extracted centroids for the corresponding pairs of objects are also depicted. Data shown are mean values ± SEM from ROIs in ISO (n=15 cells) and HYPO (n= 15 cells) pooled from N=3 independent experiments for the different Cav1 populations. **(f)** Interaction analysis using the MOSAIC suite plugin for FIJI representing the strength of interaction/affinity for each population of Cav1 clusters with JAK1. Data shown are mean values ± SEM from ROIs in ISO (n=15 cells) and HYPO (n= 15 cells) pooled from N=3 independent experiments for the different Cav1 populations. **(g)** Histogram depicting the % occurrence of JAK1 objects <100 nm and >100 nm from objects of the various Cav1 clusters (Caveolae, S2, S1B and S1A) under iso-osmotic and hypo-osmotic shock conditions. Data from various defined ROIs (2 per cell) in ISO (n=15 cells) and HYPO (n=15 cells) pooled from N=3 independent experiments were used to generate the histogram plot. Statistical significances were obtained using **(c)** two tailed unpaired t-test **(d, f, g)** using repeated measures multiple-comparison ANOVA. ****P<0.0001, ***P<0.001, **P<0.01, *P<0.05 and ns, not significant.

It has been proposed that Cav1 forms oligomers outside of caveolae, assembling into what are referred to as Cav1 scaffolds ^39^. However, these scaffolds are too small to be visually detectable using conventional fluorescence microscopy or electron microscopy techniques. Recently, multi-proximity threshold network analysis was applied to single molecule localization microscopy (SMLM) data acquired from PC3 cells that naturally lack cavin1 and from cavin1 transfected PC3 cells ^40^. This study allowed the classification of caveolae along with three distinct classes of Cav1 scaffolds. The classification was achieved using weakly supervised machine learning through cluster-based feature analysis, taking into account various parameters including size, shape, topology, hollowness, network characteristics, and oligomerization state. The smallest S1A scaffold is proposed to correspond to the previously identified 8S complex of SDS-resistant 11 Cav1 protomers, which are the minimal building blocks required to assemble the final 70S complex of budded caveolae ^17,70,71^. It has been proposed that the smallest S1A scaffolds can dimerize to form S1B scaffolds, whereas larger S2 scaffolds would correspond to a hemispherical intermediate made of several S1A scaffolds ^41^. We applied the same computational network modeling and machine learning based 3D pattern analysis to 3D STORM Cav1 localizations to enable the nanoscopic identification and visualization of caveolae and Cav1 scaffolds in resting MLEC WT cells. In addition to identifying *bona fide* caveolae, our post cluster segmentation allowed the visualization of numerous S1A, S1B and S2 Cav1 scaffolds (Fig. 4b). Based on the recent cryo-electron microscopy structure of human Cav1 at 3.5 Å, the minimal 8S assembly complex is composed of 11 Cav1 protomers ^17^. Previous quantitative TIRF studies have estimated that there are 144 ± 39 Cav1 copies per caveola ^72^, suggesting that *bona fide* caveolae are likely assembled by thirteen 8S complexes or S1A scaffolds ^41^. By extrapolating these estimates to our own data, we found that in resting cells, approximatively 49% of Cav1 molecules were present in caveolae, while the mean number of Cav1 molecules in S2 scaffolds, S1B scaffolds and S1A scaffolds accounted for 30%, 14%, and 7%, respectively (Supplementary Table 1). We examined the impact of mechanical stress on the distribution of caveolae and Cav1 scaffolds and observed a rapid and drastic effect upon 5 min hypo-osmotic shock. While the number of budded caveolae decreased by about 34% in line with our previous findings ^43^, we observed a concomitant increase in the number of Cav1 scaffolds, particularly in the S1A and S1B populations, which increased by 20% and 30%, respectively (Fig. 4c and Extended Data Fig. 3d; Supplementary Table 2).

To ensure the accuracy and reliability of the results, it was important to confirm these data using another super resolution microscopy technique. To this end, we employed DNA-based point accumulation for imaging in nanoscale topography (DNA-PAINT) to visualize the different Cav1 populations. DNA-PAINT relies on the stochastic binding of a fluorescent single-stranded DNA (imager strand) to the target-bound complement (docking strand) with sub-10 nm spatial resolutions ^73^. To image Cav1, we used an anti-GFP nanobody that was functionalized with a DNA strand in MLEC Cav1^-/-^ cells expressing Cav1-GFP. We performed DNA-PAINT to achieve super resolution imaging and investigate the structural organization of the different Cav1 populations at rest and after live cyclic stretching using a stretching device compatible with super-resolution microscopy ^73^. At rest, we were able to discern two broad and distinct distributions of nano-objects. The first and larger population (54%), with a size above 70 nm, is likely to correspond to *bona fide* caveolae. The second population comprising 46% of the total and with a size below 70 nm, is likely to represent Cav1 scaffolds (Fig. 4d). After subjecting the cells to a 30% live uni-axial cyclic stretching at a frequency of 0.5 Hz for 30 min, we observed a significant increase in the population of Cav1 scaffolds, which now represent 64% of the total Cav1 structures (Fig. 4d and Extended Data Fig. 4a,b). This occurred at the expense of *bona fide* caveolae, similar to what was observed by 3D STORM during hypo-osmotic shock experiments (Fig. 4c and Extended Data Fig. 3d). The smallest Cav1-GFP structures detected using DNA-PAINT have sizes of around 25 nm, which exceeds the experimental spatial resolution. This suggests that the smallest caveolar entities detected by DNA-PAINT are S1A scaffolds and not individual caveolins. Altogether, these data confirm that membrane tension surges induced by cell swelling or cyclic stretching lead to a reduction in *bona fide* caveolae. Furthermore, we demonstrate that caveolae flattening and disassembly are followed by the release of Cav1 oligomers that are assembled into scaffolds.

### Preferential interaction of S1A and S1B Cav1 scaffolds with JAK1 under mechanical stress

Next, we investigated whether there were preferential interactions between any of the Cav1 scaffolds and JAK1 in response to mechanical stress. For this, we used multicolor 3D STORM combined with spectral demixing to localize endogenous Cav1 and JAK1 proteins simultaneously (Extended Data Fig. 3b and Extended Data Fig. 4c). We then applied machine learning and network features-based analysis to categorize Cav1 clusters and visualize them as caveolae and Cav1 scaffolds. Next, we analyzed the interaction between Cav1 clusters and JAK1 based on their nanoscale proximity in MLEC cells at steady state, as well as in cells subjected to a 5-minute hypo-osmotic shock. Analysis of the interaction strength using the MosaicIA plugin ^74^ revealed a significant increase in the interaction index/score of JAK1 with the S1A and S1B scaffolds in response to hypo-osmotic shock, while the interaction with caveolae and S2 scaffolds remained minimal and unchanged (Fig. 4e,f). Furthermore, we generated objects of JAK1 and Cav1 clusters using PoCA software and used their centroids to calculate the distance between pairs of objects (i.e. between JAK1-Caveolae, JAK1-S2, JAK1-S1B and JAK1-S1A) (Fig. 4e and Extended Data Fig. 4d). The analysis of the occurrence of object pairs being at a centroid-to-centroid distance less than 100 nm within a defined ROI corroborates our findings from the interaction analysis wherein the non-caveolar scaffolds S1B and S1A demonstrate increased proximity to JAK1 in response to hypo-osmotic shock (Fig. 4g). We can speculate that at rest, Cav1 molecules may be shielded from its interaction with JAK1 when they are mainly assembled into caveolae or S2 scaffolds. Hypo-osmotic shock or stretching can significantly increase the proportion of highly diffusive Cav1 as S1A and S1B scaffolds. These scaffolds may be more accessible to JAK1, leading to a physical encounter between the two proteins. These results suggest that changes in membrane tension regulate the proportion of caveolae and Cav1 scaffolds at the plasma membrane. Cav1 scaffolds are more diffusive than caveolae and have exposed CSDs ^17^, which promote their interaction with JAK1 to negatively regulate the JAK/STAT signaling pathway.

### The caveolin scaffolding domain mediates the interaction between Cav1 and JAK1

STAT3 activation by IFN-α was inversely correlated with the level of Cav1 interaction with JAK1 suggesting that Cav1 can act as a negative modulator of JAK1 activity and thereby, STAT3 phosphorylation. The caveolin scaffolding domain (CSD), consisting of NH2-terminal residues 82-101 of Cav1, has been proposed to bind specifically to a limited set of signaling effectors mainly to exert an inhibitory effect ^29,30,75,76^. To determine whether the CSD mediates the interaction between Cav1 and JAK1, we expressed in MLEC Cav1^-/-^ cells a Cav1 construct carrying CSD-inactivating point mutations at Phe92 and Val94 (Cav1-RFP F92A/V94A) ^38^. In cells expressing Cav1 F92A/V94A, JAK1 was not detected in the immunoprecipitated fractions, whereas it was found to co-precipitate with Cav1 WT (Fig. 5a).

**Figure 5:**
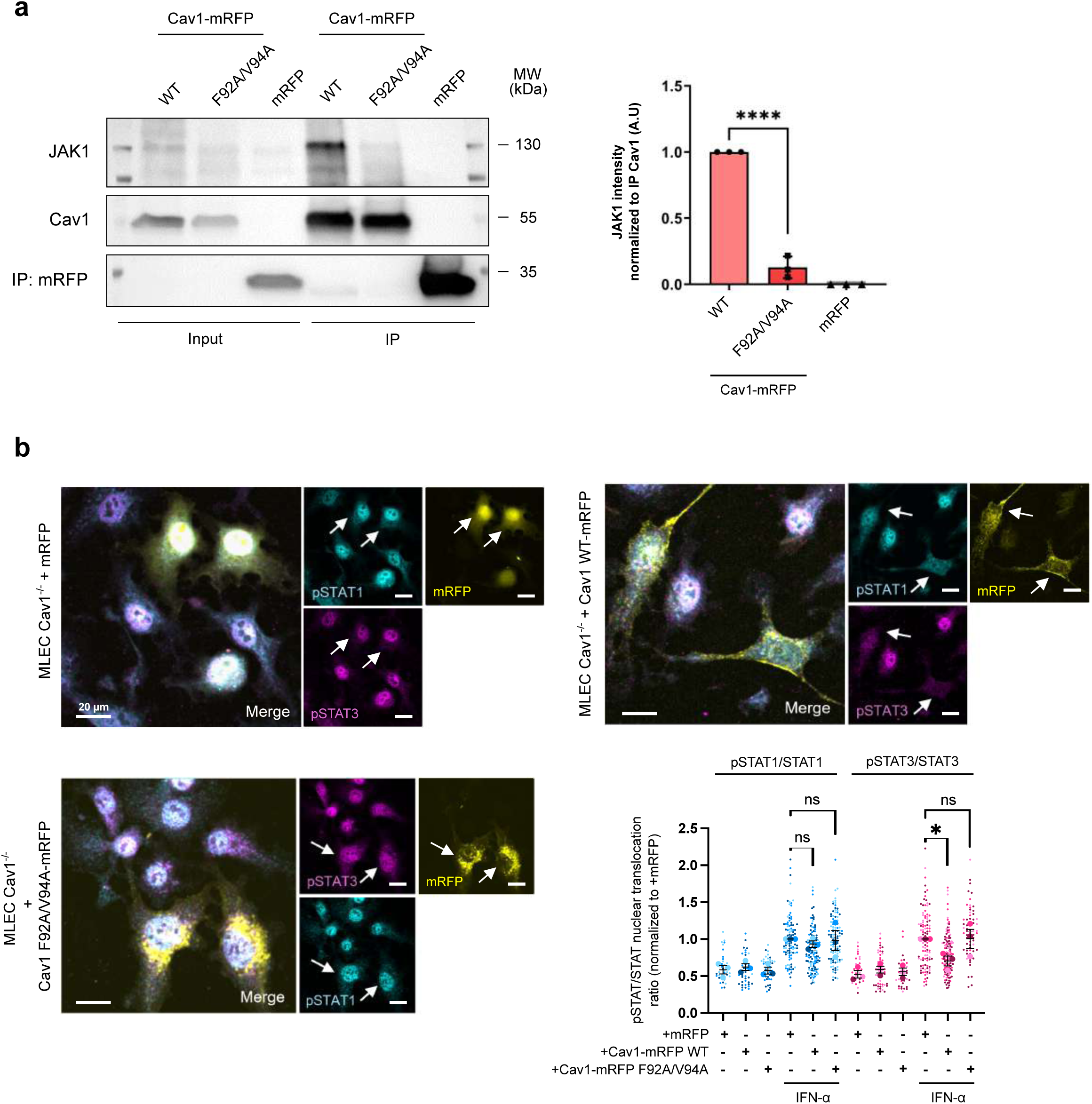
Cav1-JAK1 interaction requires the caveolin scaffolding domain. **(a)** Representative immunoblot and quantification of RFP-trap pull-down performed on MLEC Cav1^-/-^ cells expressing either Cav1(WT)-mRFP or Cav1(F92A/V94A)-mRFP or mRFP. Data shown are mean values ± SEM from N=3 independent experiments. **(b)** Representative immunofluorescence images of nuclear translocation of Tyr701 pSTAT1 (cyan) and Tyr705 pSTAT3 (magenta) in IFN-α stimulated MLEC Cav1^-/-^ cells expressing either exogenous mRFP, Cav1(WT)-mRFP or Cav1(F92A/V94A)-mRFP (yellow). Statistical significances were obtained using **(a,b)** multiple-comparison ANOVA. ****P<0.0001, *P<0.05 and ns, not significant.

We also monitored the level of pSTAT1 and pSTAT3 nuclear translocation induced by IFN-α stimulation in MLEC Cav1^-/-^ cells re-expressing either Cav1 WT or Cav1 F92A/V94A (Fig. 5b). Expression of either Cav1 WT or Cav1 F92A/V94A had no effect of the level of basal STAT3 (Extended Data Fig. 1b). The level of pSTAT1 and pSTAT3 nuclear translocation was comparable between MLEC Cav1^-/-^ cells expressing Cav1 WT and MLEC WT cells (Fig.1d). In non-transfected cells, pSTAT3 nuclear translocation occurred normally; however, in cells overexpressing Cav1 WT, a lack of pSTAT3 nuclear translocation was observed, supporting the inhibitory effect of Cav1 on JAK1 activity. In cells expressing the Cav1 double mutant F92A/V94A, the nuclear translocation of pSTAT3 was restored upon IFN-α stimulation, indicating that the inability of F92A/V94A Cav1 to negatively regulate STAT3 activation was presumably due to its inability to interact with JAK1 (Fig. 5a,b). In agreement with the insensitivity of STAT1 activation to mechanical stress and caveolae, the nuclear translocation of pSTAT1 induced by IFN-α was unaffected, regardless of whether cells express Cav1 WT or Cav1 F92A/V94A (Fig. 5b).

We further investigated the function of the Cav1 CSD using two peptides that mimic the CSD, the CavTratin peptide corresponding to the Cav1 CSD (Cav1 ^82^DGIWKASF**<u>TTF</u>**TVTKYWFYR^101^) and a peptide named CavNoxin (Cav1^82^DGIWKASF**<u>AAA</u>**TVTKYWFYR^101^) where key amino acids of the CSD are replaced by alanines, thus abolishing its inhibitory action ^77^. Upon treatment of MLEC WT with CavTratin, we observed a significant decrease of pSTAT3 levels upon IFN-α stimulation, as compared to cells treated with a control scrambled peptide (Fig. 6a), indicating that the CSD domain of Cav1 is sufficient for negatively regulating STAT3 activation. Conversely, cells treated with CavNoxin showed a significant increase of IFN-α-induced pSTAT3. Overexpression of Cav1 can generate a pool of non-caveolar Cav1 at the plasma membrane presumably due to an imbalance in the stoichiometry between the number of Cav1 molecules and the other caveolar components required for caveolae assembly, most likely cavin-1 ^37,71^. Conversely, the mutated Cav1 CSD peptide CavNoxin relieves JAK1 inhibition, most likely by competing or associating with endogenous Cav1. To exclude the possibility of the involvement of an unknown third-party interacting partner, we directly assessed the impact of CSD binding on JAK1’s ability to catalyze ATP hydrolysis in a cell free assay. The catalytic activity of human recombinant JAK1 as measured by the conversion of ATP to ADP was maintained with increasing concentrations of the control peptide. In contrast, the addition of CavTratin to the reaction mix resulted in a significant and dose dependent decrease of ADP production by JAK1 (Fig. 6b).

**Figure 6:**
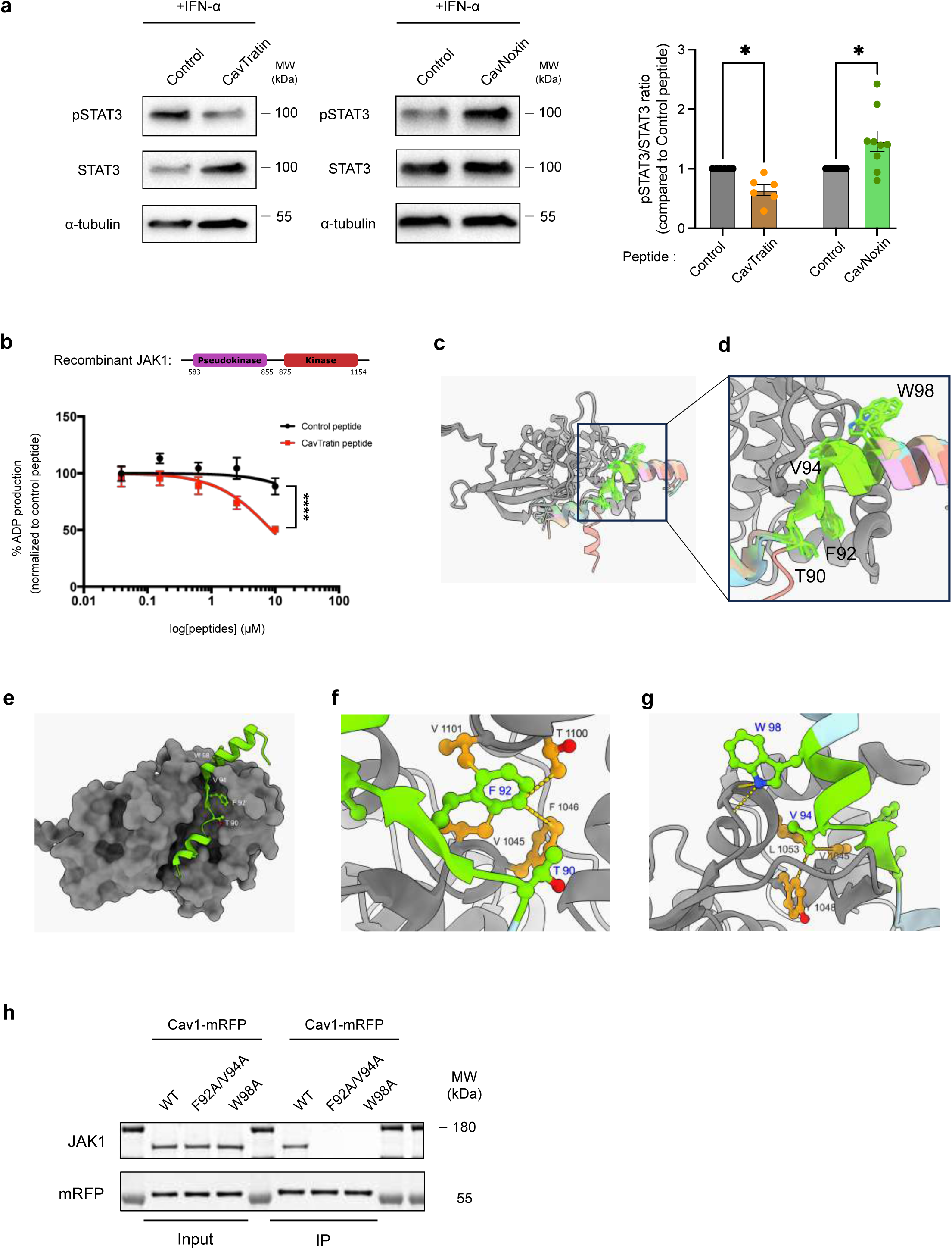
Cav1/JAK1 complex structure as predicted by AlphaFold3. **(a)** Representative immunoblots of levels of Tyr705 pSTAT3 upon IFN-α stimulation in MLEC WT cells treated with either control peptide and CavTratin (right) or control peptide and CavNoxin (left). Data shown are N=6 (CavTratin) and N=9 (CavNoxin) independent experiments. **(b)** In-vitro ADP production via ATP conversion by recombinant JAK1 relative to peptide log concentration (µM). **(c)** Superimposition of the five structures predicted by AlphaFold3 (AF3) using the JAK1[855–1154] kinase domain and the Cav1[80–110] peptide, shown in cartoon representation. JAK1[855– 1154] is coloured in grey in all models, while Cav1 [80–110] is shown in different colours for the five structures excepted the Cav1 90–98 segment highlighted in green in all five structures. **(d)** Superimposition of the side-chain conformations of Cav1 residues T90, F92, V94, and W98 in the five AF3-predicted models. **(e)** Top-ranked AF3-predicted structure of the JAK1[855–1154]–Cav1[80–110] complex. JAK1 is shown in surface representation (grey), and the Cav1 peptide is shown in cartoon representation (green). Side chains of Cav1 residues T90, F92, V94, and W98 are shown in ball-and-stick representation and coloured by heteroatom. **(f)** Close-up view of the JAK1 residues forming the binding pocket for Cav1 residue F92 in the AF3-predicted model. **(g)** Close-up view of the interactions involving Cav1 residues V94 and W98 with the JAK1 kinase domain. In panels **(f)** and **(g)**, JAK1 is shown in cartoon representation with side chains displayed as orange ball-and-stick, while the Cav1 peptide is shown in green. **(h)** Representative immunoblots of co-immunoprecipitation of Cav1-mRFP with endogenous JAK1 in MLEC Cav1^-/-^ cells over-expressing either Cav1(WT)-mRFP or Cav1(F92A/V94A)-mRFP or Cav1(W98A)-mRFP under ISO condition. Statistics significances were obtained using **(a)** two-tailed paired Wilcoxon t-test and **(b)** comparison of non-linear regression curves; *P<0.05; ****P<0.0001 and ns: not significant.

A consensus sequence known as the caveolin binding motif (CBM), identified in several Cav1-interacting proteins, has been proposed to mediate interactions with the Cav1-CSD ^78,79^. Although the existence of a *bona fide* CBM has been debated ^80^, we searched for similar motifs in JAK1 based on analogies with previously reported CBMs. Upon sequence analysis of JAK1, we identify three sequences as potential candidates: one within the FERM domain (^157^YLFAQGQY^164^), another in the pseudokinase domain (^777^WSFGTTLW^784^), and a third within the tyrosine kinase domain (^1065^WSFGVTLH^1072^). We ruled out the first sequence as it is not present in the recombinant JAK1 used to measure JAK1 catalytic activity *in vitro* (Fig. 6b). Co-immunoprecipitation experiments revealed that the sequence 1057-1071, and not the sequence 1072-1154, was required for the interaction between JAK1 and endogenous Cav1 (Extended Data Fig. 5a). Altogether, these data highlight the essential role of the Cav1-CSD in mediating the direct interaction between Cav1 and JAK1, most likely through binding to a domain located in the tyrosine kinase domain of JAK1. This interaction leads to a dose-dependent inhibitory effect of Cav1 on JAK1 catalytic activity and subsequent downstream signal transduction.

These experimental results prompted us to investigate the possible binding mode of Cav1 at the surface of JAK1. To this end, we used AlphaFold3 (AF3) to predict a structural model of the complex between the two proteins ^81^. For these predictions, we simplified the system by including only the JAK1 kinase domain (residues 855–1154) together with a series of Cav1 peptides of varying lengths, all encompassing residues 90–98 of the Cav1-CSD monomer. Restricting JAK1 to its kinase domain was supported by our experimental observation that removal of its C-terminal region (residues 1057– 1154) abolishes interaction with Cav1, indicating that Cav1 interacts with JAK1 through its kinase domain (Extended Data Fig. 5a). We focused on Cav1 peptides encompassing residues 90–98 because, unlike the wild-type protein, the Cav1 double mutant F92A/V94A failed to interact with JAK1 (Fig. 5a,b). We chose to model Cav1 peptides rather than the full-length monomer for two reasons. First, the cryo-EM structure of the Cav1 11-mer shows that residues 90–98 are not accessible for interactions with partner proteins such as JAK1 (PDB ID: 7SC0) ^17^. Second, such interactions may require a conformational rearrangement of Cav1, as suggested by coarse-grained molecular dynamics simulations of the Cav1 11-mer ^82^.

The peptides used in AF3 predictions corresponded to the wild-type Cav1 sequence but differed in length. We found that long Cav1 sequences (residues 71–121 or longer) were consistently predicted to adopt a fully helical conformation resembling that observed in the cryo-EM structure of the Cav1 11-mer (PDB ID: 7SC0), and no credible interaction with the JAK1 kinase domain was detected. In contrast, when a shorter Cav1 sequence (residues 80–110) was used, AF3 reproducibly predicted complex structures with similar structural features across five independent runs. Superimposition of these models revealed that residues 90–98 adopt a similar position at the surface of the JAK1 kinase domain (Fig. 6c), with highly conserved side-chain orientations (Fig. 6d).

In the AF3 models, Cav1 residues F92 and V94 play a central role in the interaction by inserting into a groove at the surface of the JAK1 kinase domain (Fig. 6e). F92 occupies a hydrophobic cavity formed by JAK1 residues V1045, F1046, V1101, and the methyl group of T1100 (Fig. 6f). V94 occupies a second hydrophobic pocket lined by residues Y1048, L1053, and V1045 (Fig. 6g). Moreover, W98 forms a hydrogen bond via its HNε2 atom with the backbone carbonyl oxygen of Q1055 of the JAK1 kinase domain (Fig. 6g). No clear interaction was observed between JAK1 and Cav1 residues located on the N-terminal side of F92 or on the C-terminal side of W98 in the predicted complex. The role of W98 was further confirmed biochemically, as co-immunoprecipitation experiments showed that the Cav1 W98A mutant failed to interact with JAK1 (Fig.6h).

Although the AF3 confidence metrics for these models were low, as occasionally observed in AF3 predictions, we next assessed the stability of the predicted interface using molecular dynamics (MD) simulations in explicit solvent. Simulations were initiated from the top-ranked AF3 model of the JAK1[855–1154]–Cav1[80–110] complex and aimed (1) to relax the initial model and refine intermolecular contacts, and (2) evaluate the stability of the predicted interactions. To account for stochastic effects, three independent MD trajectories were performed using different initial velocities, yielding a total simulation time of approximately 2.1 µs. During MD simulations, the JAK1 kinase domain remained structurally stable, with the Cα atom root-mean-square deviation (RMSD) values below 2.0 Å relative to the initial AF3 model (Extended Data Fig. 5b-d). This stability is further supported by low root-mean-square fluctuation (RMSF) values predominantly between 1.0 and 2.0 Å (Extended Data Fig. 5e). In contrast, the RMSD calculated for all Cα atoms of the Cav1[80–100] peptide reached values up to 8.0 Å (Extended Data Fig. 5b-d). However, when the analysis was restricted to residues 90–98, RMSD values remained mostly below 2.5 Å. Consistently, RMSF analysis showed limited fluctuations for residues 90–98 (RMSF mostly < 2.0 Å), whereas distal regions were significantly more mobile (Extended Data Fig. 5f). This indicates that the terminal regions of the peptide undergo substantial conformational fluctuations relative to the JAK1 kinase domain, whereas the 90–98 segment remains conformationally constrained. It should be noted that the MD simulations were performed using a Cav1 peptide corresponding to residues 80-110. Therefore, we cannot exclude the possibility that the fluctuations observed in the distal regions of the peptide may be reduced in the context of the full-length Cav1 protein.

Analysis of minimum interatomic distances further supported the stability of the predicted interactions throughout the MD trajectories. Cav1 residue V94 remained closely associated with JAK1 residues V1045, Y1048, and L1053 along the three trajectories, consistent with persistent hydrophobic packing (Extended Data Fig. 5g-i). F92 remained also constantly close to JAK1 residues V1045, F1046, T1100, and V1101 in trajectory 1 and 2 (Extended Data Fig. 5j-m). In trajectory 3, F92 appeared to undergo a conformational transition that preserved only proximity with JAK1 residue V1101(Extended Data Fig. 5m). Likewise, the hydrogen bond between the side chain of Cav1 residue W98 and the backbone of JAK1 residue Q1055, as predicted by AF3, was stably maintained during MD trajectory 3 (Extended Data Fig. 5n). In contrast, fluctuations observed in trajectories 1 and 2 suggest that this residue may exist in equilibrium between multiple confirmations (Extended Data Fig. 5n).

Altogether, these results support the AF3-derived model and indicate that Cav1 likely interacts with JAK1 through its 90–98 segment, with residue V94 being at the core and forming a key interaction with a set of hydrophobic residues of JAK1. This structural model fully agrees with our biochemical data and highlight a central role for this Cav1 region in JAK1 binding.

### Multiple signaling pathways are controlled by the mechanical release of Cav1

We next asked whether other signaling pathways were regulated by this mechanism. For this, we conducted a high-throughput screening experiment using reverse phase protein array (RPPA) – a miniaturized dot-blot technology that enables proteomic analysis and identification of activated or altered signaling pathways ^83^. The RPPA analysis was performed on wild-type (WT) and Cav1 knockout (Cav1^-/-^) MLEC subjected to uniaxial stretching (Extended Data Fig. 6a). RPPA analysis revealed several other signaling pathways that were also modulated by cell stretching including the threonine phosphorylation of MAPK kinase (p42/44) and the serine phosphorylation (pSer473) of AKT kinase. While MAPK was activated by stretch irrespective of the presence of Cav1, the increase in AKT pSer473 triggered by uni-axial stretching strictly required Cav1 (Extended Data Fig. 6b). Cells were also stimulated with IFN-α or IFN-β to extend the analysis to the JAK/STAT signaling pathway. RPPA showed that IFN stimulation resulted in a significant increase of STAT3 and STAT1 phosphorylation at tyrosine 705 (Tyr705 pSTAT3) and tyrosine 701 (Tyr701 pSTAT1), respectively (Extended Data Fig. 6c). RPPA confirmed that uniaxial stretching of IFN-stimulated cells led to a significant reduction in STAT3 phosphorylation in a Cav1-dependent manner while the phosphorylation of STAT1 remained unaffected (Extended Data Fig. 6d). We also analyzed previously reported interactions between endogenous Cav1 and endothelial nitric oxide synthase (eNOS) as well as protein tyrosine phosphatase 1B (PTP1B) ^31,84^. The interaction between Cav1 and eNOS, as well as between Cav1 and PTP1B, was significantly enhanced during hypo-osmotic shock and returned to basal levels upon returning to iso-osmotic conditions (Extended Data Fig. 7a). Similarly, to JAK1 (Fig. 2b), our observations revealed a significant correlation between the strength of the hypo-osmotic shock and the intensity of Cav1 interaction with eNOS, and PTP1B (Extended Data Fig. 7b). This increased interaction most likely reflects the increase of Cav1 molecules released from disassembled caveolae. As observed for JAK1 (Fig. 2c), this effect was rapid, with the maximum interaction observed after only 5 min of hypo-osmotic shock and did not further increase with longer exposure times (Extended Data Fig. 7c). The increased interaction between eNOS and Cav1 induced by hypo-osmotic shock resulted in reduced eNOS activation, as indicated by decreased phosphorylation of eNOS at serine 1177 (Extended Data Fig. 7d) ^85^. Return to iso-osmotic conditions restored basal levels of eNOS-Cav1 interaction and eNOS activation. Hypo-osmotic shock had no effect on eNOS activation in MLEC Cav1^-/-^ cells. Additionally, we confirmed the interaction between Cav1 and phosphatase and tensin homolog (PTEN), a potent tumor suppressor that functions as a negative regulator of the AKT signaling pathway ^86^ (Extended Data Fig. 8a). During hypo-osmotic shock, the increased binding of Cav1 to PTEN correlated with an increase in AKT activation, as indicated by serine 473 phosphorylation. Although this temporal association is consistent with a possible modulation of PTEN activity by Cav1 binding, and consequently of AKT signaling, these data remain correlative. We can therefore only speculate that the increase in AKT activity, also observed by RPPA in stretched cells (Extended Data Fig. 6b), reflects inhibition of PTEN phosphatase activity by Cav1, particularly given that AKT does not interact with Cav1 (Extended Data Fig. 8b). As observed for JAK1, Cav1 interaction with PTEN also required the residues F92, V94 et W98 of the Cav1-CSD domain (Extended Data Fig. 8b). Similar to the interactions observed with JAK1, eNOS, and PTP1B, the interaction between Cav1 and PTEN returned to basal levels upon restoration of iso-osmotic conditions (Extended Data Fig. 8a). Finally, we found that Cav1 did not interact with TYK2, another member of the JAK family that is also activated by IFN-α (Extended Data Fig. 8b). Caveolae and integrins have been shown to cooperate in the regulation of mechanosignaling ^87^. We also recently established a reciprocal control between caveolae and integrins that is crucial for invadosome biogenesis and activity ^88^. However, we could not immunoprecipitate integrins with Cav1 in MLEC cells (Extended Data Fig. 8c). These results establish the selectivity of the signaling pathways regulated by caveolae mechanosignaling.

### Physical model of caveolae formation under mechanical stress

We developed a theoretical model of caveolae self-assembly based on our observations (Supplementary Note 1). The model is based on equilibrium thermodynamics ^89^, owing to the reversibility of caveolae disruption under stress. The model accounts for the presence of full caveolae and hemispherical S2 scaffolds, together with smaller S1 scaffolds accounting for both S1A and S1B (Figure 7A). Minimization of the system’s free energy (detailed in Supplementary Note 1), yields a membrane tension-dependent fraction of the total Cav1 population in the different states, which is proportional to

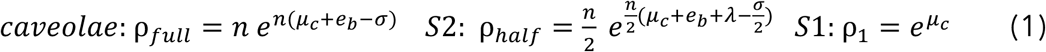

**Figure 7:**
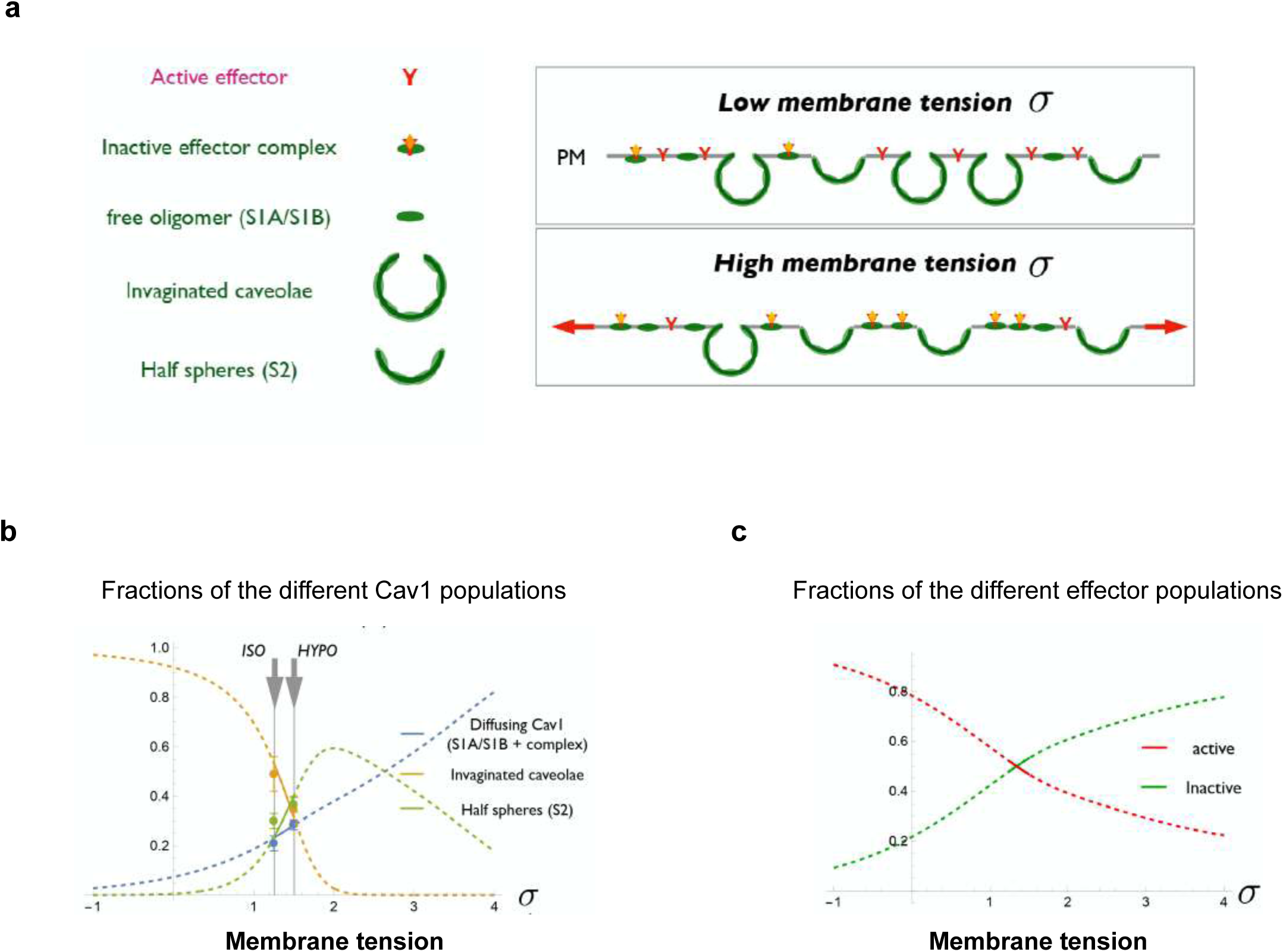
Physical model of caveolae formation under stress. **(a)** Different Cav1 and membrane-associated signaling effectors included in the physical model (free S1A/S1B oligomers, full caveolae, hemispherical S2 domains, active effectors, and complexes of Cav1 and inactive effectors). Sketch of the model (right). **(b)** Variation of the three Cav1 populations with membrane tension (the blue line represents the sum of all diffusing Cav1 states). **(c)** Variation of the fraction of active and inactive effectors with membrane tension. Solid lines are within the tension range explored in stretching experiments (data points in **b** are from Figure 4f) and dashed lines are theoretical extrapolation over a broader range of tension variation. Parameters: e_b=3.6, λ=1.76, ρ_tot=0.33, ε=2.6, ρ_(j,tot) =0.02.

where σ is the membrane tension, e_b_ is the binding energy between S1 scaffolds in larger domains (caveolae and S2), 𝜆 is the line tension associated with the boundaries of S2 domains, 𝑛 = 13 is the number of S1 in caveolae and 𝜇_𝑐_ is the Cav1 chemical potential, obtained from the conservation of the total number of Cav1 at the membrane (see below). The energetic parameters are normalized to be expressed in unit of the thermal energy 𝑘_𝐵_𝑇.

The model predicts that the fraction of Cav1 in caveolae decreases while the fraction of freely diffusing Cav1 increases in sigmoidal fashions with the membrane tension (Figure 7b). Hemispherical S2 domains exist within a limited range of membrane tension, that strongly depends on the S2 line tension. The parameters of Figure 7b are fitted to reproduce the different populations observed by super resolution (Figure 4b and 4e). Although the fit is not unique, it nevertheless suggests that the homeostatic value of the membrane tension is within the range that permits substantial variation of free Cav1 under stretch. Free Cav1 may interact with membrane-associated signaling effectors, leading to their inactivation. This is described within the same framework, where the population of effectors in the different states is proportional to

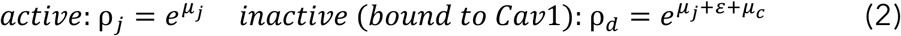

where ε is the binding energy between S1 and effectors, and 𝜇_4_ is the effector chemical potential. The two chemical potentials are obtained from the conservation of the total number of the respective proteins at the membrane: ρ_1_ + ρ_𝑑_ + ρ_𝑓𝑢𝑙𝑙_ + ρ_ℎ𝑎𝑙𝑓_ = ρ_𝑡𝑜𝑡_ and ρ_4_ + ρ_𝑑_ = ρ_4,𝑡𝑜𝑡_. Membrane tension increase leads to effector inactivation by releasing Cav1 from caveolae and S2 aggregates (Figure 7c). A more complete version of the model developed in the Supplementary Note 1 allowing for the binding of multiple effectors on free Cav1 oligomers does not qualitatively affect the picture shown in Figure 7b and 7c. Therefore, our model, based on equilibrium thermodynamics, predicts how the population of different Cav1 states varies with membrane tension and yields a quantification of remote control of signaling by caveolae. It allows to reproduce our experimental conclusions, made on the basal surface of adhered cells subjected to an osmotic shock, but also to propose quantitative predictions for a wide range of membrane tension values.

## DISCUSSION

Since their first description in 1953, research over the years has revealed a variety of roles for caveolae and/or caveolins in preserving biological functions [reviewed in ^2,22^].

Although caveolae were proposed to be involved in mechanoprotection and in maintaining cellular integrity as early as the mid-1970’s ^90–92^, it was not until our seminal discovery that caveolae sense and respond to mechanical stress that the field was prompted to reassess their classical functions in the context of cell mechanics ^2,43,50–52^. We initially hypothesized that caveolar proteins could be released upon the mechanical disassembly of caveolae, thereby mediating the mechanical response of the cell ^49^. We could indeed demonstrate that elevated membrane tension triggers the translocation of the ATPase EHD2 from the neck of caveolae to the nucleus, where it regulates gene transcription ^59^. Cavin1 is also released from the caveolar coat in response to hypo-osmotic shock ^43,58,93^. Additionally, cavin3 can interact with BRCA1 and regulate several cancer related pathways upon its release from caveolae that have been disassembled by UV exposure or hypo-osmotic shock ^94^.

Our findings reveal that mechanical stress significantly augments the extent of Cav1 interaction with JAK1, PTEN, PTP1B, and eNOS. This observation, along with the proposed role of non-caveolar Cav1 scaffolds in signaling ^39^, has led us to hypothesize that the increased interaction may be attributed to the release of Cav1 molecules originating from caveolae that have undergone mechanical disassembly. sptPALM microscopy confirmed that the primary population of non-diffusive Cav1 particles, initially constrained to confined domains under low membrane tension (likely within *bona fide* caveolae), transitioned significantly as membrane tension progressively increased. This transition was characterized by a diffusion pattern indicative of Cav1 scaffolds scattering upon caveolae disassembly, accompanied by an increased population of freely diffusing Cav1 particles and an increase in their diffusion coefficient. Additionally, in Cavin1^-/-^ cells where only non-caveolar Cav1 is present, we measured a diffusion coefficient for Cav1 particles that was strictly identical to the higher diffusion coefficient induced by hypo-osmotic shock in WT cells. These observations confirm that the highly diffusive pool of Cav1 molecules corresponds to non-caveolar Cav1. Similarly, we established a direct correlation between the level of inhibition of STAT3 phosphorylation and the quantity of non-caveolar Cav1.

The potential formation of the JAK1-Cav1 complex was evaluated using AI-based structural prediction and molecular dynamics simulations. We identified a plausible complex conformation where Cav1 residue V94 forms the core of an interacting motif that inserts into a cleft on the surface of JAK1. Cav1 residues F92 and W98 flank V94 and likely contribute to stabilize this interaction. Targeted mutagenesis confirmed the importance of these Cav1 residues for formation of the Cav1-JAK1 complex. Furthermore, this model is consistent with experimental results obtained using the JAK1Δ(1057-1154) and JAK1Δ(1072-1154) truncation mutants. Residues 1057-1071 form a helix (highlighted in red in Extended Data Fig. 5o and Supplementary Video 3) that supports a large part of the Cav1 binding site (residues 1033-1055, highlighted in cyan in Extended Data Fig. 5o and Supplementary Video 3). We therefore propose that removal of residues 1072-1154 (highlighted in orange in Extended Data Fig. 5o and Supplementary Video 3) preserves enough of the predicted Cav1 binding site to allow peptide binding, albeit with altered affinity. In contrast, deletion of the supporting helix spanning residues 1057-1071 would be expected to disrupt the Cav1 binding site and prevent the JAK1 kinase domain from binding Cav1. Together, these findings indicate that these residues are required for the functional engagement of the Cav1-CSD domain, and the AF3 modeling and MD simulations further support a direct role for the Cav1-CSD in JAK1 binding.

Super-resolution imaging and network analysis have revealed the existence of different subclasses of Cav1 scaffolds, which serve as building blocks of *bona fide* caveolae ^41^. Similar analysis of our SMLM data from cells exposed to hypo-osmotic shock or subjected to cyclic stretch provided insights into the mechanical disassembly of caveolae into Cav1 scaffolds. This disassembly was strongly correlated with a significant increase in both Cav1 diffusion and its interaction with signaling effectors. The functionality of the Cav1-CSD has been debated, as it has been proposed, based on Cav1 sequence alignment, that Cav1-CSD is a hydrophobic motif embedded in the lipids of the plasma membrane, thus preventing direct physical interactions with cytosolic proteins ^80,95^. The preferential interaction of Cav1 scaffolds, as opposed to caveolae, with signaling molecules during periods of mechanical stress provides a potential explanation for the involvement of the Cav1-CSD. Indeed, recent structural data obtained on the minimal 8S Cav1 building block complex have revealed that the CSD is positioned at the outer rim of the 8S discoid complex ^17^. Therefore, we can speculate that when caveolae are mechanically disassembled into scaffolds, along with subsequent lipid reorganization ^96^, the CSD, which may not be readily accessible within budded caveolae, could potentially be exposed and become available for binding to cytosolic proteins. In this context, the structural characterization of the Cav1 8S complex by cryo-EM has offered new insights into this interaction ^97,98^. A recent study by Doktorova and colleagues utilized the structure of the Cav1 8S complex to perform coarse-grain molecular dynamics simulations of a single Cav1 8S embedded in lipid membranes of varying compositions ^82^. During the simulations, the Cav1 8S complex localized to highly curved surfaces, leading to the exposure of several Cav1-CSD residues, including T90, V94, and W98, to the aqueous environment. Our structural predictions indicate a central role for Cav1 residue V94 in mediating interaction with the JAK1 kinase domain. Notably, the Cav1[90-98] segment, which adopts a helical conformation in the 8S Cav1 structure (PDB code 7SC0, ^17^), adopts a markedly different conformation in the predicted JAK1 complex. These observations support a model in which Cav1 undergoes a conformational transition from its 8S form, exposing residues F92, V94 and W98 of the Cav1-CSD and rendering them accessible for JAK1binding. Within this framework, our predicted structure may correspond to the final state of this transition. These data are also consistent with a recent study suggesting that the S2 and S1B Cav1 scaffolds exhibit a more exposed CSD with the surrounding molecular environment ^99^. In addition, the simultaneous release of cavins from the caveola bulb, induced by mechanical stress, may also contribute to the enhanced accessibility of Cav1-CSD. These findings provide strong evidence for the role of non-caveolar Cav1 scaffolds in the mechanical regulation of intracellular signaling.

Since the expression of cavin proteins seems limited to vertebrates, it has been proposed that caveolins can carry out their functions independently of caveolae in most organisms. Likewise, various cell types including neurons, lymphocytes, hepatocytes, and certain cancer cells do not express cavins, suggesting that caveolins may exert caveola-independent functions in these cells ^63^. Early studies have reported the ability of Cav1 to interact with several effectors through the Cav1-CSD domain ^29,30,75,76^. Using CSD mimicking peptides and point mutations in the Cav1-CSD, we provide unequivocal evidence of the CSD requirement for the direct interaction between Cav1 and JAK1. Accordingly, it has been reported that the deletion of the CSD abolished the inhibition of STAT3 phosphorylation caused by Cav1 overexpression ^100^. It is remarkable that the CSD exhibits significant primary sequence similarities to the pseudo-substrate domain of SOCS1, which mediates JAK inhibition by SOCS1 ^101,102^. In this context, it is interesting that another member of the SOCS family, SOCS3, relies on Cavin-1 for its localization at the plasma membrane in endothelial cells, and that STAT3 activation is increased when Cavin-1 is depleted ^103^. It will be interesting to test if, under mechanical stress, SOCS3 may be released from Cavin-1, which would then compete with Cav1 scaffolds to bind to JAK1.

The AF3-predicted model could provide insight into the functional significance of the Cav1–JAK1 interaction. In principle, Cav1 binding could influence JAK1 catalytic activity either through long-range effects or by altering the dynamics of the kinase domain. However, superimposition of the predicted JAK1-Cav1 complex onto the cryo-EM structure of the JAK1 dimer (PDB ID: 8EWY) ^104^ suggests an alternative mechanism. In our model, the Cav1 peptide binds to a region of JAK1 corresponding to the interface between the tyrosine kinase domains of the two monomers. This interface, which involves helical packing between the kinase domains, is thought to play a critical role in JAK1 dimerization—despite the presence of an additional interface involving the pseudo-kinase domains—and is therefore likely essential for cytokine signaling. We thus speculate that disruption of this interface, or inhibition of its formation via Cav1 binding as proposed here, could modulate cytokine signal transduction mediated by JAK1 dimerization.

JAK1 inhibition was observed even in the absence of stimulation with IFN-α, indicating that this regulatory process may be extended to other cytokines that activate JAK kinases. In the context of our previous study on IL-6/STAT3 signaling in human muscle cells, it is probable that the control exerted by caveolae mechanosensing on this signaling axis operates through the same mechanism ^53^. It is intriguing that JAK1-dependent STAT3 activation was specifically targeted while leaving STAT1 activation unaffected. Interestingly, our data indicate that TYK2, the other kinase operating in the IFN-α signaling complex, does not interact with Cav1. These findings further support the dichotomy between STAT3, which is known for its oncogenic properties, and STAT1, which is recognized as a tumor suppressor ^105^.

Our findings unveil a novel mechanism through which caveolae exert remote control over the regulation of various signaling pathways. This control takes place beyond the boundaries of the caveolae structure and relies on a dynamic exchange between caveolae and Cav1 scaffolds. The balance between these two distinct populations of Cav1 assemblies is finely tuned by variations of membrane tension and enables precise modulation of signaling outputs. The proposed mechanism finds support in a theoretical model that considers the thermodynamics of caveolae formation under mechanical stress. The model suggests that as membrane tension increases, there is a transition from caveolar Cav1 to Cav1 scaffolds, accompanied by an enhanced affinity of Cav1 towards its effectors. STORM imaging further revealed a small population of Cav1 scaffolds under resting conditions, supporting the existence of a pre-existing pool at the plasma membrane. JAK1 associates with these scaffolds, indicating that this pool is functionally engaged. These structures may arise from local caveolar disassembly triggered by spatial heterogeneities in membrane tension, as we previously reported in migrating cells ^106^. Alternatively, they may represent newly synthesized Cav1-8S complexes that arrive at the plasma membrane as scaffold intermediates prior to their assembly into mature caveolae.

Mechanical forces can regulate diverse cellular functions by directly influencing protein interactions, such as reinforcing ^107,108^ or destabilizing interactions ^43,109^ and controlling enzymatic reactions ^110^. Furthermore, mechanical deformations have been shown to uncover concealed or cryptic binding sites, as demonstrated for vinculin and talin ^111^, as well as cryptic phosphorylation ^112^ or proteolysis sites ^113^. Thus, there is a consensus that external mechanical stresses are transmitted directly and rapidly to induce local protein deformation in mechano-sensitive structures like integrin adhesions, the cytoskeleton, or the nucleus ^69,114–116^. Our findings reveal a new paradigm of mechano-transduction, challenging the notion that mechanical forces trigger immediate local effects or are rapidly transmitted through the cell via cytoskeletal elements or the membrane. Instead, we demonstrate that the conversion of *bona fide* caveolae into caveolin scaffolds generates mechanical messengers that diffuse at the plasma membrane. Cav1 scaffolds interact with signaling effectors at distant locations from the initial mechano-sensing caveolae, with a time delay. Caveolae remote mechanosignaling may also serve as a mechanism for integrating and facilitating crosstalk with other mechanosensitive structures, such as integrin adhesions and the cytoskeleton. Notably, integrin adhesion and Cav1 have been functionally interconnected ^38,117,118^ and share common cytoskeleton partners ^119^ and signaling pathways ^120–122^. In addition, caveolae have been associated with actin stress fibers and implicated in the regulation of their contractility ^32,123,124^. In this context, it is interesting that protein deformation within integrin adhesion is not directly triggered by the transmission of external stretch but mediated by a delayed acto-myosin remodeling process ^73^.

The pathophysiological implications of remote mechanosignaling by caveolae remain to be investigated. Several of the signaling effectors regulated by this process, including PTEN, PTP1B, and STAT3, are well-known for their involvement in the control of tumorigenesis. The role of Cav1 and caveolae in cancer has sparked prolonged debates due to their seemingly contradictory behavior, with reports indicating both oncogenic and tumor suppressor properties ^25,125–127^. This novel functional aspect of caveolae mechanics could have a substantial impact on tumor growth. The mechanical forces experienced by cancer cells throughout tumor progression may disrupt caveolae mechanosensing, thereby impairing the precise regulation of caveolae-mediated mechanosignaling.

Cav1 has long been known to interact with numerous proteins, including signaling molecules and membrane receptors. However, the lack of compelling evidence for the localization of these proteins inside caveolae, along with data arguing for exclusion of bulk plasma membrane proteins ^34^, raises a puzzling question regarding how caveolae can effectively regulate signaling effectors that are not found into caveolae. This study presents a new mechanistic insight into the regulation of cell signaling through caveolae. The reversible conversion of caveolae into Cav1 scaffolds enables the remote control of signaling molecules localized outside of caveolae. Taken together, our findings represent a significant breakthrough in the field of intracellular signaling, revealing caveolae as crucial mechano-signaling devices capable of remotely fine-tuning specific signal transduction processes originating from the plasma membrane. This novel understanding of caveolae not only contributes to our comprehension of their functions but also holds profound implications for human pathophysiology ^22^.

## Supporting information

Supplementary_note_Theoretical_analysis_Kailasam_et_al_2026

Movie 1: ISO

Movie 2: HYPO

## Acknowledgements

We are grateful to Dr. Anne Kenworthy, Dr. Bing Han, and Dr. Milka Doktorova for sharing their preliminary data on coarse-grain molecular dynamics ^82^ and for their stimulating discussions. We are also indebted to Irina S. Moreira (University of Coimbra, Portugal) for sharing preliminary data on docking experiments and for insightful discussions. The core facilities and the CurieCoreTech recombinant antibodies platform of Institut Curie, the scientific and technical assistance from staff of the Cell and Tissue Imaging (PICT-IBiSA) and the Nikon Imaging Centre at Institut Curie, member of the French National Research Infractucture France-BioImaging (ANR10-INBS-04) are acknowledged. We would like to thank C. Schietroma from Abbelight for technical assistance with SMLM (STORM) experiments. The help of the following people for providing materials or expertise is acknowledged: Dr. Radu. V. Stan (Dartmouth College, USA), and Dr. Miguel Del Pozo (Spanish National Centre for Cardiovascular research, Spain). This work was supported by institutional grants from the Curie Institute, Institut National de la Sante et de la Recherche Medicale, CNRS, and by specific grants from Agence Nationale de la Recherche (ANR-19-CE15-0020-02) and INCa 2018-1-PL BIO-08-ICR-1 (Decision N° 2018-154) to C.L. Funding from France Canada Research Fund: NVKA GR013361 - FCRF to C.L. and I.R.N. is acknowledged. Exchanges between the C.L. and I.R.N. laboratories are supported by an International Research Project “IRP Cav1”. G.G. was supported by the INCA (AAP PLBIO no. 2020-109), the ANR (ANR-21-CE11-0004-01), the French government in the framework of the University of Bordeaux’s IdEx "Investments for the Future" program / GPR BRAIN_2030, GPR LIGHT. O.R. was supported by the ANR (ANR-20-CE42-0003-02) and the Nouvelle-Aquitaine Regional Council (AAPR2021-2020-12041310). S.K.M was supported by a PhD fellowship from Ligue Nationale contre le Cancer, N.T. by a PhD fellowship from Ministère de l’Enseignement Supérieur et de la Recherche, M.D. by a PhD fellowship from Association Française contre les Myopathies (AFM): CAV-MUT (17151), and V.B. by a postdoctoral fellowship from Fondation de France (WB-2024-54138). The Lamaze and Sens teams are members of Labex Cell(n)scale ANR-10-LBX-0038, part of the IDEX PSL ANR-10-IDEX-0001-02. Molecular Dynamics Simulations were carried out on A100 GPU resources provided by GENCI at IDRIS@CNRS (Jean Zay supercomputer; allocation A0190715663). The funders had no role in study design, data collection and analysis, decision to publish or preparation of the manuscript.

## Author contributions statement

S.K.M., N.T., P.C. and C.M.B designed and performed the experiments, analyzed and interpreted the data, and wrote the manuscript. O.R. designed and performed the spt-PALM experiments, analyzed and interpreted the corresponding data. X.Z., F.N.V. and G.G. designed and performed the DNA-PAINT experiments. A.R., C.G., P.G.T., R.R. and M.D. performed experiments. I.B. performed RPPA analysis. I.K., I.R.N., V.B. and G.H. analyzed and interpreted the 3D STORM data using 3D SMLM network analysis. P.S. designed the physical model for theoretical validation of the study. G.G., O.R., I.K., I.R.N., P.C., P.S. and C.M.B proofread and edited the manuscript. C.L. supervised the study, designed the experiments, interpreted the data, and wrote the manuscript. All authors discussed the results and commented on the manuscript.

## Declaration of interests

The authors declare no competing interests.

## Materials and Methods

### Cell culture, transfection and cell treatments

All cell lines were cultured at 37°C under 5% CO_2_ in their respective culture media. Wild-type (WT) and Cav1*^-/-^* mouse lung endothelial cell lines (MLECs) [a gift from Radu.V. Stan] were cultured in Endothelial Cell Growth Medium-2 (EGM^TM^-2) Bulletkit^TM^ (Lonza, cat. #CC-3162) composed of EBM-2 Basal Medium (cat. #CC-3156) and supplemented with EGM-2 SingleQuots^TM^ (cat. #CC-4176) containing hydrocortisone, hFGF-B, VEGF, R3-IGF-1, ascorbic acid, hEGF, GA-1000 (Gentamicin, Amphotericin-B) and heparin along with 10% FBS (Gibco, Life Technologies). All Murine Embryonic Fibroblast cell lines used in this study (MEF WT, MEF Cavin1^-/-^ and MEF Cavin1^-/-^ expressing low/medium/high levels of Cav1) [a gift from Miguel del Pozo] were cultured in Dulbecco’s modified Eagle medium (DMEM) GlutaMAX^TM^ (Gibco, Life Technologies) supplemented with 10% FBS (Gibco, Life Technologies), 100 U/ml penicillin-streptomycin, 1 mM sodium pyruvate and 15 mM HEPES.

Plasmids were transfected either by electroporation using Ingenio® electroporation solution (Mirus Bio LLC) or by lipofection using Lipofectamine LTX with Plus reagent (Invitrogen, Life Technologies), Lipofectamine 3000 (Invitrogen, Life Technologies) or HiPerFect transfection reagents (Qiagen Inc.) following the manufacturer’s instructions. Electroporation of cells was performed with a pulse of 220 V and 975 µF using a Gene Pulser II module (Bio-Rad). siRNA transfections were performed using the HiPerFect kit and cells were incubated for 3 days before further experimentation. Depletion efficiency was assessed by immunoblotting.

Unless otherwise stated, stimulation with IFN-α or IFN-β (recombinant mouse IFN-α; BioLegend cat. #752806, mouse IFN-β; tebubio cat. #12400-1) was performed at a concentration of 1000 U/ml and 500 U/ml respectively. For cell stretching experiments, cells were grown on a rectangular PDMS sheet (thickness ∼100 μm, dimensions ∼12 × 7 mm) coated with fibronectin and stretched uni-axially using a custom-built device equipped with a motorized linear actuator (PI, Karlsruhe, Germany) and a temperature controller. Cells were pre-stretched by 25% for 2 minutes and stretch was maintained during IFN-α stimulation. For hypo-osmotic shock (HYPO), cells were subjected to culture media diluted in water and processed for subsequent experiments. Unless otherwise stated, cells were subjected to 30 mOsm hypo-osmotic shock (10% culture media and 90% water) for a duration of 5 minutes. Recovery (REC) of cells were performed by first subjecting the cells to a 5-min hypo-osmotic shock and immediately replenishing with normal culture media (300 mOsm). Unless otherwise stated, recovery (REC) is performed for a duration of 5 minutes post hypo-osmotic shock.

### Plasmids and antibodies

Cav1-GFP and cavin-1-mEGFP were a gift from Ari Helenius (Addgene plasmid # 14433; https://n2t.net/addgene:14433; RRID: Addgene_14433 and Addgene plasmid # 27709; http://n2t.net/addgene:27709; RRID: Addgene_27709 respectively). Cav1-mRFP and Cav1F92A/V94A-mRFP plasmids are described elsewhere ^38^. Cav1-W98-mRFP was generated using site-directed mutagenesis from Cav1-mRFP. mEos3.2-Caveolin-C-10 was a gift from Michael Davidson (Addgene plasmid # 57447; http://n2t.net/addgene:57447; RRID: Addgene 57447). mEos2-CAAX was generated by amplifying the coding DNA sequence of the corresponding protein by PCR and inserted into the pcDNAm-FRT-PC-mEos2 blue at Fse1/Asc1 sites. The fidelity of all constructs was verified by sequencing.

The following primary antibodies were used: mouse anti-α-tubulin (Sigma-Aldrich, clone B512, Cat. #T5168, 1:1000 for WB); mouse anti-STAT3 (Cell signaling, clone 124H6, Cat. #9139, 1:1000 for WB); rabbit anti-pSTAT3 (Tyr705) (Cell signaling technologies, clone D3A7, Cat. #9145, 1:1000 for WB, 1:100 for IF); rabbit anti-STAT1 (Cell signaling technologies, Cat. #9172, 1:1000 for WB); mouse anti-pSTAT1 (Tyr701) (Cell signaling technologies, clone 58D6, Cat. #9167, 1:1000 for WB, 1:100 for IF); mouse anti-Cav1 (BD Transduction, Cat. #610407, 1:1000 for WB); rabbit anti-Cav1 (Cell Signaling Technologies Cat. #3238S, 1:1000 for WB, 2-5 µg/condition for IP, 1:50 for dSTORM, 1:150 for IF); mouse anti-PTRF (BD Transduction Cat. #611258, 1:1000 for WB); rabbit anti-PTRF (Cat. #ab48824, Abcam – discontinued, 1:1000 for WB, 1:50 for dSTORM, 1:150 for IF); rabbit anti-JAK1 (Cell signaling technologies, Cat. #3332S, 1:2000 for WB); mouse anti-JAK1 (Santa Cruz Biotechnology, Cat. #sc-1677, 1:50 for STORM).

The following secondary antibodies were used: Donkey anti-mouse-HRP (Jackson ImmunoResearch, cat. #715-035-151) and Donkey anti-rabbit-HRP (Jackson ImmunoResearch, cat. #711-035-152) were used at a dilution of 1:5,000 for WB; Donkey anti-mouse-AF647 (Jackson ImmunoResearch, cat. #715-606-150) and Donkey anti-rabbit-CF680 (Biotium, cat. #20820) were used at a dilution of 1:200 and 1:400 respectively, for STORM imaging. Donkey anti-mouse DyLight 800 (cat. #SA5-10172); Donkey anti-goat DyLight 800 (cat. #SA5-10044).

### CSD mimicking peptides

CSD mimicking peptides were synthetized from Biomatik. Control peptide sequence: HHHHHH-RQIKIWFQNRRMKWKKWGIDKASFTTFTVTKYWFRY; CavTratin sequence: HHHHHH-QIKIWFQNRRMKWKKDGIWKASFTTFTVTKY; CavNoxin sequence: HHHHH H-RQIKIWFQNRRMKWKKDGIWKASFAAATVTKWYFYR. Cells were treated for 6 hours with 1 μM CSD mimicking peptide resuspended in EGM-2 culture medium.

### In vitro Kinase activity measurement

In-vitro kinase assay was performed using purified JAK1 (ProQinase 1480-0000-1 JAK1 aa583-1154), RBER-IRStide (ProQinase 0863-0000-1). Kinase reaction was performed in Kinase reaction buffer ([ATP] 100 μM, RBER-IRStide 80 μg/ml, DMSO according to peptide concentration) at 30 °C for 1 hr. Measurement of ADP production was performed using Promega ADP-Glo™ Kinase Assay. Luminescence measurement was performed using BMG Labtech FLUOstar Omega plate reader.

### Immunoblotting

Cells were lysed in sample buffer (62.5 mM Tris–HCl, pH 6.0, 2% (vol/vol) SDS, 10% (vol/vol) glycerol, 40 mM dithiothreitol and 0.03% (wt/vol) phenol red). The lysates were analysed by SDS–PAGE on 4–20% mini-PROTEAN TGX or TGX stain-free precast protein gels (Bio-Rad), and immunoblotted with the indicated primary and secondary antibodies that were either horseradish peroxidase-conjugated or fluorescently labelled. The chemiluminescence signal was revealed using Pierce ECL western blotting, SuperSignal west dura extended duration or SuperSignal west femto (Thermo Scientific Life Technologies) substrate. Acquisition and quantification were performed using a ChemiDoc MP imaging system (Bio-Rad). For STAT1, the phosphorylated and total protein levels were assayed on the same blot with the primary antibodies mouse anti-pSTAT1 and rabbit anti-STAT1, and visualized using fluorescence and luminescence, respectively. The ratio of phosphorylated-to-total protein was determined for each time point.

### Immunofluorescence

For immunofluorescence analysis, cells were either cultured on coverslips or PDMS as per the experimental procedure, treated as described earlier and then fixed with ice-cold methanol for 15 min at -20 °C. After washing with 0.2% (wt/vol) BSA in PBS, the cells were subsequently incubated with the indicated primary antibody and fluorescence-conjugated secondary antibody for 1h at room temperature. The coverslips were mounted in Fluoromount-G mounting medium (eBioscience) with 2 µg/ml 4,6-diamidino-2-phenylindole (Sigma-Aldrich) to counterstain nuclei. Images were acquired on a Leica DM 6000B inverted wide-field microscope equipped with a HCX PL Apo ×63, 1.40 numerical aperture (NA) oil immersion objective and an electron-multiplying charge-coupled device (CCD) camera (Photometrics CoolSMAP HQ). Nuclear translocation of pSTAT1/pSTAT3 was quantified using a homemade plugin in the ImageJ software (NIH) by calculating the nuclear-to-cytoplasmic ratio of the pSTAT1/pSTAT3 signal (nuclei masks were realized with 4,6-diamidino-2-phenylindole staining).

### Co-immunoprecipitation

Cells were lysed in 1% NP-40 in ice-cold TNE (10 mM Tris–HCl pH 7.5, 150 mM NaCl and 0.5 mM EDTA) with protease inhibitor cocktail (Roche) for 30 min. For conventional co-immunoprecipitation, cleared lysates (16,000 g for 10 min at 4 °C) were incubated overnight with 1 µg/ml of the indicated antibody at 4 °C, with rotation, followed by incubation for 1 h with 25 µl protein A/G magnetic beads (Thermo Scientific) in the case of endogenous proteins. For tagged proteins, 25 µl of GFP-Trap or RFP-Trap beads (Chromotek) were used. After three washes in TNE, the immunoprecipitated beads were eluted following the manufacturer’s instructions. Magnetic crosslink co-immunoprecipitation (Pierce™ Crosslink Magnetic IP/Co-IP Kit, cat. #88805) was performed as per the manunfacturer’s instruction. Briefly, the desired antibody was covalently crosslinked to protein A/G magnetic beads using the DSS (disuccinimidyl suberate) crosslinker and subsequently incubated overnight with pre-cleared lysates at 4 °C. Incubated beads were then eluted and the immunoprecipitates were analyzed by immunoblotting.

### dSTORM sample preparation

MLEC WT cells grown on high resolution #1.5 glass coverslips (THOR labs) were washed three times with PHEM solution (60 mM PIPES, 25 mM HEPES, 5 mM EGTA and 2 mM Mg acetate adjusted to pH 6.9 with 1 M KOH) and fixed for 20 min in 4% PFA. They were then washed 3 times in PBS (137 mM NaCl, 2.7 mM KCl, 8 mM Na_2_HPO_4_, and 2 mM KH_2_PO_4_). Up to this fixation step, all chemical reagents were pre- warmed at 37°C. The cells were then quenched for auto-fluorescence from PFA in 50mM NH_4_Cl for 20 min at RT. The cells were washed in PBS three times before being blocked and permeabilized in blocking buffer (1X PBS / 1% BSA / 0.1% Saponin) for 1 hr at RT. Fixed cells were incubated for 1 hr at 37°C with the respective primary antibodies diluted in blocking buffer and washed three times with PBS. This was followed by 1 hr incubation at 37°C with corresponding secondary antibodies diluted in blocking buffer and washed three times with PBS. After immunolabelling, a post-fixation step was performed using PBS with 3.6% formaldehyde for 15 min. The cells were washed in PBS three times and then reduced for 10 min with 50 mM NH_4_Cl (Sigma Aldrich, 254134), followed by three additional washes in PBS.

### Dual color dSTORM imagihng

Fluorophores Alexa-Fluor^TM^ 647 (AF647) and CF680 photo switch under reducing and oxygen-free buffer conditions, making them suitable for dSTORM single molecule imaging, which enables the localization of the emitters with sub-diffraction localization precision ^128^. Thanks to their close spectral proximity, AF647 was excited and acquired simultaneously with CF680 in the same dSTORM buffer (Abbelight^TM^ SMART-Kit) using a 640 nm laser (Oxxius), and their respective signals discriminated after single molecule localization using a spectral demixing strategy ^129^. To implement spectral demixing dSTORM of JAK1-A647 and Cav1-CF680, we used a dual-view Abbelight^TM^ SAFe360, equipped with two Hamamatsu Fusion sCMOS cameras and mounted on an Olympus Ix83 inverted microscope with a 100X 1.5 NA TIRF objective. The SAFe360 uses astigmatic PSF engineering to extract the axial position and achieves quasi-isotropic 3D localization precision, and a long-pass dichroic mirror to split fluorescence from single emitters on the two cameras. Samples were illuminated in HILO at 80% of max laser power and imaged at 50 ms exposure time for 100000 frames. Single molecule localization, drift correction, spectral demixing and data visualization were performed using Abbelight^TM^ NEO software.

### 3D SMLM network analysis

A total of 30 cells (15 cells each for ISO and HYPO conditions) stained for Cav1-CF680 and JAK1-AF647 were imaged using the Abbelight SAFe 360 microscope utilizing the spectral demixing technique. Cav1 localizations were processed, and the resulting clusters were classified using the 3D SMLM network analysis pipeline described in ^40^. The datasets were analyzed using the 3D SMLM network analysis based on the following parameters: merging threshold = 19 nm (to correct multiple blinking of single fluorophores); proximity threshold = 80 nm (for network construction); alpha = 2 (for noise filtering); bandwidth = 120 nm (for segmenting the localization into blobs/clusters using mean shift algorithm). Following this, 28 features/descriptors (shape, topology, network, size, hollowness, etc.) were extracted for every segmented blob/cluster. The clusters/blobs feature of every condition were then grouped into four groups using the x-means algorithm. The biological names for the groups i.e., caveolae and S2, S1A, and S1B scaffolds, were obtained by comparing the group centers with the Cav1 groups that were obtained previously in ^40^ and were assigned based on the best match (highest similarity).

### Spatial pattern and interaction analysis

The interaction analysis was performed using the MosaicIA plugin ^74^ for Fiji by loading the 3D co-ordinates of JAK1 and the center of mass of the corresponding Cav1 clusters identified using the 3D SMLM network analysis pipeline (namely caveolae, S2, S1B and S1A scaffolds) from a total of 30 cells (15 cells each for ISO and HYPO condition) and selecting 2 random ROIs from each cell. The workflow for interaction analysis described in ^74^ was followed. Briefly, we compute the cross–nearest-neighbor distance distribution between JAK1 and Cav1 localizations and fit a pairwise Gibbs interaction model that infers the interaction potential’s strength (ε) and range (σ) that best explain the observed pattern after context correction (controls for density and mask geometry). In this framework, positive ε indicates spatial attraction (co-organization) and negative ε indicates repulsion relative to the null. The following parameters were used for computing distance distributions: Grid spacing = 0.2 (this value was chosen by sequentially reducing the grid spacing until the q(d) does not significantly change); Kernel wt(q) = 0.001; Kernel wt(p) was used as suggested by the software. To determine the best parametric potential for the dataset, a non-parametric potential was first used to estimate the shape of interaction. The various available parametric potentials were then tested to determine the one that best fits the estimated shape of interaction. The Linear L1 potential resulted in the best fit and hence was used for subsequent datasets. The strength of interaction was plotted for each Cav1 cluster type under ISO and HYPO conditions.

### Co-localization/nearest localization using distance to centroid

The open-source software Point Cloud Analyst (PoCA) [https://github.com/flevet/ PoCA] was used to determine the distance to centroids between objects of interest (i.e. between JAK1 and the various classes of Cav1 clusters) within a defined ROI. In brief, the 3D coordinates of JAK1 and Cav1 clusters (namely caveolae, S2, S1B and S1A scaffolds) identified using the 3D SMLM network analysis pipeline were loaded individually and the corresponding voronoi diagram or tessellation was generated using an in-built algorithm. For creating objects from the voronoi tessellations, a cut-off threshold of 75 and 100 with a minimum number of localizations of 2 and 11 was used for JAK1 and Cav1 respectively. The centroids of the resulting objects were extracted following which the centroids of JAK1 and the various Cav1 cluster objects were superimposed in pairs (eg: JAK1-Caveolae, JAK1-S2, JAK1-S1B and JAK1-S1A). The distance to centroids for each pair was calculated within several defined ROIs using the Cav1 cluster centroids as the reference i.e. the distance computation was done between JAK1 objects (centroids) and the outline of the reference (Cav1 objects).

### Micromechanical stretching device compatible with super-resolution microscopy

The stretching device compatible with super-resolution microscopy (used for DNA-PAINT and single particle tracking in the current manuscript) has been described previously ^73,130^. Briefly, a plasma-cleaned polydimethylsiloxane (PDMS) sheet (10 μm; Sylgard 184, DE9330, Samaro) was deposited on a plasma-cleaned glass coverslip, lubricated by a thin layer of low-viscous glycerol (glycerol for fluorescence microscopy; CAS 56-81-5; Merck; 1040950250), and reinforced by a thicker elastomer frame (40 µm, PF film X0; 1.5 mil; Gel-Pak). The 40 μm elastomer frame was pre-cut to the size of the glass coverslip with a squared (3 mm × 3 mm) observation chamber using a Graphtec cutting plotter (Graphtec Craft ROBO pro; CE5000-40-CRP). Uniaxial stretch was applied using a milled (Charlyrobot, Mecanumic) poly (methyl methacrylate) (PMMA) device consisting of a fixed holding arm and a mobile arm, positioned on opposite sides of the observation chamber on the elastomer frame. The mobile arm was connected to a mechanical motor (MTS-65, 52 mm; Linear stage with stepper motor; 0.1 μm resolution; PI). PDMS substrate was coated with human fibronectin (10 μg ml^−1^) for 90 min at 37 °C. Then, after electroporation with Cav1-GFP, MLEC Cav1*^-/-^* cells were plated on the stretching device and spread overnight at 37 °C.

### DNA-PAINT acquisition and analysis

The stretching device was mounted on an inverted motorized microscope (Nikon Ti) equipped with a CFI Apochromat TIRF 100× oil, NA 1.49 objective and a perfect focus system (PFS-2). First, live stretching onto the microscope was performed at 37 °C. To calibrate the strain on each stretching device, we adsorbed 0.1-μm fluorescent beads (TetraSpeck Microspheres; 0.1 μm; Thermo Fisher Scientific; T7279) on the stretching chamber and used a small stretch (2% to 3%) prior to the stretching protocol. Strain is calculated using distances of same beads before and after stretch: Strain=(L_stretch_-L_0_)/L_0_, where L_0_ is the beads distance before stretch and L_stretch_ is the distance for same beads after stretch. Then, a uniaxial cyclic stretch (stretch: 30% strain, 0.5 Hz, 30 min) was applied, followed by rapid cell fixation in 4% paraformaldehyde in PBS buffer for 15 min. Then, cells were quenched with glycine (150 mM) for 20 min, and blocked for 90 min with 3% BSA and 0.2% Triton X-100 in PBS, then incubated with GFP nanobody conjugated with a DNA-strand P1 for 4 hours (5’ Nanobody GFP - TTA TAC ATC TA 3’), followed by a second fixation with 4% paraformaldehyde and 0.2% glutaraldehyde in PBS for 20 min.

DNA-PAINT acquisitions ^131^ after live stretching and fixation were performed at 25°C in stretching devices on the same microscope. Prior to DNA-PAINT, we acquired low-resolution images of Cav1-GFP. Then, super-resolution imaging was performed thanks to the perfect focus system, allowing long acquisition in TIRF illumination mode, required for single-molecule localization microscopy including DNA-PAINT. To register super-resolution intensity images, we adsorbed 90-nm gold nanoparticles (Cytodiagnostics) on the stretching chamber that were imaged during the entirety of the DNA-PAINT acquisitions. Cy3B-labeled DNA imager strands (5’ CTA GAT GTA T - Cy3b 3’) were added to the stretching chamber at variable concentrations (0.2 to 1 nM), and visualized with a 561-nm laser (Coherent Obis FP series lasers) with 20 mW power at the sample plane. Fluorescence was collected by the combination of a dichroic filter and emission filters (dichroic, Di01-R561; emission, FF01-617/73; Semrock) and a sensitive scientific complementary metal-oxide semiconductor (ORCA-Flash4.0, Hammamatsu). Cav1-GFP was imaged using a conventional GFP filter cube (excitation, FF01-472/30; dichroic, FF-495Di02; emission, FF02-520/28; Semrock). The acquisitions were steered by MetaMorph software (Molecular Devices) in streaming mode at 100 ms for 90 000 frames. DNA-PAINT image reconstruction and drift correction were carried out using the Picasso software ^131^. The spatial resolution of DNA-PAINT acquisitions obtained on stretching device was ∼15.7 ± 1.5 nm (full width at half-maximum (FWHM)). SR-Tesseler ^132^ was used to segment DNA-PAINT super-resolution images, to obtain the size distribution of Cav1-GFP clusters without stretching, or after cyclic stretching. Single-molecule localizations were used to compute Voronoï diagram. Clusters were segmented using average density factor of 20. To filter background noise of DNA-PAINT acquisitions, often corresponding to unspecific binding of some DNA imager strands to the cover-glass, we set a threshold for the number of localizations (50), enabling to select only genuine signals associated with binding to DNA docking strands. To analyze the smallest Cav1-GFP structures detected using DNA-PAINT, we manually selected isolated clusters that exhibited DNA-PAINT signals throughout the entire acquisition period. The size of these clusters was measured using SR-Tesseler ^131^.

### sptPALM sample preparation

For both MLEC and MEF cells, transient transfections of plasmids were performed 1 day before experiments using the Nucleofactor™ transfection kit for MEF-1 and Nucleofactor™ IIb device (Amaxa™, Lonza). For MLECs, cells were detached with accutase solution (Sigma Aldrich, cat. #SLBT9789). The accutase was inactivated using EGM-2 medium, and the cells were washed and suspended in EGM-2 medium. Cells were then seeded overnight in EGM-2 medium on nitric-acid cleaned glass coverslips. The next day, EGM-2 medium was rinsed once with PBS and left for experiment in serum-free Ringer medium (150 mM NaCl, 5 mM KCl, 2 mM CaCl2, 2 mM MgCl2, 10 mM HEPES, pH=7.4) supplemented with 11 mM glucose. For MEFs, cells were detached with 0.05% trypsin, 0.02% EDTA solution (Gibco Cat. #25300054). The trypsin was inactivated using soybean trypsin inhibitor (1 mg/ml in DMEM, Sigma), and the cells were washed and suspended in serum-free Ringer medium (150 mM NaCl, 5 mM KCl, 2 mM CaCl2, 2 mM MgCl2, 10 mM HEPES, pH=7.4) supplemented with 11 mM glucose.

Cells were then seeded on human fibronectin-coated surface (fibronectin: 10 μg/ml, Roche). MEF PTRF/Cavin1^-/-^ cell line was a gift from Miguel del Pozo (Spanish National Centre for Cardiovascular Research, Spain) and are described elsewhere ^42^. Absence of mycoplasma contamination was assessed using the MycoAlert detection kit (Lonza Cat. No. LT07-318). For sptPALM, 120,000 MLEC or 50,000 MEF cells were seeded on #1.5H glass coverslips (Marienfeld). When mentioned, hypo-osmotic shock was induced by replacing the observation medium (Ringer+glucose) with a Ringer+glucose solution diluted 10 times with MQ grade deionized water, at least 5 minutes before acquisition.

### sptPALM optical setup and image acquisition

All acquisitions were steered by MetaMorph software (Molecular Devices) with an inverted motorized microscope (Nikon Ti) equipped with a temperature control system (The Cube, The Box, Life Imaging Services), a Nikon CFI Apo TIRF 100X oil, NA 1.49 objective and a perfect focus system, allowing long acquisition in TIRF illumination mode. The coverslip was mounted in a Ludin chamber (Life Imaging Services) before acquisition. For photoactivation localization microscopy, cells expressing mEos3.2 tagged constructs were photoactivated using a 405 nm laser (Omicron) and the resulting photoconverted single molecule fluorescence was excited with a 561 nm laser (Cobolt Jive™). Both lasers illuminated the sample simultaneously. Their respective power was adjusted to keep the number of the stochastically activated molecules constant and well separated during the acquisition. Fluorescence was collected by the combination of a dichroic and emission filters (D101-R561 and F39-617 respectively; Chroma) and a sensitive EMCCD (electron-multiplying charge-coupled device, Evolve, Photometric). The acquisition was performed in streaming mode at 50 Hz. Cav1-GFP, Cavin1-meGFP was imaged using a conventional GFP filter cube (ET470/40, T495LPXR, ET525/50, Chroma). Using this filter cube does not allow spectral separation of the unconverted pool of mEos3.2 from the GFP fluorescent signal. For this reason, in the case of MEF Cavin1^-/-^ cell line, we were able to detect caveolin1-based structures (scaffolds) with the unconverted pool of mEos3.2-caveolin1 (whose emission spectra is similar to the one of GFP).

### Single molecule segmentation and tracking

A typical sptPALM experiment leads to a set of at least 4000 images per cell, analyzed to extract molecule localization and dynamics. Single molecule fluorescent spots were localized and tracked over time using a combination of wavelet segmentation and simulated annealing algorithms ^133,134,135^. Under the experimental conditions described above, the resolution of the system was quantified to 59 nm (Full Width at Half Maximum, FWHM). This spatial resolution depends on the image signal to noise ratio and the segmentation algorithm ^136^ and was determined using fixed mEos3.2 samples. We analyzed 130 2D distributions of single molecule positions belonging to long trajectories (>50 frames) by bi-dimensional Gaussian fitting, the resolution being determined as 2.3 *s_xy_*, where *s_xy_* is the pointing accuracy.

For the trajectory analysis, cell contours were identified manually from Cav1-GFP or unconverted pool of mEos3.2-caveolin1 images. We analyzed trajectories lasting at least 260 ms (≥13 points) with a custom Matlab routine analyzing the mean squared displacement (MSD), which describes the diffusion properties of a molecule, computed as (Eq. 1):

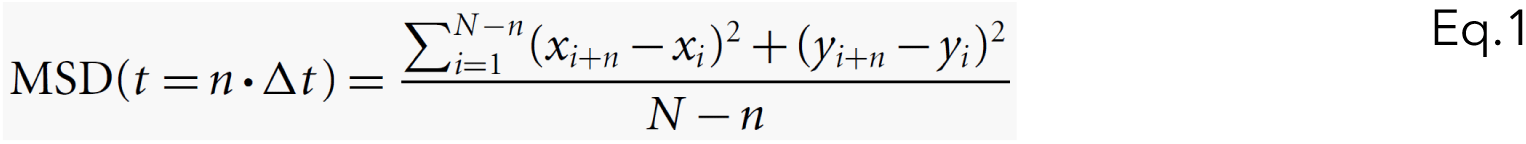

where *xi* and *yi* are the coordinates of the label position at time *I x Δt*. We defined the measured diffusion coefficient *D* as the slope of the affine regression line fitted to the *n*=1 to 4 values of the MSD (*n x Δt)*. The MSD was computed then fitted on a duration equal to 80% (minimum of 10 points, 200 ms) of the whole stretch by (Eq. 2):

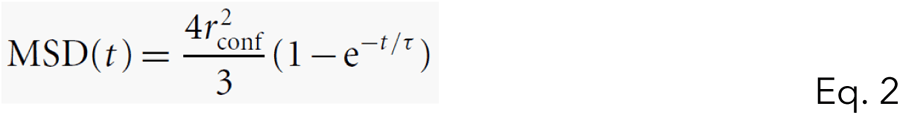

where *r*_conf_ is the measured confinement radius and τ the time constant τ= (*r*_conf_² / 3*D*_conf_). To reduce the inaccuracy of the MSD fit due to down sampling for larger time intervals, we used a weighted fit. Trajectories were sorted in 3 groups: immobile, confined and diffusive. Immobile trajectories were defined as trajectories with *D*<0.011 μm^2^.s^-1^, corresponding to molecules which explored an area inferior to the one defined by the image spatial resolution ∼(0.05 μm)² during the time used to fit the initial slope of the MSD ^60^ (4 points, 80 ms): *D*_threshold_=(0.059 μm)²/(4x4x0.02 s)∼0.011 μm^2^.s^-1^. To separate confined and diffusive trajectories, we used the time constant calculated τ for each trajectory. Confined and diffusive events were defined as trajectories with a time constant inferior and superior to half the time interval used to compute the MSD (100 ms) respectively.

### Molecular Modelling and Simulation

Structural predictions were performed using AlphaFold3 ^81^ through the AlphaFold Server (https://alphafoldserver.com). Protein sequences were retrieved from UniProt [The UniProt Consortium, 2025] (Q03135, CAV1_HUMAN; P23458, JAK1_HUMAN). Structural models were analysed using UCSF ChimeraX 1.11 ^137^.

The AF3-predicted structure of the JAK1 kinase domain in complex with Cav1 [80–110] was further investigated by molecular dynamics simulations in explicit solvent. The initial structure was built and refined using successive steps of energy minimization and restrained molecular dynamics with CHARMM v49b1 ^138^, using the CHARMM36m force field ^139^. The system was solvated in a TIP3P water orthorhombic box using the solvate module of VMD ^140^ with a minimum distance of 12 Å between the protein/peptide and the box boundaries in all directions. The system was subsequently neutralized and supplemented with Na⁺ and Cl⁻ ions to reach an ionic strength of 150 mM. Energy minimization was performed for 10,000 steps under positional restraints to maintain the structure close to the initial conformation. Several minimization cycles were applied, during which the force constant on atomic positions was gradually reduced from 10 kcal·mol⁻¹·Å⁻² to 0. This step was followed by a 2 ns molecular dynamics equilibration step in the NVT ensemble. Production simulations were then carried out in the NPT ensemble for 0.7 µs. To account for stochastic effects, three MD trajectories were run with different initial velocities. Temperature was maintained at 310 K and pressure at 1 atm using Langevin dynamics. During production, a time step of 2 fs was used, and multiple time stepping was applied using the rRESPA algorithm, with long-range electrostatic interactions computed every 4 fs using the Particle Mesh Ewald (PME) method with a real-space grid spacing of 1 Å. Periodic boundary conditions were applied. A cutoff distance of 12 Å was used for short-range electrostatic and van der Waals interactions. Molecular dynamics simulations were performed using NAMD 3.0.1 (CUDA version) ^141^. Calculations were carried out on A100 GPU resources provided by GENCI at IDRIS@CNRS (Jean Zay supercomputer). Trajectory analyses were conducted using CHARMM v49b1. Molecular dynamics (MD) trajectories were analysed using ChimeraX 1.11 and VMD. Figures were generated with ChimeraX 1.11 and Gnuplot 6.0 (http://www.gnuplot.info (2023).

## Extended Data Figure Legends

**Extended Data Figure 1. (a)** Representative immunoblots showing levels of Tyr 705 pSTAT3 (top) and STAT3 (bottom) in MLEC WT and MLEC Cav1^-/-^ under iso-osmotic and hypo-osmotic conditions in the presence or absence of IFN-α activation as indicated. Quantification is based on the signal intensity ratio of pSTAT3 relative to the total protein obtained from stain-free blot. **(b)** Representative immunoblots of lysates of MLEC WT expressing RFP, Cav1(WT)-RFP or Cav1(F92A/V94A)-RFP and probed for pSTAT3, STAT3 and α-tubulin at basal state. Statistics denote the ratio of STAT3/α-tubulin expression level in comparison to cells expressing RFP only. All panels exhibit representative immunoblots for N=3 independent experiments. Data shown are mean values ± SD. Statistic significances were obtained using **(a,b)** multiple-comparison ANOVA. ****P<0.0001 and ns: not significant.

**Extended Data Figure 2. (a)** Distribution of the diffusion coefficient D computed from the trajectories of mEos2-CAAX in MLEC WT cells in ISO (dark orange) and HYPO (light orange) conditions, shown in a logarithmic scale. The grey area including D values inferior to 0.011 µm².s-1 corresponds to immobilized proteins. Fraction of mEos3.2-CAAX undergoing free diffusion, confined diffusion or immobilization in MLEC WT cells under ISO and HYPO conditions. Data from various defined ROIs in ISO (n=9 cells) and HYPO (n=11 cells) pooled from N=2 independent experiments. Statistical significance was obtained using multiple-comparison ANOVA. ns: not significant.

**Extended Data Figure 3. (a) (Left)** Representative 2D dSTORM acquisitions of MLEC WT cells stained for Cav1-AF647 and subjected to iso-osmotic (ISO), hypo-osmotic (HYPO) and recovery (REC) conditions. **(Middle)** Line intensity profiles for Cav1 positive structures depicting the changes in caveolae diameter in each condition. **(Right)** Box-and-whisker plots showing the distribution of Cav1 cluster area (nm²) quantified for iso-osmotic (ISO), hypo-osmotic shock (HYPO) and recovery (REC) conditions. Each data point represents one segmented Cav1 cluster (n=73 clusters per condition). Whiskers denote minimum to maximum values. Statistical significance was assessed by ordinary one-way ANOVA followed by Tukey’s multiple-comparisons test; ****P < 0.0001; ns, not significant. **(b)** Representative images of Cav1-CF680 and Cavin1-CF680 acquired through 3D STORM spectral demixing in MLEC WT cells and estimation of caveolar Cav1 and Cavin1 cluster sizes from dSTORM acquisitions in MLEC WT cells at steady state. Bold line denotes the mean value. **(c)** Side-view of 3D point cloud localizations depicting the morphology of a bona fide budded caveolae in MLEC WT cells at steady state. Dotted line represents the potential localization of plasma membrane. **(d)** Representative 3D STORM localizations of Cav1 in MLEC WT cells under hypo-osmotic shock (HYPO) processed and segmented using the 3D SMLM network/graph-based analysis pipeline to classify Cav1 localizations into caveolae and non-caveolar scaffold populations (S2, S1B and S1A). For (**b**) Data shown for (n=40) particles are mean values ± SEM. Statistical significance was obtained using unpaired two-tailed t-test. ns: not significant.

**Extended Data Figure 4. (a)** Representative DNA-PAINT localizations of Cav1 in MLEC WT cells under Control (no stretch) and 30 min uniaxial stretch. DNA-PAINT coordinate datasets were processed and segmented using the same 3D SMLM network/graph-based analysis pipeline to classify Cav1 localizations into caveolae and non-caveolar scaffold populations (S2, S1B and S1A). **(b)** Corresponding quantification of the density of Cav1 clusters per unit area for each class are depicted. **(c)** Visualization and colocalization of 3D point cloud localizations of Cav1-CF680 and JAK1-AF647 imaged using spectral demixing in MLEC WT cells at steady state depicting ‘visual’ (qualitative) co-localization between clusters of Cav1 and JAK1. **(d)** Extracted centroids for the corresponding pairs of objects from representative 3D STORM image of localizations of JAK1 and the various Cav1 populations described in Fig. 4e, superimposed with the JAK1 localizations within a defined ROI under iso-osmotic (left) and hypo-osmotic shock (right) conditions. For (**b**), data shown for control (n=3) and stretched (n=4) cells are mean ± SD. Statistical significance was obtained using two-tailed paired Mann-Whitney t-test. ns, not significant.

**Extended Data Figure 5. (a) Left:** Representative blots of immunoprecipitation from N=3 independent experiments of Cav1 in MLEC WT cells overexpressing ALFA-tagged constructs of either JAK1 WT, JAK1 D^1072–1154^ or JAK1 D^1057–1154^. **Right:** Schematic representation of the ALFA-tagged JAK1 constructs. (**b–d**) Root-mean-square deviation (RMSD) of Cα atoms along MD trajectories 1 (b), 2 (c), and 3 (d) for the JAK1[855–1154] kinase domain (green), the Cav1[80–110] peptide (orange), and the Cav1[90–98] segment (blue). **(e)** Root-mean-square fluctuation (RMSF) of Cα atoms for JAK1[855–1154]. **(f)** RMSF of Cα atoms for the Cav1[80–110] peptide. In **(e)** and **(f)**, dashed lines indicate RMSF values of 1.0 Å and 2.0 Å. **(g–i)** Minimum distances along MD trajectories 1 (green), 2 (blue), and 3 (orange) between the side chain of Cav1 residue V94 and JAK1 residues V1045 **(g)**, Y1048 **(h)**, and L1053 **(i)**. **(j–m)** Minimum distances between the side chain of Cav1 residue F92 and JAK1 residues V1045 **(j)**, F1046 **(k)**, T1100 **(l)**, and V1101 **(m)**. **(n)** Distance between the Nε2 atom of Cav1 residue W98 and the backbone carbonyl oxygen of JAK1 residue Q1055 along the MD trajectories. **(o)** Cartoon representation of the AF3-refined JAK1[855–1154]–Cav1[80–110] complex. JAK1 is shown in grey, except for segment 1033–1055 (cyan), segment 1056–1070 (red), and segment 1071–1154 (orange). Cav1[80–110] is shown in green, and the side chains of residues T90, F92, V94, and W98 are displayed in ball-and-stick representation.

**Extended Data Figure 6. (a)** Heat map of activation of signaling effectors in MLEC WT and Cav1^-/-^ cells treated or not with Type-I IFN under conditions of resting or uniaxial stretch. **(b)** p44/42 MAPK and AKT phosphorylation ratios (phospho-protein/total protein) under conditions of resting or uniaxial stretch in MLEC WT and Cav1^-/-^ cells are depicted. **(c)** STAT1 and STAT3 phosphorylation ratios in MLEC WT and Cav1^-/-^ cells stimulated or not with Type-I IFN. **(d)** STAT1 and STAT3 phosphorylation ratios under conditions of resting or uniaxial stretch upon Type-I IFN stimulation in MLEC WT and Cav1^-/-^ cells. Data shown as box-and-whisker plot are median, min and max values in log2 form from N=3 biological replicates.

**Extended Data Figure 7. (a)** Representative blots of co-immunoprecipitation of endogenous eNOS and PTP1B with Cav1 in MLEC WT cells under ISO, HYPO and REC conditions. **(b)** Representative blots of co-immunoprecipitation of endogenous eNOS and PTP1B with Cav1 in MLEC WT cells under ISO condition and increasing percentages of hypo-osmotic (HYPO) shock. **(c)** Representative blots of co-immunoprecipitation of endogenous JAK1, eNOS and PTP1B with Cav1 in MLEC WT cells under ISO conditions and increasing exposure times to 90% hypo-osmotic (HYPO) shock. For **(a-c),** quantification is based on the signal intensity ratio relative to the corresponding immunoprecipitated Cav1 protein levels. **(d) Left:** MLEC WT and MLEC Cav1 -/- cells were subjected to iso-osmotic (ISO), hypo-osmotic shock (HYPO) or recovery (REC; return to iso-osmotic medium after HYPO) and all conditions were stimulated with H₂O₂ prior to lysis. Representative immunoblots show p-eNOS (Ser1177), total eNOS and Cav1. **Right:** quantification of p-eNOS (Ser1177) intensity normalized to total eNOS. Data shown for (N=3) independent replicates are mean ± SEM. Statistical significance was obtained using multiple-comparison ANOVA. *P<0.05; **P<0.01; ****P<0.0001 and ns: not significant.

**Extended Data Figure 8. (a)** Co-immunoprecipitation of endogenous PTEN with Cav1 and corresponding Ser473 pAKT levels in MLEC WT cells under ISO, HYPO and REC conditions. Quantification for PTEN is based on the signal intensity ratio of PTEN relative to the intensity of immuno-precipitated Cav1 while quantification for Ser473 pAKT is based on the signal intensity ratio of Ser473 pAKT relative to the intensity of total protein obtained from strain-free blot. **(b)** Representative immunoblots of co-immunoprecipitation of Cav1-mRFP with endogenous TYK2, PTEN, and AKT in MLEC Cav1^-/-^ cells expressing either Cav1(WT)-mRFP, Cav1(F92A/V94A)-mRFP, or Cav1(W98A)-mRFP under ISO conditions. The immunoblot for RFP is the same as that shown in Fig. 6h. **(c)** Representative immunoblot for immunoprecipitation of Cav1 and probing for co-immunoprecipitation of β1-integrin under ISO, HYPO and REC conditions. All panels exhibit representative immunoblots for N=3 independent experiments. Data shown are mean values ± SD. Statistics were performed using repeated measures multiple-comparison one-way ANOVA. ****P<0.0001.

## Legends for Supplementary Videos

**Supplementary Video 1**: Dynamics of mEos3.2-Cav1 under steady state (ISO) conditions.

Visualization of mEos3.2-Cav1 trajectories in MEF WT cells at steady state (ISO) reveals that most of the Cav1 molecules are immobile or confined, presumably within *bona fide* caveolae. Images were acquired at an interval of 400 ms for 79.6 s. Scale bar: 3 µm

**Supplementary Video 2**: Dynamics of mEos3.2-Cav1 under conditions of mechanical stress induced by hypo-osmotic shock (HYPO).

Visualization of mEos3.2-Cav1 trajectories in MEF WT cells under conditions of increased membrane tension induced by hypo-osmotic shock reveals that there is an exponential increase in the number of highly diffusive Cav1 molecules. Images were acquired at an interval of 400 ms for 79.6 s. Scale bar: 3 µm

## Notes

### Competing Interest Statement

The authors have declared no competing interest.

### Summary of Updates

This revised version includes substantial updates made in response to peer-review feedback. We performed multiple new experiments to address key reviewer comments and to strengthen the mechanistic support for our conclusions. In parallel, we reorganized and rearranged the figure set (including updated panels and new data) to improve clarity and logical progression. The manuscript text has been extensively rewritten and streamlined to provide a more coherent narrative, clearer rationale for experimental design, and improved integration of results with the discussion. Overall, the revision enhances the rigor, completeness, and readability of the work while preserving the central findings of the original preprint.

## References

1. Lamaze, C., Tardif, N., Dewulf, M., Vassilopoulos, S., and Blouin, C.M. (2017). The caveolae dress code: structure and signaling. Curr Opin Cell Biol 47, 117–125. 10.1016/j.ceb.2017.02.014.

2. Parton, R.G., Del Pozo, M.A., Vassilopoulos, S., Nabi, I.R., Le Lay, S., Lundmark, R., Kenworthy, A.K., Camus, A., Blouin, C.M., Sessa, W.C., and Lamaze, C. (2020). Caveolae: The FAQs. Traffic 21, 181–185. 10.1111/tra.12689.

3. Palade, G.E. (1953). The fine structure of blood capillaries. J. Appl. Phys. 24, 1424.

4. Yamada, E. (1955). The fine structure of the gall bladder epithelium of the mouse. J Biophys Biochem Cytol 1, 445–458.

5. Rothberg, K.G., Heuser, J.E., Donzell, W.C., Ying, Y.S., Glenney, J.R., and Anderson, R.G. (1992). Caveolin, a protein component of caveolae membrane coats. Cell 68, 673–682.

6. Way, M., and Parton, R.G. (1996). M-caveolin, a muscle-specific caveolin-related protein. FEBS Lett 378, 108–112.

7. Scherer, P.E., Okamoto, T., Chun, M., Nishimoto, I., Lodish, H.F., and Lisanti, M.P. (1996). Identification, sequence, and expression of caveolin-2 defines a caveolin gene family. Proc Natl Acad Sci U S A 93, 131–135.

8. Aboulaich, N., Vainonen, J.P., Stralfors, P., and Vener, A.V. (2004). Vectorial proteomics reveal targeting, phosphorylation and specific fragmentation of polymerase I and transcript release factor (PTRF) at the surface of caveolae in human adipocytes. Biochem J 383, 237–248.

9. Hill, M.M., Bastiani, M., Luetterforst, R., Kirkham, M., Kirkham, A., Nixon, S.J., Walser, P., Abankwa, D., Oorschot, V.M., Martin, S., et al. (2008). PTRF-Cavin, a conserved cytoplasmic protein required for caveola formation and function. Cell 132, 113–124.

10. Liu, L., and Pilch, P.F. (2008). A critical role of cavin (polymerase I and transcript release factor) in caveolae formation and organization. J Biol Chem 283, 4314–4322.

11. Bastiani, M., Liu, L., Hill, M.M., Jedrychowski, M.P., Nixon, S.J., Lo, H.P., Abankwa, D., Luetterforst, R., Fernandez-Rojo, M., Breen, M.R., et al. (2009). MURC/Cavin-4 and cavin family members form tissue-specific caveolar complexes. J Cell Biol 185, 1259–1273.

12. Sverdlov, M., Shinin, V., Place, A.T., Castellon, M., and Minshall, R.D. (2009). Filamin A regulates caveolae internalization and trafficking in endothelial cells. Mol Biol Cell 20, 4531–4540. 10.1091/mbc.e08-10-0997.

13. Senju, Y., Itoh, Y., Takano, K., Hamada, S., and Suetsugu, S. (2011). Essential role of PACSIN2/syndapin-II in caveolae membrane sculpting. J Cell Sci 124, 2032–2040.

14. Hansen, C.G., Howard, G., and Nichols, B.J. (2011). Pacsin 2 is recruited to caveolae and functions in caveolar biogenesis. J Cell Sci 124, 2777–2785.

15. Moren, B., Shah, C., Howes, M.T., Schieber, N.L., McMahon, H.T., Parton, R.G., Daumke, O., and Lundmark, R. (2012). EHD2 regulates caveolar dynamics via ATP-driven targeting and oligomerization. Mol Biol Cell 23, 1316–1329.

16. Koch, D., Westermann, M., Kessels, M.M., and Qualmann, B. (2012). Ultrastructural freeze-fracture immunolabeling identifies plasma membrane-localized syndapin II as a crucial factor in shaping caveolae. Histochem Cell Biol 138, 215–230. 10.1007/s00418-012-0945-0.

17. Porta, J.C., Han, B., Gulsevin, A., Chung, J.M., Peskova, Y., Connolly, S., McHaourab, H.S., Meiler, J., Karakas, E., Kenworthy, A.K., and Ohi, M.D. (2022). Molecular architecture of the human caveolin-1 complex. Sci Adv 8, eabn7232. 10.1126/sciadv.abn7232.

18. Patel, H.H., Murray, F., and Insel, P.A. (2008). Caveolae as organizers of pharmacologically relevant signal transduction molecules. Annu Rev Pharmacol Toxicol 48, 359–391.

19. Lajoie, P., and Nabi, I.R. (2010). Lipid rafts, caveolae, and their endocytosis. International review of cell and molecular biology 282, 135–163. 10.1016/s1937-6448(10)82003-9.

20. Pilch, P.F., Meshulam, T., Ding, S., and Liu, L. (2011). Caveolae and lipid trafficking in adipocytes. Clin Lipidol 6, 49–58.

21. Parton, R.G., McMahon, K.A., and Wu, Y. (2020). Caveolae: Formation, dynamics, and function. Curr Opin Cell Biol 65, 8–16. 10.1016/j.ceb.2020.02.001.

22. Lamaze, C., Blouin, C.M., and Sens, P. (accepted in principle October 23^rd^). Caveolae Mechanics in Cellular Functions and Disease. Nature Reviews Molecular Cell Biology.

23. Minshall, R.D., Sessa, W.C., Stan, R.V., Anderson, R.G., and Malik, A.B. (2003). Caveolin regulation of endothelial function. Am J Physiol Lung Cell Mol Physiol 285, L1179–1183. 10.1152/ajplung.00242.2003.

24. Williams, J.J., and Palmer, T.M. (2014). Cavin-1: caveolae-dependent signalling and cardiovascular disease. Biochem Soc Trans 42, 284–288. 10.1042/bst20130270.

25. Singh, V., and Lamaze, C. (2020). Membrane tension buffering by caveolae: a role in cancer? Cancer metastasis reviews 2, 505–517. 10.1007/s10555-020-09899-2.

26. Liu, L. (2020). Lessons from cavin-1 deficiency. Biochem Soc Trans 48, 147–154. 10.1042/bst20190380.

27. Boscher, C., and Nabi, I.R. (2012). Caveolin-1: role in cell signaling. Advances in experimental medicine and biology 729, 29–50. 10.1007/978-1-4614-1222-9_3.

28. Fridolfsson, H.N., Roth, D.M., Insel, P.A., and Patel, H.H. (2014). Regulation of intracellular signaling and function by caveolin. Faseb j 28, 3823–3831. 10.1096/fj.14-252320.

29. Couet, J., Sargiacomo, M., and Lisanti, M.P. (1997). Interaction of a receptor tyrosine kinase, EGF-R, with caveolins. Caveolin binding negatively regulates tyrosine and serine/threonine kinase activities. J Biol Chem 272, 30429–30438. 10.1074/jbc.272.48.30429.

30. Nystrom, F.H., Chen, H., Cong, L.N., Li, Y., and Quon, M.J. (1999). Caveolin-1 interacts with the insulin receptor and can differentially modulate insulin signaling in transfected Cos-7 cells and rat adipose cells. Mol Endocrinol 13, 2013–2024. 10.1210/mend.13.12.0392.

31. Bernatchez, P.N., Bauer, P.M., Yu, J., Prendergast, J.S., He, P., and Sessa, W.C. (2005). Dissecting the molecular control of endothelial NO synthase by caveolin-1 using cell-permeable peptides. Proc Natl Acad Sci U S A 102, 761–766.

32. Grande-Garcia, A., Echarri, A., de Rooij, J., Alderson, N.B., Waterman-Storer, C.M., Valdivielso, J.M., and del Pozo, M.A. (2007). Caveolin-1 regulates cell polarization and directional migration through Src kinase and Rho GTPases. J Cell Biol 177, 683–694. 10.1083/jcb.200701006.

33. Ariotti, N., Fernandez-Rojo, M.A., Zhou, Y., Hill, M.M., Rodkey, T.L., Inder, K.L., Tanner, L.B., Wenk, M.R., Hancock, J.F., and Parton, R.G. (2014). Caveolae regulate the nanoscale organization of the plasma membrane to remotely control Ras signaling. J Cell Biol 204, 777–792. 10.1083/jcb.201307055.

34. Shvets, E., Bitsikas, V., Howard, G., Hansen, C.G., and Nichols, B.J. (2015). Dynamic caveolae exclude bulk membrane proteins and are required for sorting of excess glycosphingolipids. Nature communications 6, 6867. 10.1038/ncomms7867.

35. Lajoie, P., Partridge, E.A., Guay, G., Goetz, J.G., Pawling, J., Lagana, A., Joshi, B., Dennis, J.W., and Nabi, I.R. (2007). Plasma membrane domain organization regulates EGFR signaling in tumor cells. J Cell Biol 179, 341–356.

36. Joshi, B., Strugnell, S.S., Goetz, J.G., Kojic, L.D., Cox, M.E., Griffith, O.L., Chan, S.K., Jones, S.J., Leung, S.P., Masoudi, H., et al. (2008). Phosphorylated caveolin-1 regulates Rho/ROCK-dependent focal adhesion dynamics and tumor cell migration and invasion. Cancer Res 68, 8210–8220. 10.1158/0008-5472.Can-08-0343.

37. Moon, H., Lee, C.S., Inder, K.L., Sharma, S., Choi, E., Black, D.M., KA, L.C., Winterford, C., Coward, J.I., Ling, M.T., et al. (2014). PTRF/cavin-1 neutralizes non-caveolar caveolin-1 microdomains in prostate cancer. Oncogene 33, 3561–3570. 10.1038/onc.2013.315.

38. Meng, F., Saxena, S., Liu, Y., Joshi, B., Wong, T.H., Shankar, J., Foster, L.J., Bernatchez, P., and Nabi, I.R. (2017). The phospho-caveolin-1 scaffolding domain dampens force fluctuations in focal adhesions and promotes cancer cell migration. Mol Biol Cell 28, 2190–2201. 10.1091/mbc.E17-05-0278.

39. Lajoie, P., Goetz, J.G., Dennis, J.W., and Nabi, I.R. (2009). Lattices, rafts, and scaffolds: domain regulation of receptor signaling at the plasma membrane. J Cell Biol 185, 381–385.

40. Khater, I.M., Meng, F., Wong, T.H., Nabi, I.R., and Hamarneh, G. (2018). Super Resolution Network Analysis Defines the Molecular Architecture of Caveolae and Caveolin-1 Scaffolds. Scientific reports 8, 9009. 10.1038/s41598-018-27216-4.

41. Khater, I.M., Liu, Q., Chou, K.C., Hamarneh, G., and Nabi, I.R. (2019). Super-resolution modularity analysis shows polyhedral caveolin-1 oligomers combine to form scaffolds and caveolae. Scientific reports 9, 9888. 10.1038/s41598-019-46174-z.

42. Lolo, F.N., Walani, N., Seemann, E., Zalvidea, D., Pavón, D.M., Cojoc, G., Zamai, M., Viaris de Lesegno, C., Martínez de Benito, F., Sánchez-Álvarez, M., et al. (2022). Caveolin-1 dolines form a distinct and rapid caveolae-independent mechanoadaptation system. Nat Cell Biol 25. 10.1038/s41556-022-01034-3.

43. Sinha, B., Koster, D., Ruez, R., Gonnord, P., Bastiani, M., Abankwa, D., Stan, R.V., Butler-Browne, G., Vedie, B., Johannes, L., et al. (2011). Cells respond to mechanical stress by rapid disassembly of caveolae. Cell 144, 402–413.

44. Cheng, J.P., Mendoza-Topaz, C., Howard, G., Chadwick, J., Shvets, E., Cowburn, A.S., Dunmore, B.J., Crosby, A., Morrell, N.W., and Nichols, B.J. (2015). Caveolae protect endothelial cells from membrane rupture during increased cardiac output. J Cell Biol 211, 53–61. 10.1083/jcb.201504042.

45. Lo, H.P., Nixon, S.J., Hall, T.E., Cowling, B.S., Ferguson, C., Morgan, G.P., Schieber, N.L., Fernandez-Rojo, M.A., Bastiani, M., Floetenmeyer, M., et al. (2015). The caveolin-cavin system plays a conserved and critical role in mechanoprotection of skeletal muscle. J Cell Biol 210, 833–849. 10.1083/jcb.201501046.

46. Elliott, M.H., Ashpole, N.E., Gu, X., Herrnberger, L., McClellan, M.E., Griffith, G.L., Reagan, A.M., Boyce, T.M., Tanito, M., Tamm, E.R., and Stamer, W.D. (2016). Caveolin-1 modulates intraocular pressure: implications for caveolae mechanoprotection in glaucoma. Scientific reports 6, 37127. 10.1038/srep37127.

47. Lim, Y.W., Lo, H.P., Ferguson, C., Martel, N., Giacomotto, J., Gomez, G.A., Yap, A.S., Hall, T.E., and Parton, R.G. (2017). Caveolae Protect Notochord Cells against Catastrophic Mechanical Failure during Development. Curr Biol 27, 1968–1981.e1967. 10.1016/j.cub.2017.05.067.

48. Garcia, J., Bagwell, J., Njaine, B., Norman, J., Levic, D.S., Wopat, S., Miller, S.E., Liu, X., Locasale, J.W., Stainier, D.Y.R., and Bagnat, M. (2017). Sheath Cell Invasion and Trans-differentiation Repair Mechanical Damage Caused by Loss of Caveolae in the Zebrafish Notochord. Curr Biol 27, 1982–1989.e1983. 10.1016/j.cub.2017.05.035.

49. Nassoy, P., and Lamaze, C. (2012). Stressing caveolae new role in cell mechanics. Trends Cell Biol 22, 381–389. S0962-8924(12)00074-8 [pii]10.1016/j.tcb.2012.04.007 [doi].

50. Parton, R.G., and Del Pozo, M.A. (2013). Caveolae as plasma membrane sensors, protectors and organizers. Nat Rev Mol Cell Biol 14, 98–112.

51. Cheng, J.P., and Nichols, B.J. (2016). Caveolae: One Function or Many? Trends Cell Biol 26, 177-189. 10.1016/j.tcb.2015.10.010.

52. Del Pozo, M.A., Lolo, F.N., and Echarri, A. (2021). Caveolae: Mechanosensing and mechanotransduction devices linking membrane trafficking to mechanoadaptation. Curr Opin Cell Biol 68, 113–123. 10.1016/j.ceb.2020.10.008.

53. Dewulf, M., Koster, D.V., Sinha, B., Viaris de Lesegno, C., Chambon, V., Bigot, A., Bensalah, M., Negroni, E., Tardif, N., Podkalicka, J., et al. (2019). Dystrophy-associated caveolin-3 mutations reveal that caveolae couple IL6/STAT3 signaling with mechanosensing in human muscle cells. Nature communications 10, 1974. 10.1038/s41467-019-09405-5.

54. Schreiber, G., and Piehler, J. (2015). The molecular basis for functional plasticity in type I interferon signaling. Trends in immunology 36, 139–149. 10.1016/j.it.2015.01.002.

55. de Weerd, N.A., Kurowska, A.K., Mendoza, J.L., and Schreiber, G. (2024). Structure-function of type I and III interferons. Current opinion in immunology 86, 102413. 10.1016/j.coi.2024.102413.

56. Geletu, M., Taha, Z., Arulanandam, R., Mohan, R., Assi, H.H., Castro, M.G., Nabi, I.R., Gunning, P.T., and Raptis, L.P. (2019). Effect of Caveolin-1 upon Stat3-ptyr705 levels in breast and lung carcinoma cells. Biochemistry and cell biology = Biochimie et biologie cellulaire. 10.1139/bcb-2018-0367.

57. Geletu, M., Mohan, R., Arulanandam, R., Berger-Becvar, A., Nabi, I.R., Gunning, P.T., and Raptis, L. (2018). Reciprocal regulation of the Cadherin-11/Stat3 axis by caveolin-1 in mouse fibroblasts and lung carcinoma cells. Biochimica et biophysica acta. Molecular cell research 1865, 794–802. 10.1016/j.bbamcr.2018.02.004.

58. Gambin, Y., Ariotti, N., McMahon, K.A., Bastiani, M., Sierecki, E., Kovtun, O., Polinkovsky, M.E., Magenau, A., Jung, W., Okano, S., et al. (2013). Single-molecule analysis reveals self assembly and nanoscale segregation of two distinct cavin subcomplexes on caveolae. eLife 3, e01434. 10.7554/eLife.01434.

59. Torrino, S., Shen, W.W., Blouin, C.M., Mani, S.K., Viaris de Lesegno, C., Bost, P., Grassart, A., Koster, D., Valades-Cruz, C.A., Chambon, V., et al. (2018). EHD2 is a mechanotransducer connecting caveolae dynamics with gene transcription. J Cell Biol 217, 4092–4105. 10.1083/jcb.201801122.

60. Rossier, O., Octeau, V., Sibarita, J.B., Leduc, C., Tessier, B., Nair, D., Gatterdam, V., Destaing, O., Albiges-Rizo, C., Tampe, R., et al. (2012). Integrins beta1 and beta3 exhibit distinct dynamic nanoscale organizations inside focal adhesions. Nat Cell Biol 14, 1057–1067. 10.1038/ncb2588.

61. Mehidi, A., Kage, F., Karatas, Z., Cercy, M., Schaks, M., Polesskaya, A., Sainlos, M., Gautreau, A.M., Rossier, O., Rottner, K., and Giannone, G. (2021). Forces generated by lamellipodial actin filament elongation regulate the WAVE complex during cell migration. Nat Cell Biol 23, 1148–1162. 10.1038/s41556-021-00786-8.

62. Orré, T., Joly, A., Karatas, Z., Kastberger, B., Cabriel, C., Böttcher, R.T., Lévêque-Fort, S., Sibarita, J.B., Fässler, R., Wehrle-Haller, B., et al. (2021). Molecular motion and tridimensional nanoscale localization of kindlin control integrin activation in focal adhesions. Nature communications 12, 3104. 10.1038/s41467-021-23372-w.

63. Pol, A., Morales-Paytuví, F., Bosch, M., and Parton, R.G. (2020). Non-caveolar caveolins - duties outside the caves. J Cell Sci 133. 10.1242/jcs.241562.

64. Orre, T., Mehidi, A., Massou, S., Rossier, O., and Giannone, G. (2018). Using Single-Protein Tracking to Study Cell Migration. Methods in molecular biology (Clifton, N.J.) 1749, 291–311. 10.1007/978-1-4939-7701-7_21.

65. Ouyang, W., Aristov, A., Lelek, M., Hao, X., and Zimmer, C. (2018). Deep learning massively accelerates super-resolution localization microscopy. Nature biotechnology 36, 460–468. 10.1038/nbt.4106.

66. Schermelleh, L., Ferrand, A., Huser, T., Eggeling, C., Sauer, M., Biehlmaier, O., and Drummen, G.P.C. (2019). Super-resolution microscopy demystified. Nat Cell Biol 21, 72–84. 10.1038/s41556-018-0251-8.

67. Khater, I.M., Nabi, I.R., and Hamarneh, G. (2020). A Review of Super-Resolution Single-Molecule Localization Microscopy Cluster Analysis and Quantification Methods. Patterns (N Y) 1, 100038. 10.1016/j.patter.2020.100038.

68. Möckl, L., Roy, A.R., and Moerner, W.E. (2020). Deep learning in single-molecule microscopy: fundamentals, caveats, and recent developments [Invited]. Biomedical optics express 11, 1633–1661. 10.1364/boe.386361.

69. Nunes Vicente, F., Chen, T., Rossier, O., and Giannone, G. (2023). Novel imaging methods and force probes for molecular mechanobiology of cytoskeleton and adhesion. Trends Cell Biol 33, 204–220. 10.1016/j.tcb.2022.07.008.

70. Monier, S., Parton, R.G., Vogel, F., Behlke, J., Henske, A., and Kurzchalia, T.V. (1995). VIP21- caveolin, a membrane protein constituent of the caveolar coat, oligomerizes in vivo and in vitro. Mol Biol Cell 6, 911–927.

71. Hayer, A., Stoeber, M., Bissig, C., and Helenius, A. (2009). Biogenesis of Caveolae: Stepwise Assembly of Large Caveolin and Cavin Complexes. Traffic 3, 3.

72. Pelkmans, L., and Zerial, M. (2005). Kinase-regulated quantal assemblies and kiss-and-run recycling of caveolae. Nature 436, 128–133.

73. Massou, S., Nunes Vicente, F., Wetzel, F., Mehidi, A., Strehle, D., Leduc, C., Voituriez, R., Rossier, O., Nassoy, P., and Giannone, G. (2020). Cell stretching is amplified by active actin remodelling to deform and recruit proteins in mechanosensitive structures. Nature Cell Biology. 10.1038/s41556-020-0548-2.

74. Shivanandan, A., Radenovic, A., and Sbalzarini, I.F. (2013). MosaicIA: an ImageJ/Fiji plugin for spatial pattern and interaction analysis. BMC Bioinformatics 14, 349. 10.1186/1471-2105-14-349.

75. Li, S., Couet, J., and Lisanti, M.P. (1996). Src tyrosine kinases, Galpha subunits, and H-Ras share a common membrane-anchored scaffolding protein, caveolin. Caveolin binding negatively regulates the auto-activation of Src tyrosine kinases. J Biol Chem 271, 29182–29190.

76. García-Cardeña, G., Martasek, P., Masters, B.S., Skidd, P.M., Couet, J., Li, S., Lisanti, M.P., and Sessa, W.C. (1997). Dissecting the interaction between nitric oxide synthase (NOS) and caveolin. Functional significance of the nos caveolin binding domain in vivo. J Biol Chem 272, 25437–25440. 10.1074/jbc.272.41.25437.

77. Bernatchez, P., Sharma, A., Bauer, P.M., Marin, E., and Sessa, W.C. (2011). A noninhibitory mutant of the caveolin-1 scaffolding domain enhances eNOS-derived NO synthesis and vasodilation in mice. J Clin Invest 121, 3747–3755.

78. Couet, J., Li, S., Okamoto, T., Ikezu, T., and Lisanti, M.P. (1997). Identification of peptide and protein ligands for the caveolin-scaffolding domain. Implications for the interaction of caveolin with caveolae-associated proteins. J Biol Chem 272, 6525–6533. 10.1074/jbc.272.10.6525.

79. Oka, N., Yamamoto, M., Schwencke, C., Kawabe, J., Ebina, T., Ohno, S., Couet, J., Lisanti, M.P., and Ishikawa, Y. (1997). Caveolin interaction with protein kinase C. Isoenzyme-dependent regulation of kinase activity by the caveolin scaffolding domain peptide. J Biol Chem 272, 33416–33421. 10.1074/jbc.272.52.33416.

80. Collins, B.M., Davis, M.J., Hancock, J.F., and Parton, R.G. (2012). Structure-based reassessment of the caveolin signaling model: do caveolae regulate signaling through caveolin-protein interactions? Dev Cell 23, 11–20.

81. Abramson, J., Adler, J., Dunger, J., Evans, R., Green, T., Pritzel, A., Ronneberger, O., Willmore, L., Ballard, A.J., Bambrick, J., et al. (2024). Accurate structure prediction of biomolecular interactions with AlphaFold 3. Nature 630, 493–500. 10.1038/s41586-024-07487-w.

82. Doktorova, M., Daum, S., Reagle, T.R., Cannon, H.I., Ebenhan, J., Neudorf, S., Han, B., Sharma, S., Kasson, P., Levental, K.R., et al. (2025). Caveolin assemblies displace one bilayer leaflet to organize and bend membranes. Proc Natl Acad Sci U S A 122, e2417024122. 10.1073/pnas.2417024122.

83. Boellner, S., and Becker, K.F. (2015). Reverse Phase Protein Arrays-Quantitative Assessment of Multiple Biomarkers in Biopsies for Clinical Use. Microarrays (Basel) 4, 98–114. 10.3390/microarrays4020098.

84. Lee, H., Xie, L., Luo, Y., Lee, S.Y., Lawrence, D.S., Wang, X.B., Sotgia, F., Lisanti, M.P., and Zhang, Z.Y. (2006). Identification of phosphocaveolin-1 as a novel protein tyrosine phosphatase 1B substrate. Biochemistry 45, 234–240. 10.1021/bi051560j.

85. Saeedi Saravi, S.S., Eroglu, E., Waldeck-Weiermair, M., Sorrentino, A., Steinhorn, B., Belousov, V., and Michel, T. (2020). Differential endothelial signaling responses elicited by chemogenetic H(2)O(2) synthesis. Redox Biol 36, 101605. 10.1016/j.redox.2020.101605.

86. Caselli, A., Mazzinghi, B., Camici, G., Manao, G., and Ramponi, G. (2002). Some protein tyrosine phosphatases target in part to lipid rafts and interact with caveolin-1. Biochem Biophys Res Commun 296, 692–697. 10.1016/s0006-291x(02)00928-2.

87. Lolo, F.-N., Pavón, D.M., Grande, A., Elósegui Artola, A., Segatori, V.I., Sánchez, S., Trepat, X., Roca-Cusachs, P., and del Pozo, M.A. (2022). Caveolae couple mechanical stress to integrin recycling and activation. eLife 11, e82348. 10.7554/eLife.82348.

88. Monteiro, P., Remy, D., Lemerle, E., Routet, F., Macé, A.S., Guedj, C., Ladoux, B., Vassilopoulos, S., Lamaze, C., and Chavrier, P. (2023). A mechanosensitive caveolae-invadosome interplay drives matrix remodelling for cancer cell invasion. Nat Cell Biol 25, 1787–1803. 10.1038/s41556-023-01272-z.

89. Sens, P., and Turner, M.S. (2004). Theoretical model for the formation of caveolae and similar membrane invaginations. Biophys J 86, 2049–2057. 10.1016/s0006-3495(04)74266-6.

90. Dulhunty, A.F., and Franzini-Armstrong, C. (1975). The relative contributions of the folds and caveolae to the surface membrane of frog skeletal muscle fibres at different sarcomere lengths. J Physiol 250, 513–539.

91. Prescott, L., and Brightman, M.W. (1976). The sarcolemma of Aplysia smooth muscle in freeze-fracture preparations. Tissue Cell 8, 248–258.

92. Gabella, G., and Blundell, D. (1978). Effect of stretch and contraction on caveolae of smooth muscle cells. Cell and tissue research 190, 255–271.

93. Qifti, A., Balaji, S., and Scarlata, S. (2022). Deformation of caveolae impacts global transcription and translation processes through relocalization of cavin-1. J Biol Chem 298, 102005. 10.1016/j.jbc.2022.102005.

94. McMahon, K.A., Stroud, D.A., Gambin, Y., Tillu, V., Bastiani, M., Sierecki, E., Polinkovsky, M.E., Hall, T.E., Gomez, G.A., Wu, Y., et al. (2021). Cavin3 released from caveolae interacts with BRCA1 to regulate the cellular stress response. eLife 10, e61407. 10.7554/eLife.61407.

95. Byrne, D.P., Dart, C., and Rigden, D.J. (2012). Evaluating caveolin interactions: do proteins interact with the caveolin scaffolding domain through a widespread aromatic residue-rich motif? PLoS One 7, e44879. 10.1371/journal.pone.0044879.

96. Kenworthy, A.K., Han, B., Ariotti, N., and Parton, R.G. (2023). The Role of Membrane Lipids in the Formation and Function of Caveolae. Cold Spring Harbor perspectives in biology. 10.1101/cshperspect.a041413.

97. Han, B., Porta, J.C., Hanks, J.L., Peskova, Y., Binshtein, E., Dryden, K., Claxton, D.P., McHaourab, H.S., Karakas, E., Ohi, M.D., and Kenworthy, A.K. (2020). Structure and assembly of CAV1 8S complexes revealed by single particle electron microscopy. Sci Adv 6. 10.1126/sciadv.abc6185.

98. Lim, J.E., Bernatchez, P., and Nabi, I.R. (2024). Scaffolds and the scaffolding domain: an alternative paradigm for caveolin-1 signaling. Biochem Soc Trans 52, 947–959. 10.1042/bst20231570.

99. Wong, T.H., Khater, I.M., Joshi, B., Shahsavari, M., Hamarneh, G., and Nabi, I.R. (2021). Single molecule network analysis identifies structural changes to caveolae and scaffolds due to mutation of the caveolin-1 scaffolding domain. Scientific reports 11. 10.1038/s41598-021-86770-6.

100. Okada, S., Raja, S.A., Okerblom, J., Boddu, A., Horikawa, Y., Ray, S., Okada, H., Kawamura, I., Murofushi, Y., Murray, F., and Patel, H.H. (2019). Deletion of caveolin scaffolding domain alters cancer cell migration. Cell Cycle 18, 1268–1280. 10.1080/15384101.2019.1618118.

101. Jasmin, J.F., Mercier, I., Sotgia, F., and Lisanti, M.P. (2006). SOCS proteins and caveolin-1 as negative regulators of endocrine signaling. Trends in endocrinology and metabolism: TEM 17, 150–158. 10.1016/j.tem.2006.03.007.

102. Kershaw, N.J., Murphy, J.M., Lucet, I.S., Nicola, N.A., and Babon, J.J. (2013). Regulation of Janus kinases by SOCS proteins. Biochem Soc Trans 41, 1042–1047. 10.1042/bst20130077.

103. Williams, J.J.L., Alotaiq, N., Mullen, W., Burchmore, R., Liu, L., Baillie, G.S., Schaper, F., Pilch, P.F., and Palmer, T.M. (2018). Interaction of suppressor of cytokine signalling 3 with cavin-1 links SOCS3 function and cavin-1 stability. Nature communications 9, 168. 10.1038/s41467-017-02585-y.

104. Caveney, N.A., Saxton, R.A., Waghray, D., Glassman, C.R., Tsutsumi, N., Hubbard, S.R., and Garcia, K.C. (2023). Structural basis of Janus kinase trans-activation. Cell reports 42, 112201. 10.1016/j.celrep.2023.112201.

105. Owen, K.L., Brockwell, N.K., and Parker, B.S. (2019). JAK-STAT Signaling: A Double-Edged Sword of Immune Regulation and Cancer Progression. Cancers 11. 10.3390/cancers11122002.

106. Singh, V., Breton, V., Viaris de Lesegno, C., Macé, A.S., Bun, P., Blouin, C.M., Vishen, A.S., Sens, P., and Lamaze, C. (2025). Spatiotemporal coupling of caveolae mechanosensing and RhoA-GEFs regulates cell polarity and directional migration. Nature communications 17, 398. 10.1038/s41467-025-67090-z.

107. Friedland, J.C., Lee, M.H., and Boettiger, D. (2009). Mechanically activated integrin switch controls alpha5beta1 function. Science 323, 642–644. 10.1126/science.1168441.

108. Giannone, G., Jiang, G., Sutton, D.H., Critchley, D.R., and Sheetz, M.P. (2003). Talin1 is critical for force-dependent reinforcement of initial integrin-cytoskeleton bonds but not tyrosine kinase activation. J Cell Biol 163, 409–419. 10.1083/jcb.200302001.

109. Jiang, G., Giannone, G., Critchley, D.R., Fukumoto, E., and Sheetz, M.P. (2003). Two-piconewton slip bond between fibronectin and the cytoskeleton depends on talin. Nature 424, 334–337. 10.1038/nature01805.

110. Wang, Y., Botvinick, E.L., Zhao, Y., Berns, M.W., Usami, S., Tsien, R.Y., and Chien, S. (2005). Visualizing the mechanical activation of Src. Nature 434, 1040–1045. 10.1038/nature03469.

111. del Rio, A., Perez-Jimenez, R., Liu, R., Roca-Cusachs, P., Fernandez, J.M., and Sheetz, M.P. (2009). Stretching single talin rod molecules activates vinculin binding. Science 323, 638–641. 10.1126/science.1162912.

112. Sawada, Y., Tamada, M., Dubin-Thaler, B.J., Cherniavskaya, O., Sakai, R., Tanaka, S., and Sheetz, M.P. (2006). Force sensing by mechanical extension of the Src family kinase substrate p130Cas. Cell 127, 1015–1026.

113. Gordon, W.R., Zimmerman, B., He, L., Miles, L.J., Huang, J., Tiyanont, K., McArthur, D.G., Aster, J.C., Perrimon, N., Loparo, J.J., and Blacklow, S.C. (2015). Mechanical Allostery: Evidence for a Force Requirement in the Proteolytic Activation of Notch. Dev Cell 33, 729–736. 10.1016/j.devcel.2015.05.004.

114. Iskratsch, T., Wolfenson, H., and Sheetz, M.P. (2014). Appreciating force and shape-the rise of mechanotransduction in cell biology. Nat Rev Mol Cell Biol 15, 825–833. 10.1038/nrm3903.

115. Ladoux, B., and Mège, R.M. (2017). Mechanobiology of collective cell behaviours. Nat Rev Mol Cell Biol 18, 743–757. 10.1038/nrm.2017.98.

116. Kirby, T.J., and Lammerding, J. (2018). Emerging views of the nucleus as a cellular mechanosensor. Nat Cell Biol 20, 373–381. 10.1038/s41556-018-0038-y.

117. del Pozo, M.A., Balasubramanian, N., Alderson, N.B., Kiosses, W.B., Grande-Garcia, A., Anderson, R.G., and Schwartz, M.A. (2005). Phospho-caveolin-1 mediates integrin-regulated membrane domain internalization. Nat Cell Biol 7, 901–908.

118. Osmani, N., Pontabry, J., Comelles, J., Fekonja, N., Goetz, J.G., Riveline, D., Georges-Labouesse, E., and Labouesse, M. (2018). An Arf6- and caveolae-dependent pathway links hemidesmosome remodeling and mechanoresponse. Mol Biol Cell 29, 435–451. 10.1091/mbc.E17-06-0356.

119. Muriel, O., Echarri, A., Hellriegel, C., Pavon, D.M., Beccari, L., and Del Pozo, M.A. (2011). Phosphorylated filamin A regulates actin-linked caveolae dynamics. J Cell Sci 124, 2763–2776.

120. Goetz, J.G., Joshi, B., Lajoie, P., Strugnell, S.S., Scudamore, T., Kojic, L.D., and Nabi, I.R. (2008). Concerted regulation of focal adhesion dynamics by galectin-3 and tyrosine-phosphorylated caveolin-1. J Cell Biol 180, 1261–1275.

121. Gottlieb-Abraham, E., Shvartsman, D.E., Donaldson, J.C., Ehrlich, M., Gutman, O., Martin, G.S., and Henis, Y.I. (2013). Src-mediated caveolin-1 phosphorylation affects the targeting of active Src to specific membrane sites. Mol Biol Cell 24, 3881–3895. 10.1091/mbc.E13-03-0163.

122. Zimnicka, A.M., Husain, Y.S., Shajahan, A.N., Sverdlov, M., Chaga, O., Chen, Z., Toth, P.T., Klomp, J., Karginov, A.V., Tiruppathi, C., et al. (2016). Src-dependent phosphorylation of caveolin-1 Tyr-14 promotes swelling and release of caveolae. Mol Biol Cell 27, 2090–2106. 10.1091/mbc.E15-11-0756.

123. Joshi, B., Bastiani, M., Strugnell, S.S., Boscher, C., Parton, R.G., and Nabi, I.R. (2012). Phosphocaveolin-1 is a mechanotransducer that induces caveola biogenesis via Egr1 transcriptional regulation. J Cell Biol 199, 425–435. 10.1083/jcb.201207089.

124. Sotodosos-Alonso, L., Pulgarín-Alfaro, M., and Del Pozo, M.A. (2023). Caveolae Mechanotransduction at the Interface between Cytoskeleton and Extracellular Matrix. Cells 12. 10.3390/cells12060942.

125. Goetz, J.G., Lajoie, P., Wiseman, S.M., and Nabi, I.R. (2008). Caveolin-1 in tumor progression: the good, the bad and the ugly. Cancer metastasis reviews 27, 715–735.

126. Lamaze, C., and Torrino, S. (2015). Caveolae and Cancer: A New Mechanical Perspective. Biomedical journal 38, 367–379. 10.4103/2319-4170.164229.

127. Simón, L., Campos, A., Leyton, L., and Quest, A.F.G. (2020). Caveolin-1 function at the plasma membrane and in intracellular compartments in cancer. Cancer metastasis reviews 39, 435–453. 10.1007/s10555-020-09890-x.

128. Heilemann, M., van de Linde, S., Schüttpelz, M., Kasper, R., Seefeldt, B., Mukherjee, A., Tinnefeld, P., and Sauer, M. (2008). Subdiffraction-resolution fluorescence imaging with conventional fluorescent probes. Angewandte Chemie (International ed. in English) 47, 6172–6176. 10.1002/anie.200802376.

129. Lampe, A., Tadeus, G., and Schmoranzer, J. (2015). Spectral demixing avoids registration errors and reduces noise in multicolor localization-based super-resolution microscopy. Methods Appl Fluoresc 3, 034006. 10.1088/2050-6120/3/3/034006.

130. Nunes Vicente, F., Lelek, M., Tinevez, J.Y., Tran, Q.D., Pehau-Arnaudet, G., Zimmer, C., Etienne-Manneville, S., Giannone, G., and Leduc, C. (2022). Molecular organization and mechanics of single vimentin filaments revealed by super-resolution imaging. Sci Adv 8, eabm2696. 10.1126/sciadv.abm2696.

131. Schnitzbauer, J., Strauss, M.T., Schlichthaerle, T., Schueder, F., and Jungmann, R. (2017). Super-resolution microscopy with DNA-PAINT. Nature protocols 12, 1198–1228. 10.1038/nprot.2017.024.

132. Levet, F., Hosy, E., Kechkar, A., Butler, C., Beghin, A., Choquet, D., and Sibarita, J.B. (2015). SR-Tesseler: a method to segment and quantify localization-based super-resolution microscopy data. Nat Methods 12, 1065–1071. 10.1038/nmeth.3579.

133. Izeddin, I., Boulanger, J., Racine, V., Specht, C.G., Kechkar, A., Nair, D., Triller, A., Choquet, D., Dahan, M., and Sibarita, J.B. (2012). Wavelet analysis for single molecule localization microscopy. Opt Express 20, 2081–2095. 10.1364/oe.20.002081.

134. Racine, V., Sachse, M., Salamero, J., Fraisier, V., Trubuil, A., and Sibarita, J.B. (2007). Visualization and quantification of vesicle trafficking on a three-dimensional cytoskeleton network in living cells. Journal of microscopy 225, 214–228. 10.1111/j.1365-2818.2007.01723.x.

135. Racine, V., Hertzog, A., Jouanneau, J., Salamero, J., Kervrann, C., and Sibarita, J.B. (2006). Multiple-target tracking of 3D fluorescent objects based on simulated annealing. 6-9 April 2006. pp. 1020–1023.

136. Cheezum, M.K., Walker, W.F., and Guilford, W.H. (2001). Quantitative comparison of algorithms for tracking single fluorescent particles. Biophys J 81, 2378–2388. 10.1016/s0006-3495(01)75884-5.

137. Meng, E.C., Goddard, T.D., Pettersen, E.F., Couch, G.S., Pearson, Z.J., Morris, J.H., and Ferrin, T.E. (2023). UCSF ChimeraX: Tools for structure building and analysis. Protein Sci 32, e4792. 10.1002/pro.4792.

138. Brooks, B.R., Brooks, C.L., 3rd, Mackerell, A.D., Jr., Nilsson, L., Petrella, R.J., Roux, B., Won, Y., Archontis, G., Bartels, C., Boresch, S., et al. (2009). CHARMM: the biomolecular simulation program. J Comput Chem 30, 1545–1614. 10.1002/jcc.21287.

139. Huang, J., Rauscher, S., Nawrocki, G., Ran, T., Feig, M., de Groot, B.L., Grubmüller, H., and MacKerell, A.D., Jr. (2017). CHARMM36m: an improved force field for folded and intrinsically disordered proteins. Nat Methods 14, 71–73. 10.1038/nmeth.4067.

140. Humphrey, W., Dalke, A., and Schulten, K. (1996). VMD: visual molecular dynamics. J Mol Graph 14, 33–38, 27-38. 10.1016/0263-7855(96)00018-5.

141. Phillips, J.C., Braun, R., Wang, W., Gumbart, J., Tajkhorshid, E., Villa, E., Chipot, C., Skeel, R.D., Kalé, L., and Schulten, K. (2005). Scalable molecular dynamics with NAMD. J Comput Chem 26, 1781–1802. 10.1002/jcc.20289.

