## Extended Data Figures for "Remote Control of Cell Signaling through Caveolae Mechanics"

Extended Data figure 1:

a

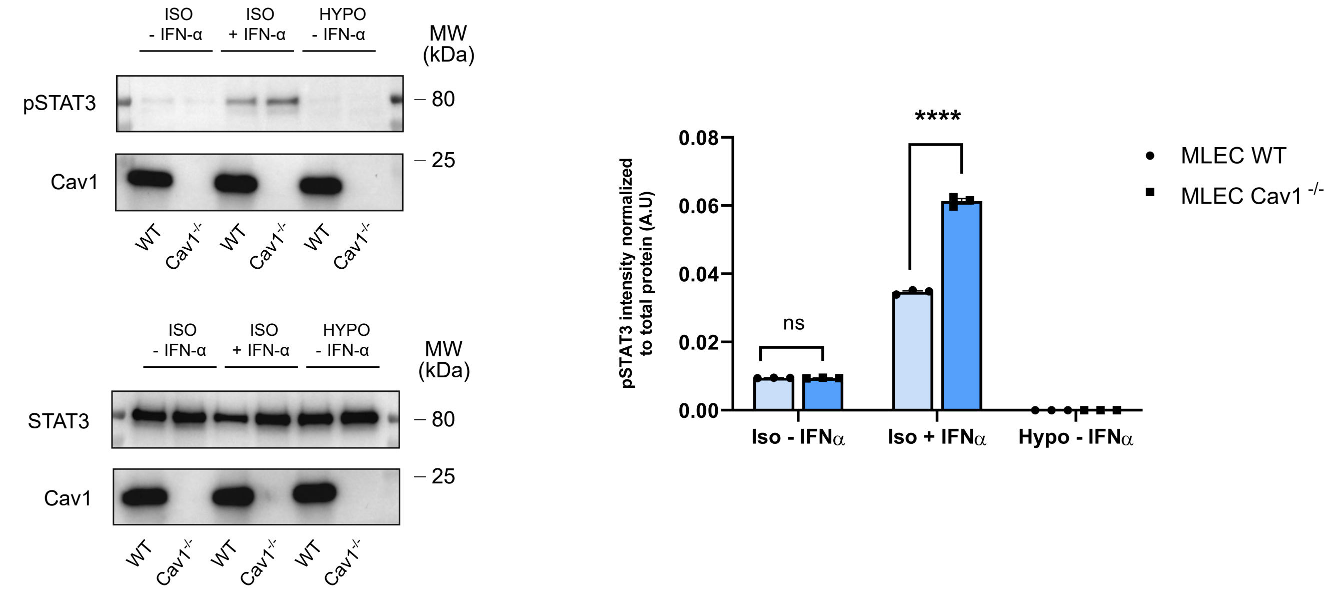


b

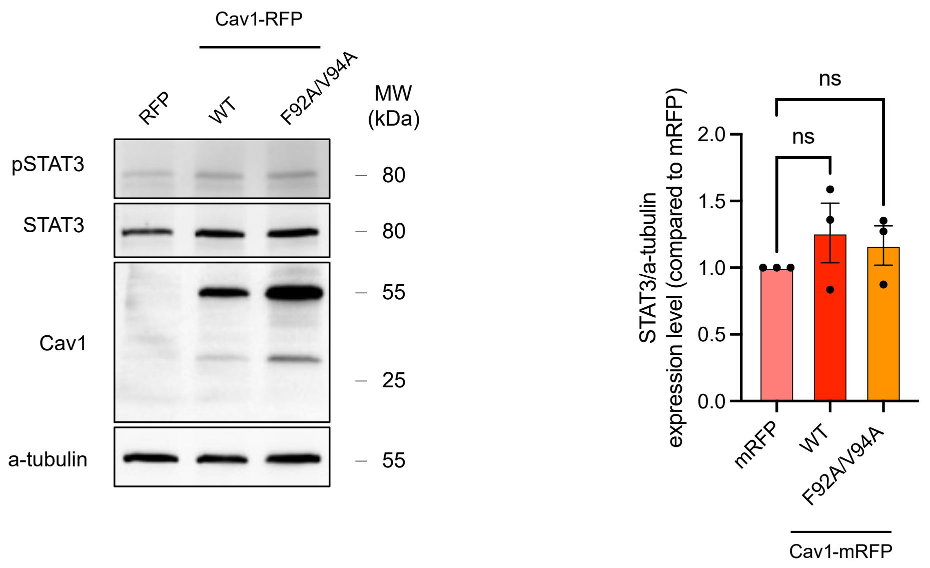


Extended Data figure 2:

a

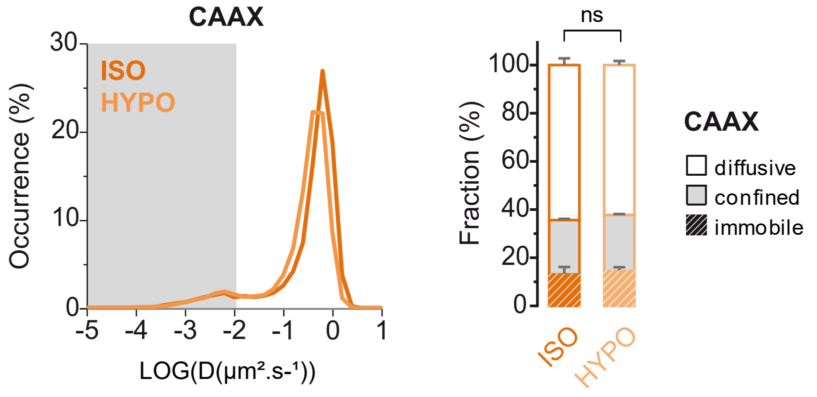

Extended Data figure 3:

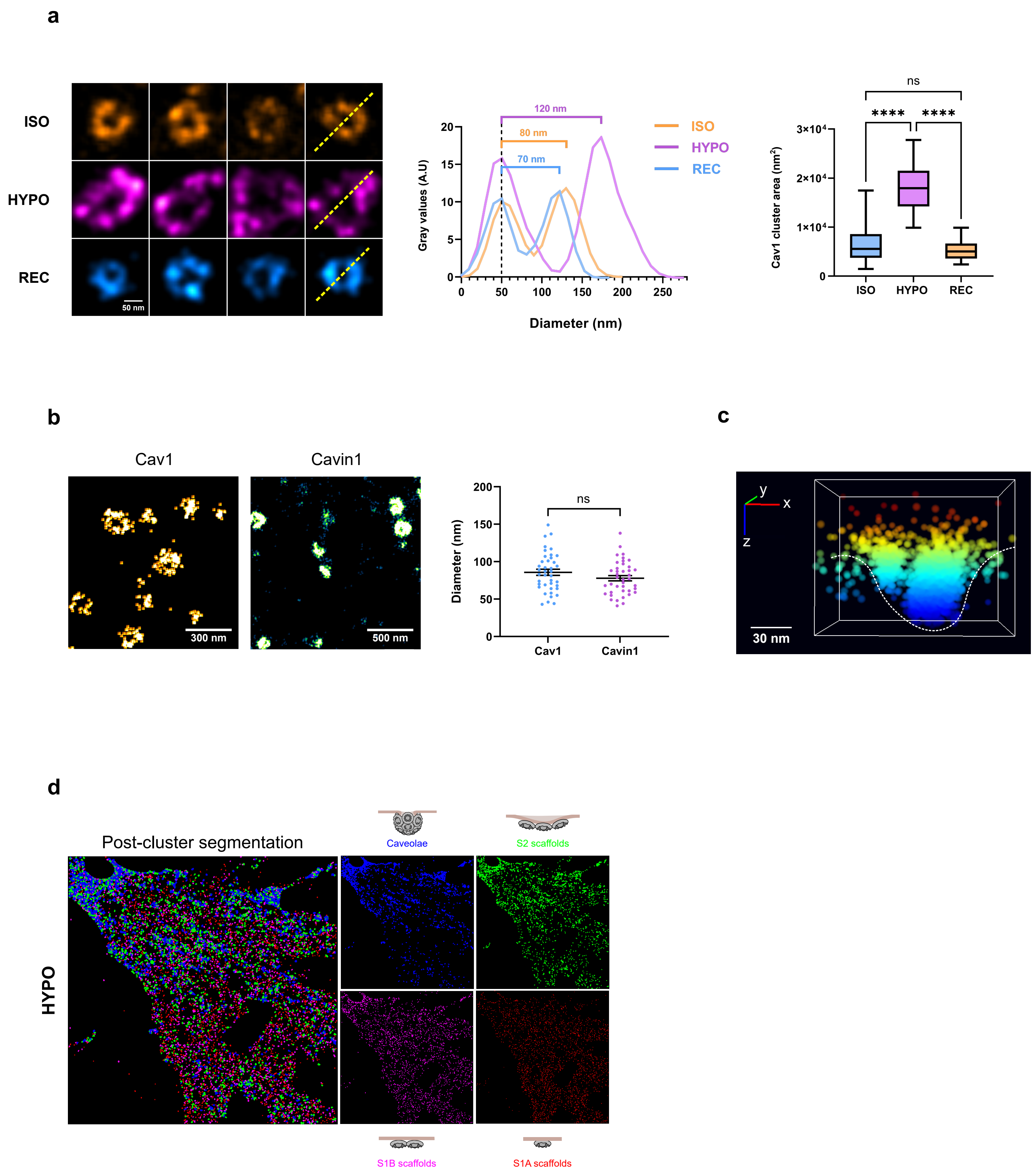

Extended Data figure 4:

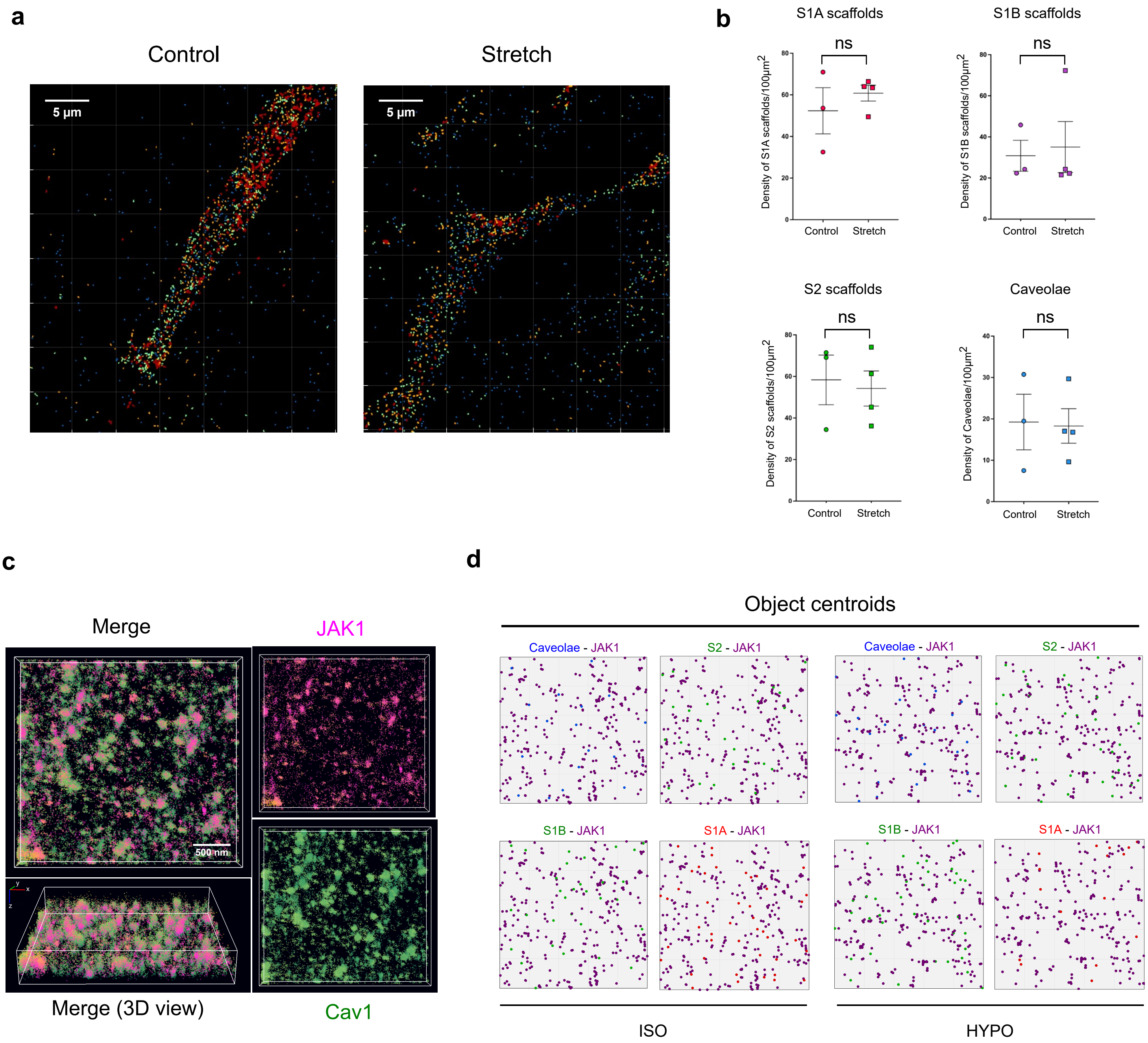

Extended Data figure 5:

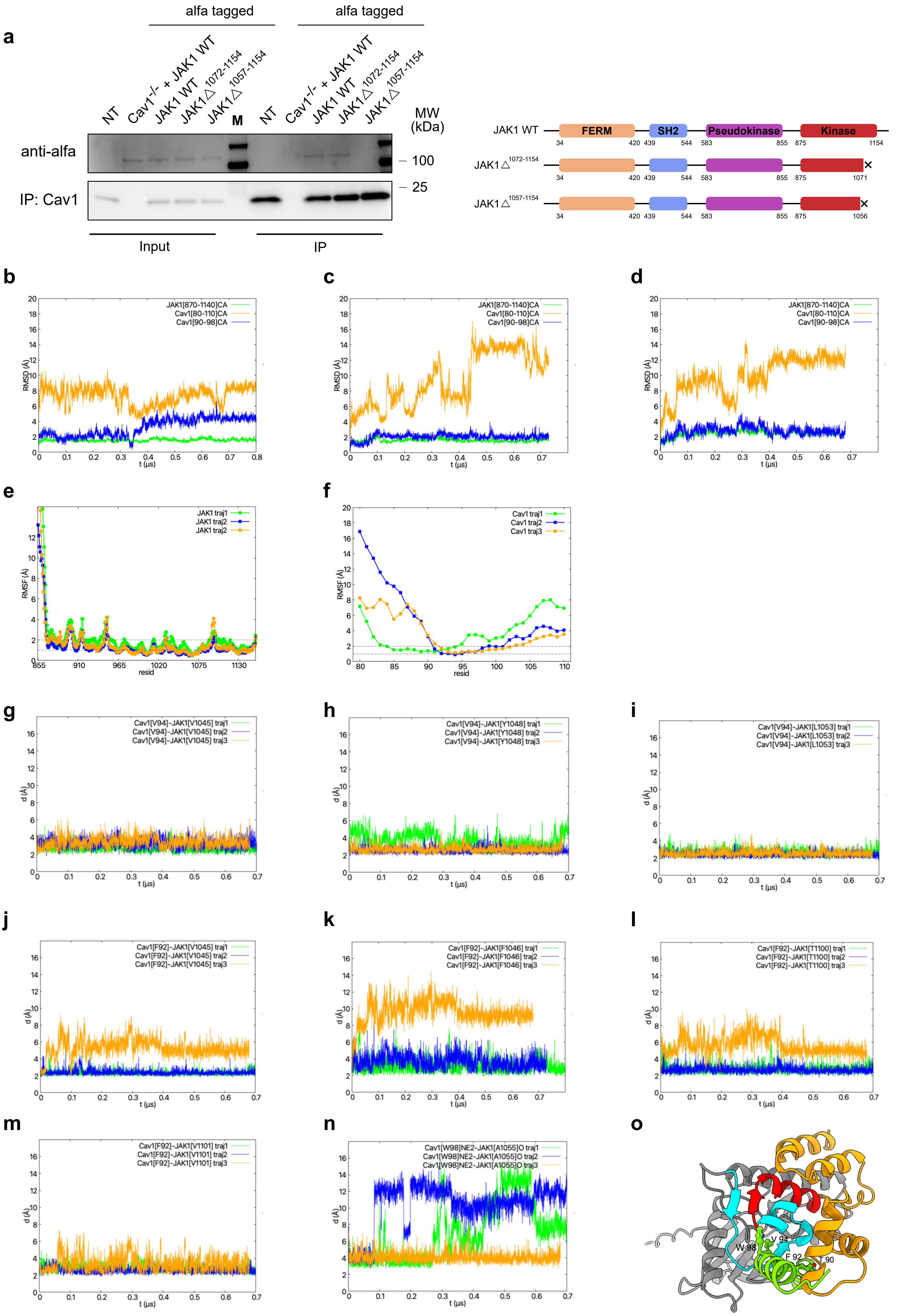

Extended Data figure 6:

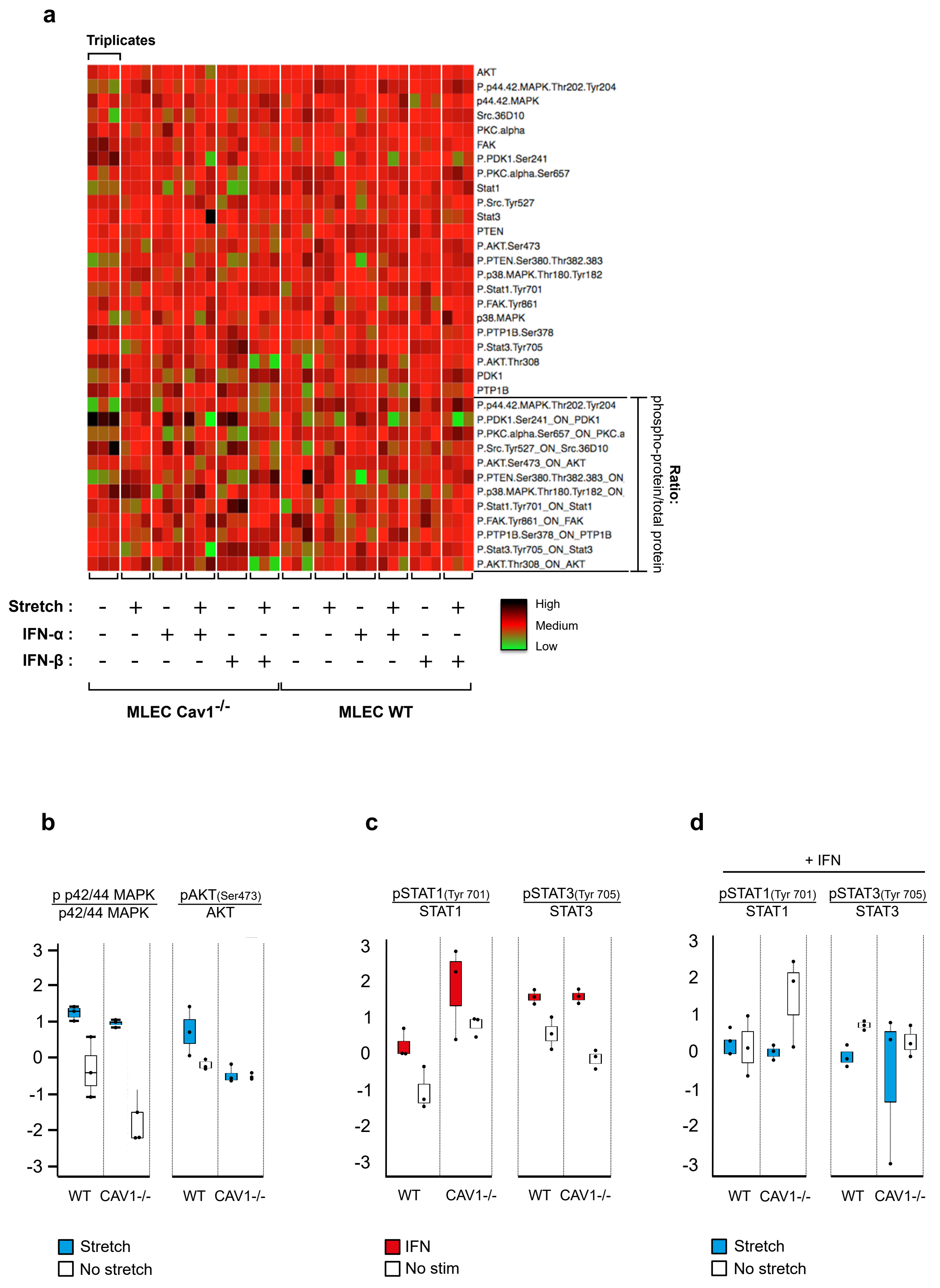

Extended Data figure 7:

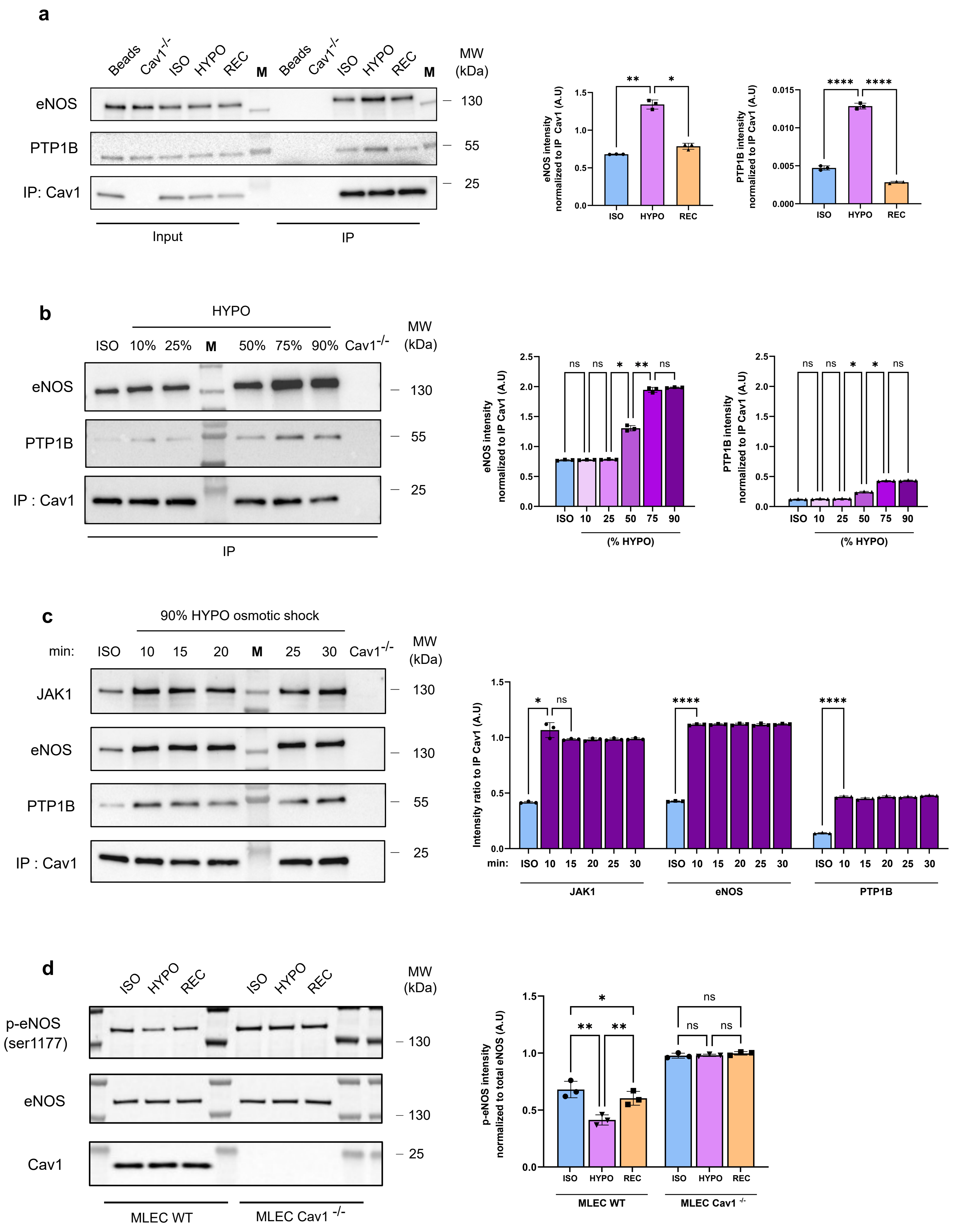

Extended Data figure 8:

a

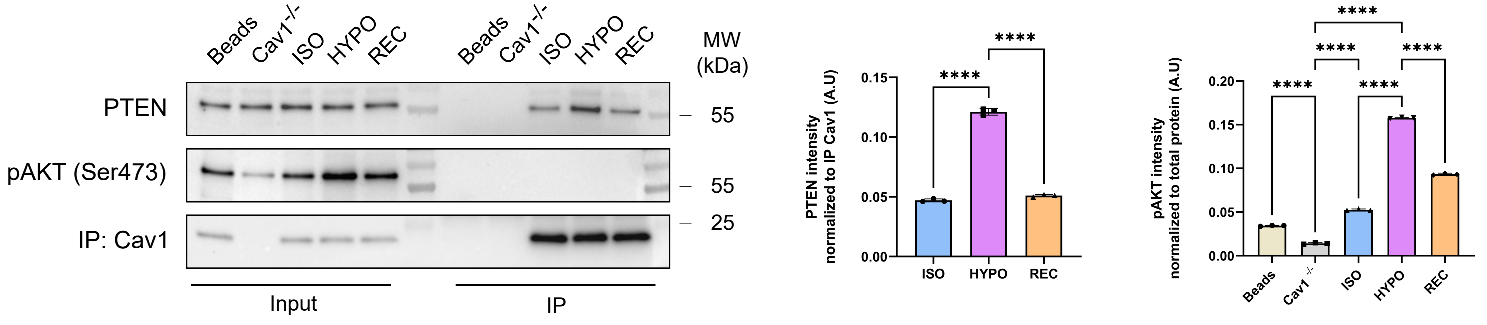


b

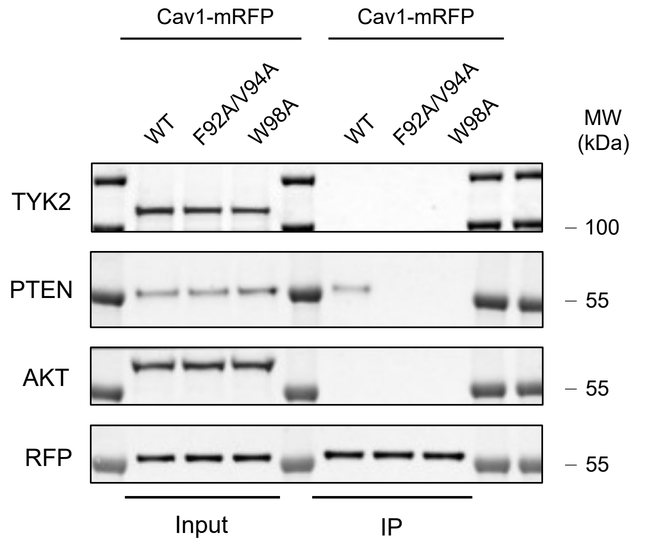


c

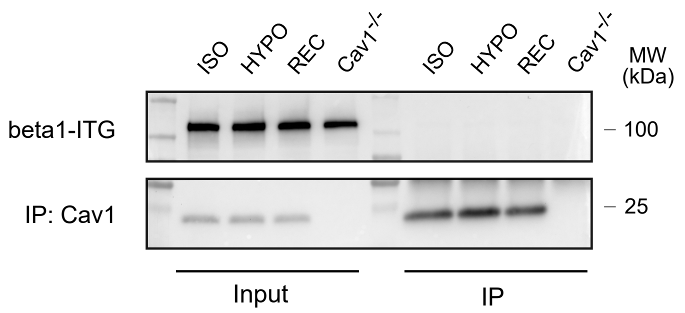
