## Extended Data Tables for "Remote Control of Cell Signaling through Caveolae Mechanics"

### Extended data table 1:

| Cluster type | ISO* (mean number of Cav1 molecules per unit area) | HYPO* (mean number of Cav1 molecules per unit area) | ISO (proportion of Cav1 molecules) | HYPO (proportion of Cav1 molecules) |
|---|---|---|---|---|
| Caveolae | 18,878.4 | 12,498.88 | 48.87% | 35.01% |
| S2 scaffolds | 11,728.8 | 13,046.4 | 30.36% | 36.5% |
| S1B scaffolds | 5365.8 | 6971.8 | 13.89% | 19.5% |
| S1A scaffolds | 2652.1 | 3180.1 | 6.86% | 8.90% |
| Total | 38,625.1 | 35,693.18 | | |

* Calculated on the basis that:
  - Caveolae is composed of 144 Cav1 molecules
  - S2 is composed of 72 Cav1 molecules
  - S1B is composed of 22 Cav1 molecules
  - S1A is composed of 11 Cav1 molecules

### Extended data table 2:

| Cluster type | ISO (mean number of blobs per unit area) | HYPO (mean number of blobs per unit area) | % difference (as blobs per unit area) |
|---|---|---|---|
| Caveolae | 131.1 | 86.77 | - 33.81% |
| S2 scaffolds | 162.9 | 181.2 | + 11.23% |
| S1B scaffolds | 243.9 | 316.9 | + 29.9% |
| S1A scaffolds | 241.1 | 289.1 | + 19.9% |
