## Supplementary_note_Theoretical_analysis_Kailasam_et_al_2026 for "Remote Control of Cell Signaling through Caveolae Mechanics"

### Supplementary Note 1 - Theoretical Analysis

In this supplementary note, we develop a simplified model for the disassembly of caveolae under stress based on equilibrium thermodynamics to quantify the principle of remote signalling described in the main text. Caveolae are self-assembled invaginated membrane domains. Their formation can be understood as a phase separation process driven by intramolecular interactions between components of the caveolar membrane domain and their affinity for a particular membrane curvature [1, 2, 3]. Membrane tension increases the cost of membrane deformation, such that curved domains are stable only below a membrane tension threshold that depends on the concentration of Cav1 at the membrane [1]. As the tension increases, domains disassemble and release their content, such that the concentration of freely diffusing Cav1 and their interaction with signalling effectors at the plasma membrane increase. The goal of the model, presented in Fig.6 of the main text is to describe this process in a semi-quantitative way.

In order to simplify the analysis, the model uses the following assumptions:

- The oligomerisation process can be described within the framework of equilibrium thermodynamics, which assumes reversible interactions. This is supported by the observation that the level of Cav1 interaction with effectors return to basal level upon recovery after an hypo-osmotic shock. However, the model does not include the temporal dependence of the disassembly process or the possibility that some structures are kinetically trapped *e.g.* due to their interaction with the cytoskeleton.

- To simplify the thermodynamic description, only five possible states are included, as shown in Fig.6A-B of the main text: full caveolae (considered to be full spheres), partial caveolae (so-called S2 scaffolds, considered to be half-spheres), freely diffusing Cav1 oligomers (S1A and S1B scaffolds) not bound to a signalling effector, freely diffusing individual effectors (active) and freely diffusing complexes of Cav1 and signalling effectors (inactive). A more complete model could allow a more progressive disassembly of caveolae and the existence of invaginated domains different from half-sphere [4]. The stoichiometry of the S1-effector complex can be either fixed to a 1:1 ratio, or allowed to vary up to a maximum number of effector per complex.

#### S1 Model

##### S1.1 fixed 1:1 stoichiometry

The starting point of the model is a free energy per unit surface that includes the energetics and entropy of the different states:

$$\begin{aligned}\mathcal{F}/k_{\mathrm{B}}T = {} & \rho_j \log[\rho_j/e] + \rho_1 \log[\rho_1/e] + \rho_d(\log[\rho_d/e] - \epsilon_d) \\ & +\rho_f(\log[\rho_f/e] + E_f) + \rho_h(\log[\rho_h/e] + E_h) \\ & -\mu_j(\rho_j + \rho_d) - \mu_c(\rho_1 + N\rho_f + N/2\rho_h)\end{aligned} \quad \text{(S1)}$$

where $e$ is the Euler's number. Here, $\rho_x$ is the concentration of the state $x$, with $x = j$ correspond to the active signalling effectors (not bound to Cav1), $x = 1$ to the freely diffusing Cav1 oligomers (S1A and S1B), $x = d$ to the Cav1-effector complexes associated to a binding energy $\epsilon_d$, $x = f$ to the full spheres (caveolae) - containing $N$ Cav1 oligomers and associated to an energy $E_f$ and $x = h$ to the half spheres (S2) - associated to an energy $E_h$. The conservation of Cav1 and effectors is enforced by introducing the chemical potentials $\mu_c$ and $\mu_j$, which are Lagrange multipliers associated to the total concentration of Cav1 ($\rho_{1,tot}$) and of effectors ($\rho_{j,tot}$), respectively. Energies are expressed in units of the thermal energy $k_{\mathrm{B}}T$ ($k_B$ is the Boltzmann constant).

Minimisation of the free energy with respect to the different concentrations ($\partial\mathcal{F}/\partial\rho_x = 0$) yields the concentration of the different states:

$$\begin{aligned}
&\rho_j = e^{\mu_j},\ \rho_d = e^{\mu_j+\mu_c+\epsilon_d} \quad \text{with} \quad e^{\mu_j}\left(1+e^{\epsilon_d+\mu_c}\right) = \rho_{j,tot} = e^{\mu_{j,tot}} \\
&\rho_1 = e^{\mu_c},\ \rho_f = e^{N\mu_c-E_f},\ \rho_h = e^{N/2\mu_c-E_h} \\
&\text{with} \quad e^{\mu_c}\left(1+e^{\mu_j+\epsilon_d}\right) + Ne^{N\mu_c-E_f} + \frac{N}{2}e^{N/2\mu_c-E_h} = \rho_{1,tot} = e^{\mu_{c,tot}}
\end{aligned} \tag{S2}$$

where $\mu_{j,tot}$ and $\mu_{c,tot}$ are the maximum possible values for the chemical potential of effectors and Cav1, respectively, in the absence of complexation ($\rho_j = \rho_{j,tot}$ and $\rho_1 = \rho_{1,tot}$)

##### S1.2 Variable stoichiometry

Considering that diffusing Cav1 oligomers contain several CAV1, one could envision that each oligomer can bind several signaling effectors. The above model is easily be modified to include a maximum stoichiometry of one S1 oligomer for an arbitrary number of $n_j$ effector. The free energy becomes

$$\begin{aligned}
\mathcal{F}/k_{\mathrm{B}}T = \rho_j \log[\rho_j/e] + \rho_1 \log[\rho_1/e] + \sum_{n=1}^{n_j} \rho_{d,n}(\log[\rho_{d,n}/e] - n\epsilon_d) \\
+\rho_f(\log[\rho_f/e] + E_f) + \rho_h(\log[\rho_h/e] + E_h) \\
-\mu_j(\rho_j + n\rho_d) - \mu_c(\rho_1 + N\rho_f + N/2\rho_h)
\end{aligned} \tag{S3}$$

where $\rho_{d,n}$ is the density of complexes containing one Cav1 oligomer and $n$ effectors, and $\epsilon_d$ is (as before) the binding energy per effector. Minimisation of the free energy with respect to the different concentrations yields an extension of Eq. S2 for the concentration of the different states:

$$\begin{aligned}
&\rho_j = e^{\mu_j},\ \rho_{d,n} = e^{n(\mu_j+\epsilon_d)+\mu_c} \quad \text{with} \quad e^{\mu_j} + \sum_{n=1}^{n_j} ne^{n(\epsilon_d+\mu_j)+\mu_c} = \rho_{j,tot} = e^{\mu_{j,tot}} \\
&\rho_1 = e^{\mu_c},\ \rho_f = e^{N\mu_c-E_f},\ \rho_h = e^{N/2\mu_c-E_h} \\
&\text{with} \quad e^{\mu_c}\left(1+\sum_{n=1}^{n_j} e^{n(\mu_j+\epsilon_d)}\right) + Ne^{N\mu_c-E_f} + \frac{N}{2}e^{N/2\mu_c-E_h} = \rho_{1,tot} = e^{\mu_{c,tot}}
\end{aligned} \tag{S4}$$

##### S1.3 Model of caveola mechanics

Membrane tension affect the energy of caveolae and S2 complexes [1, 5]. In a first approximation, we may assume that the domains are sufficiently rigid so that tension does not modify their curvature. In this case, the energy of the full and half spheres can be written

$$E_f = -N\epsilon_b + N\sigma s \quad \text{and} \quad E_h = -\frac{N}{2}\epsilon_b + \frac{N}{4}\sigma s + \lambda l\sqrt{N} \tag{S5}$$

where $\sigma$ is the membrane tension, $\epsilon_b$ is the binding energy between oligomers within a caveolae and $\lambda l$ represents the cost associated to the domain's line tension $\lambda$, which exist for half-spheres [5]. $s$ is the area occupied by a Cav1 oligomer in caveolae and $l = \sqrt{\pi s}$ is the length scale associated to the line tension energy. The factor 1/4 in front of the tension term in the energy of half spheres is due to the fact that this term is proportional to the difference between the domain area ($\sim N/2$) and its projection on the membrane ($\sim N/4$) [5].

#### S2 Results

Eqs.(S2,S5) entirely define the equilibrium distribution of the different states and how it varies with membrane tension. Expected orders of magnitude for the different parameters are as follows. The different binding energies $\epsilon_d$ and $\epsilon_b$ are expected to be of order a few $k_{\rm B}T$. Caveolae have a typical diameter of order 50 nm and contains $N \simeq 13$ oligomers, so that $s \simeq 500\,\mathrm{nm}^2$ and $l \simeq 40\,\mathrm{nm}$. The typical value of cell membrane tension varies (widely) between $10^{-6}$ N/m and $10^{-4}$ N/m [6] and the thermal membrane tension scale is $k_{\rm B}T/s \simeq 10^{-5}$ N/m, so one may expect $\sigma s$ (in $k_{\rm B}T$ unit) to vary between 0.1 and 10. The typical line tension of membrane domain is within the range $0.1 - 1$ pN [7] and the thermal line tension scale is $k_{\rm B}T/l \simeq 0.1$ pN so we expect $\lambda l$ (in $k_{\rm B}T$ unit) to be in the range $1 - 10$.

We first analyse the results of Eqs.(S2,S5) for the formation of caveolae disregarding Cav1 interaction with effectors, by setting $\mu_{j,tot}$ to a very low value. The results are shown in Fig. S1. As discussed above, caveolae can only form under low tension and most Cav1 are in freely diffusing oligomers (S1A/S1B) for large tension. Half-spheres (S2) are present only at intermediate tension, and there existence in conditioned to having a relatively low line tension (examples for high and low line tension are shown in Fig. S1). The level of tension at which the transitions occurs and the prominence of S2 structures depend on the total concentration of Cav1 at the membrane (examples for high and low concentration are shown in Fig. S1).

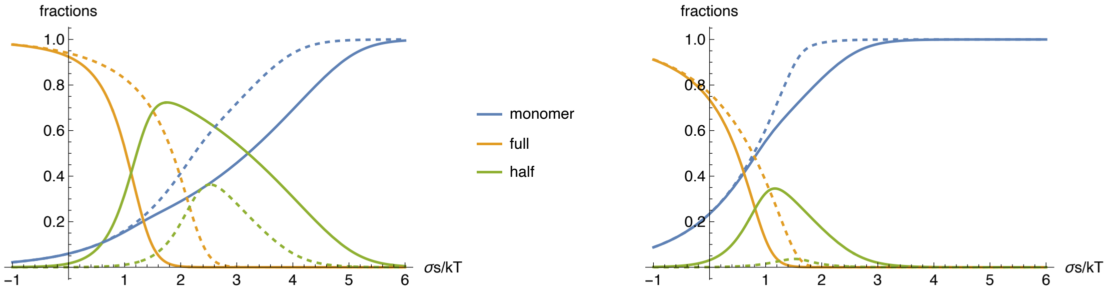


Figure S1: Fraction of the total Cav1 population in the three different states: monomers (S1A/S1B), half spheres (S2) and full spheres (caveolae) in the absence of interaction with effectors ($\mu_{j,tot} \to -\infty$). The solid curves are for low line tension ($\lambda l k_{\rm B}T = 1.5$) and the dashed curves for higher line tension ($\lambda l k_{\rm B}T = 2.5$). The left panel in for high Cav1 concentration ($\mu_{c,tot} = -1.5 k_{\rm B}T$) and the right panel for a low concentration ($\mu_{c,tot} = -3 k_{\rm B}T$). Other parameters are: $e_b = 4k_{\rm B}T,\ N = 13$.

Next we analyse the effect of caveolae disruption on the fraction of active effectors at the membrane for a finite value of $\mu_{j,tot}$. The results for a stoichiometry 1:1 are shown in Fig. S2. Effectors are progressively inactivated as tension increases, even though caveolae and half-spheres are still present at the membrane. Under high tension, the ratio of inactive to active effector is directly proportional to the Cav1 concentration (examples for high and low concentration are shown in Fig. S2).

Similar results for a maximum stoichiometry of 1:$n_j$ are shown in Fig. S3.

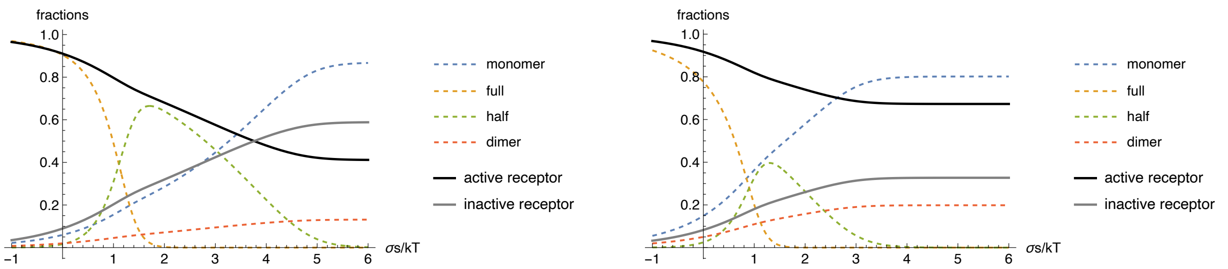


Figure S2: Fraction of the membrane effector in the active state (not bound to a Cav1 oligomer) and in the inactive state, with $\mu_{j,tot} = -3k_{\mathrm{B}}T$ and $\epsilon_d = 2k_{\mathrm{B}}T$. The different populations of Cav1 complexes are shown as dashed lines. Other parameters are as in Fig. S1, with ($\lambda l k_{\mathrm{B}} T = 1.5$). The left panel in for high Cav1 concentration and the right panel for a low concentration.

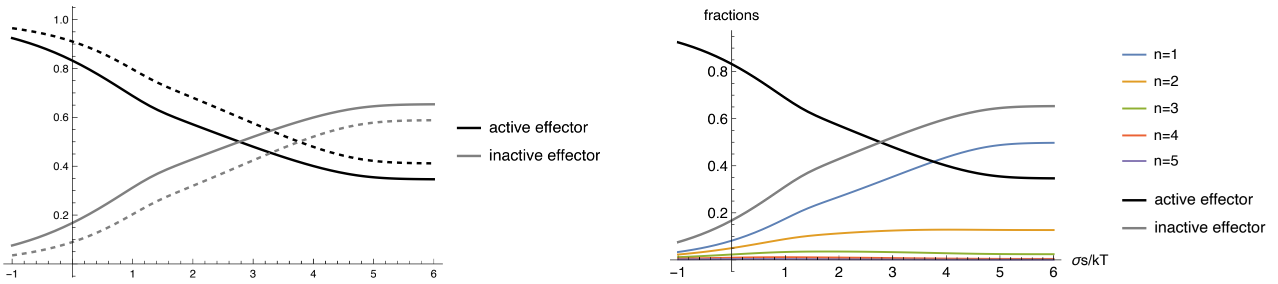


Figure S3: Fraction of the membrane effector in the active and inactive states when multiple effectors can bind to a single Cav1 oligomer. Parameters are as in Fig. S2 (left panel, high Cav1 concentration, $\mu_{c,tot} = -1.5k_{\mathrm{B}}T$). The left panel compares with the results for $n_j = 1$ (as in Fig. S2) in dashed black and gray lines and for $n_j = 5$. The right panel shows the fraction of Cav1 oligomers with different number of effector bounds (from $n = 1$ to $n = n_j = 5$).

### References

[1] P. Sens and M. S. Turner, “Theoretical model for the formation of caveolae and similar membrane invaginations,” *Biophys. J.*, vol. 86, pp. 2049–2057, 2004.

[2] B. J. Reynwar, G. Illya, V. A. Harmandaris, M. M. Müller, K. Kremer, and M. Deserno, “Aggregation and vesiculation of membrane proteins by curvature-mediated interactions,” *Nature*, vol. 447, no. 7143, pp. 461–464, 2007.

[3] F.-N. Lolo, N. Walani, E. Seemann, D. Zalvidea, D. M. Pavón, G. Cojoc, M. Zamai, C. Viaris de Lesegno, F. Martínez de Benito, M. Sánchez-Álvarez, J. J. Uriarte, A. Echarri, D. Jiménez-Carretero, J.-C. Escolano, S. A. Sánchez, V. R. Caiolfa, D. Navajas, X. Trepat, J. Guck, C. Lamaze, P. Roca-Cusachs, M. M. Kessels, B. Qualmann, M. Arroyo, and M. A. del Pozo, “Caveolin-1 dolines form a distinct and rapid caveolae-independent mechanoadaptation system,” *Nature Cell Biology*, 2022.

[4] N. Sarkar, C. Lamaze, and P. Sens, “Mechano-sensitivity of multi-component caveolae,” *bioRxiv*, 2023.

[5] P. Sens and M. Turner, “Budded membrane microdomains as tension regulators,” *Phys. Rev. E*, vol. 73, p. 031918, 2006.

[6] P. Sens and J. Plastino, “Membrane tension and cytoskeleton organization in cell motility,” *J. Phys. Condens. Matter*, vol. 27, p. 273103, 2015.

[7] T. Baumgart, S. T. Hess, and W. W. Webb, “Imaging coexisting fluid domains in biomembrane models coupling curvature and line tension,” *Nature*, vol. 425, pp. 821–824, 2003.
